# Reinterpreting Keratoconus Gene Expression in Light of Inherited Stromal Weakness

**DOI:** 10.64898/2026.10.03.754609

**Authors:** Hanyu Jiang, Fei Gao, Wei Zhang, Ying Jie, Ying Li, Yang Jiang

## Abstract

**Purpose:** To reinterpret keratoconus (KC) gene expression in light of inherited stromal weakness, and to define how expression should guide model and target selection.

**Methods:** We combined genetic and chromatin data with expression from eight studies (182 donors). We compared the same KC samples with normal and pathological controls, built references matched to control source and tissue layer and tested them in independent tissues, and indexed published intervention-outcome records. Stimulated cells and genetic analyses assessed repair capacity and inherited risk.

**Results:** Changing only the control reversed expression directions, flipping KC-like classification in 21–27 of 42 models and supplement-or-inhibit direction in 361–385 of 1,090 targets. Our references addressed this, predicting direction more accurately than published methods in independent spatial stroma and two bulk comparisons. Against these references, 41 of 46 target nominations required a different comparison, measurement, or use. A correct direction described only the stromal state: across control types, KC stroma showed lower inflammatory, immediate-early (stress-response), and transforming growth factor-β (TGF-β) programs, which signaled neither lost repair capacity (stimulated KC cells increased matrix production) nor disease cause (no genetic enrichment). Inherited risk converged on the matrix: collagen assembly was the only replicated pathway family, and genetically predicted higher corneal resistance lowered disease odds independently of thickness. Because these expression changes do not drive disease, reversing them does not ensure functional benefit: among 2,769 intervention-outcome records, the two were separate outcomes.

**Conclusions:** Expression direction therefore reflects the choice of control and the stromal state, and does not indicate how to intervene. Because inherited risk lies in matrix structure and resistance to deformation, repair should restore organized, load-bearing matrix rather than reverse expression changes. Models should therefore be judged against matched references, and targets by measured functional outcomes.

## Introduction

Keratoconus (KC) progressively distorts the cornea as its stroma loses resistance to deformation, affects an estimated 23.7 million people worldwide, and is usually diagnosed in young adults.^1,2^ Genetic studies implicate collagen assembly and corneal biomechanics,^3−6^ yet tissue expression profiles do not agree with one another. One transcriptome found particularly low expression of partners of the transcription factor activator protein-1 (AP-1),^7^ whereas another found higher extracellular matrix (ECM) pathways;^8^ cone-apex epithelium shows higher inflammatory and lower matrix gene expression,^9^ while another cohort shows lower acute-phase and tissue-injury responses but higher matrix transcripts;^10^ and a further study reports enrichment of immune and inflammatory genes.^11^ Matrix and inflammatory programs are thus reported as both higher and lower in KC.

These directions are being used to choose treatments. Higher inflammatory expression has prompted inhibition of AP-1 and PTGS2 (cyclooxygenase-2),^11−13^ and an eye drop acting through AP-1 signaling has been reported to prevent progression in a rabbit KC model,^13^ although an earlier transcriptome found AP-1 partners particularly downregulated.^7^ Lower matrix expression^14^ has prompted replacement-cargo nomination.^15^ Yet the profiles differ in layer and in control, from corneas of eyes enucleated for melanoma^7^ and pathological grafts^14^ to epithelium of refractive-surgery patients,^9^ and a direction exists only relative to its control: a transcript lower than in a pathological graft need not be lower than in normal cornea. A single control source within each study can conceal this dependence during data integration. Every decision drawn from expression direction therefore inherits the choice of control.

Even recurring findings, such as low immediate-early (stress-response) expression,^7,8,10^ leave their meaning open. They could reflect a cause of disease, a response to it, or lost repair capacity, and baseline expression, stimulus responses, and inhibitory feedback on transforming growth factor-β (TGF-β) signaling measure different aspects of repair. Which applies decides whether reversing an expression change would help, and how these changes relate to the inherited weakness of the stroma remains unresolved.

Here we reinterpret KC gene expression in light of inherited stromal weakness. We held the same KC corneas fixed and changed only the control, and asked which stromal changes persist across controls and whether they mark inherited risk or lost repair capacity. We then built references matched to control source and tissue layer, tested them in independent tissues, and indexed published interventions with their measured outcomes. The direction of expression change depended on the control, and the changes that persisted reflected the diseased stroma rather than inherited risk, which centered on the matrix and its resistance to deformation. These findings show how gene-expression differences between KC and control corneas should be read, as the state of the diseased stroma, and how models and targets should be chosen, against matched references and measured outcomes.

## Methods

The core computational analysis used public, deidentified data, without new human sampling or animal use. Source studies obtained ethical approval and consent; the rabbit series used commercially supplied eyes. Control provenance and tissue definitions were taken from the source publications (Table 1; Supplementary Table S1; Supplementary Methods).

**Table 1.**
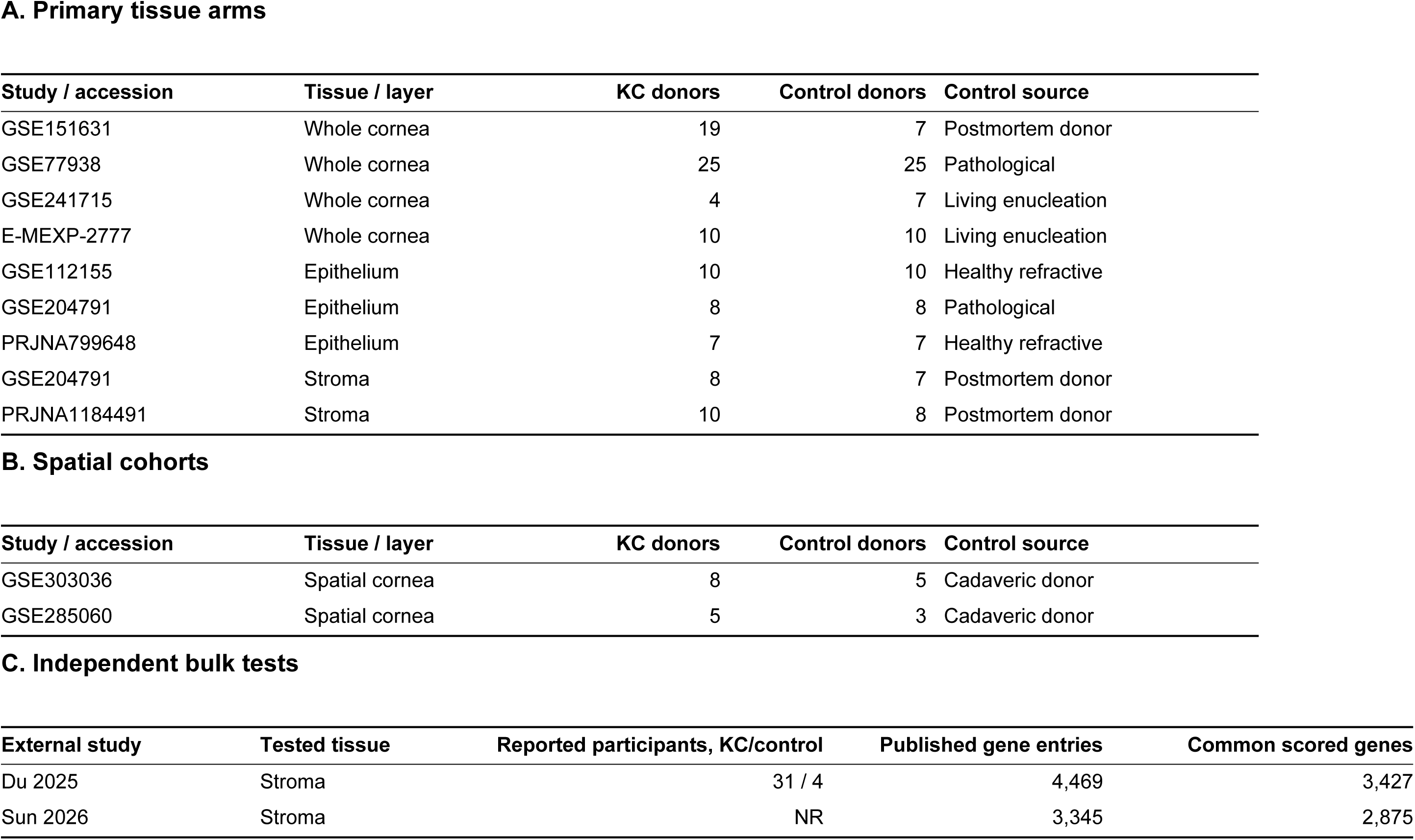
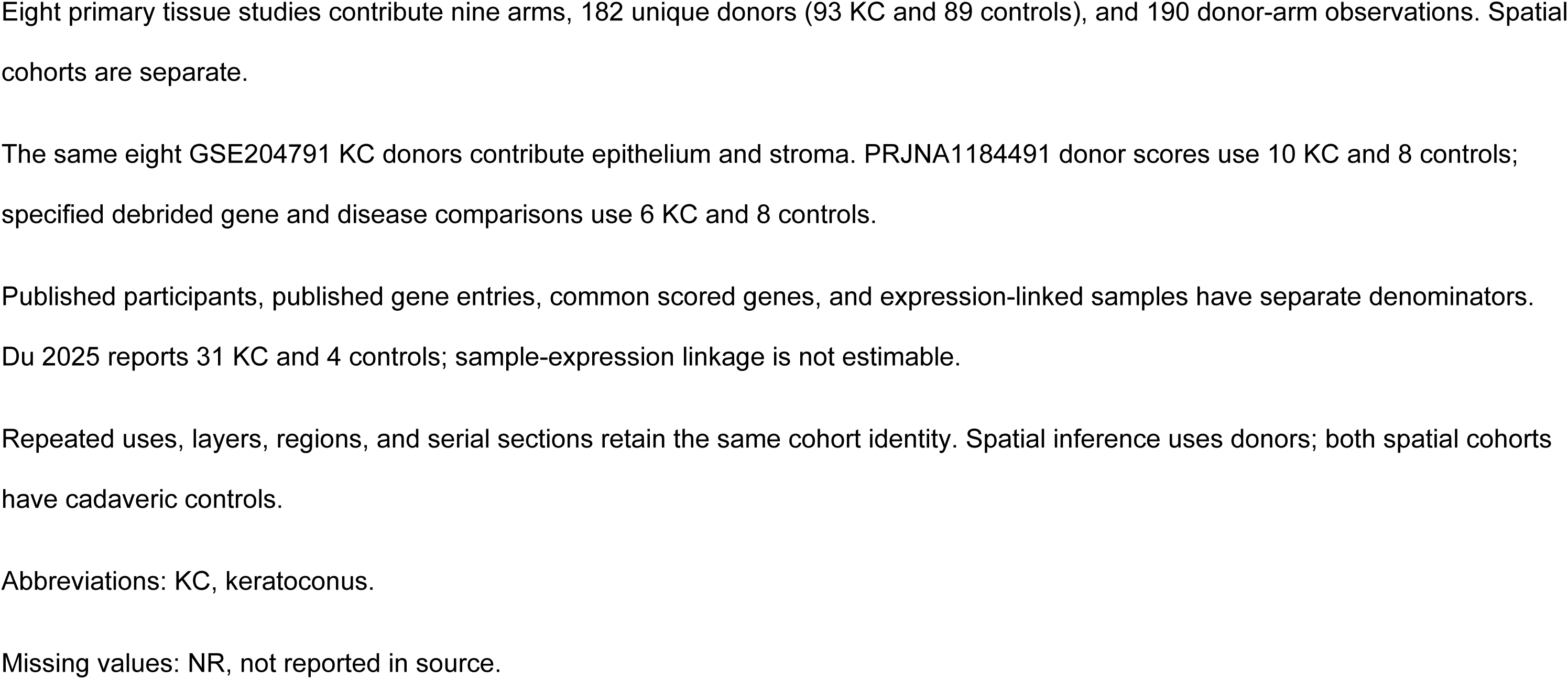
Human tissue cohorts and independent expression tests.

| **A. Primary tissue arms** |  |  |  |  |
|---|---|---|---|---|
| Study / accession | Tissue / layer | KC donors | Control donors | Control source |
| GSE151631 | Whole cornea | 19 | 7 | Postmortem donor |
| GSE77938 | Whole cornea | 25 | 25 | Pathological |
| GSE241715 | Whole cornea | 4 | 7 | Living enucleation |
| E-MEXP-2777 | Whole cornea | 10 | 10 | Living enucleation |
| GSE112155 | Epithelium | 10 | 10 | Healthy refractive |
| GSE204791 | Epithelium | 8 | 8 | Pathological |
| PRJNA799648 | Epithelium | 7 | 7 | Healthy refractive |
| GSE204791 | Stroma | 8 | 7 | Postmortem donor |
| PRJNA1184491 | Stroma | 10 | 8 | Postmortem donor |
| **B. Spatial cohorts** |  |  |  |  |
| **Study / accession** | **Tissue / layer** | **KC donors** | **Control donors** | **Control source** |
| GSE303036 | Spatial cornea | 8 | 5 | Cadaveric donor |
| GSE285060 | Spatial cornea | 5 | 3 | Cadaveric donor |
| **C. Independent bulk tests** |  |  |  |  |
| **External study** | **Tested tissue** | **Reported participants, KC/control** | **Published gene entries** | **Common scored genes** |
| Du 2025 | Stroma | 31 / 4 | 4,469 | 3,427 |
| Sun 2026 | Stroma | NR | 3,345 | 2,875 |

Eight primary tissue studies contribute nine arms, 182 unique donors (93 KC and 89 controls), and 190 donor-arm observations. Spatial cohorts are separate.
The same eight GSE204791 KC donors contribute epithelium and stroma. PRJNA1184491 donor scores use 10 KC and 8 controls; specified debrided gene and disease comparisons use 6 KC and 8 controls.
Published participants, published gene entries, common scored genes, and expression-linked samples have separate denominators. Du 2025 reports 31 KC and 4 controls; sample-expression linkage is not estimable.
Repeated uses, layers, regions, and serial sections retain the same cohort identity. Spatial inference uses donors; both spatial cohorts have cadaveric controls.
Abbreviations: KC, keratoconus.
Missing values: NR, not reported in source.

### Datasets

The three independent disease sources were two KC genome-wide association studies (GWAS), Hardcastle (GCST90013442)^3^ and Million Veteran Program (MVP) European (GCST90477661),^16^ and the FinnGen R13 corneal-deformation endpoint (H7_CORNEALDEFORM).^17^ The overlapping MVP mixed-ancestry analysis was used for reference-panel comparisons. Mendelian randomization (MR) and the original structural-module analyses used central corneal thickness (CCT; GCST006366, 17,803 participants)^18^ and corneal resistance factor (CRF; GCST90100568, 76,029 participants).^19^ CRF was measured in mm Hg using the average of both eyes. Expanded colocalization used the larger CCT meta-analysis (GCST90000654, 44,039 participants),^20^ alongside the original CCT input. Genetically CCT-residualized CRF (CRF⊥CCT) came from genomic structural equation modeling.^21^

Nine tissue arms came from eight studies: GSE151631,^10^ GSE77938,^14^ GSE112155,^22^ GSE241715,^23^ GSE204791,^24^ E-MEXP-2777,^7^ PRJNA1184491,^25^ and PRJNA799648.^11^ Controls were postmortem donor, healthy/refractive, pathological, or living-enucleation tissue. The donor analysis included 93 KC and 89 control individuals. Eight KC donors contributed both epithelial and stromal arms, giving 190 donor–arm observations. PRJNA1184491 contributed 10 KC and eight control donors after averaging central and peripheral expression within donor; gene-effect synthesis and disease comparisons used its six-KC/eight-control stromal subset.

Disease comparisons included bacterial/fungal and viral keratitis (GSE58291^26^ and GSE241715) and Fuchs endothelial corneal dystrophy (FECD; GSE142538^27^ and GSE171830^28^). Spatial cohorts were GSE303036^8^ and GSE285060.^29^ Independent bulk tests used the published expression tables of Du et al.^30^ (31 KC stromal specimens from deep anterior lamellar keratoplasty (DALK) and four eye-bank controls) and Sun et al.^31^ (KC versus stroma from small-incision lenticule extraction (SMILE)). Other inputs included human and rabbit stromal-cell perturbations (GSE260476^32,33^ and GSE287915^34^), reduced-representation bisulfite sequencing (RRBS; GSE107189),^35^ and N^6^-methyladenosine (m6A; GSE181618). Clinical, regional, blood, and culture inputs are detailed in Supplementary Methods.

### Gene Sets and Genetic Enrichment

Six modules were fixed before testing: extracellular matrix (ECM)/proteoglycan, stress/immediate-early, general inflammation, corneal-local inflammation, oxidative stress/NRF2, and literature-derived atopic/systemic genes. TGF-β signaling and canonical type-2 atopy were additional tissue sets (Supplementary Table S2). Alias resolution^36^ and a common gene universe retained MHC-region genes and genes analyzable across sources and linkage disequilibrium (LD) panels. MAGMA v1.10^37^ used 1000 Genomes LD^38^ for competitive enrichment.^39^ Marginal tests fitted modules separately; joint tests fitted all six. Source-specific thresholds were 0.05/6 for modules and 0.05/5 for attribution tests. Structural-trait, coefficient-contrast, matched-set, and deletion analyses are detailed in Supplementary Methods and Supplementary Table S3.

The public-pathway scan used MSigDB 2026.1.Hs Reactome, GO Biological Process, KEGG MEDICUS, and ten canonical NABA sets.^40^ Sets with 10–500 complete Entrez members were eligible; an extension allowed 10–1,000. Each set was intersected with the source-specific MAGMA universe. Discovery required positive enrichment with Benjamini–Hochberg^41^ *q* < 0.05 in Hardcastle; replication required positive enrichment with *P* < 0.05 in MVP European or FinnGen. Overlapping terms were summarized as families. Tissue effects used the unshrunk gene-effect model below (Supplementary Table S3).

### Mendelian Randomization

Forward MR estimated disease odds per 1-mm Hg higher CRF or 1-μm higher CCT; reverse MR estimated trait change per unit increase in genetic log odds of disease. Analyses used inverse-variance weighting (IVW), MR-Egger, weighted median, weighted mode, and MR-PRESSO.^42−46^ We assessed heterogeneity, the Egger intercept, instrument strength, and Steiger directionality, and repeated forward IVW after excluding instruments within 500 kb of genome-wide significant disease variants.

CAUSE compared causal and shared-pleiotropy models in both directions for each source.^47^ Multivariable MR jointly fitted CRF and CCT using their combined instruments, reclumped at *P* < 5 × 10^−8^, r^2^ = 0.001, and 10,000 kb.^48^ The model used multiplicative random-effects IVW without an intercept and normal inference.^49^ Instruments and diagnostics are in Supplementary Methods and Supplementary Table S4.

### Colocalization Across KC Risk Regions

Genome-wide significant disease variants were grouped using European LD, a 500-kb radius, and lead-variant r^2^ < 0.1. Overlapping windows were merged across sources into 40 physical regions. SuSiE^50,51^ and coloc^52,53^ used p_1_ = p_2_ = 10^−4^ and p_12_ = 5 × 10^−6^; coloc.abf supplied posteriors when no usable SuSiE pair was available. Sharing required PP4 ≥ 0.8 for any valid source–trait pair. Primary channels were updated CCT, CRF, and CRF⊥CCT; original CCT was a comparison channel. An expanded set added 21 published lead-centered windows, analyzed against the same disease sources. Original and expanded windows and source-specific posteriors remained separate (Supplementary Table S5). At COL5A1, we resolved shared components to their credible variants, retained each trait’s original posterior inclusion probability (PIP), and checked exact coordinates against adult stromal peaks. Reporter amplicons from Mustafa’s thesis^54^ were mapped by primer pairs to GRCh38. Within each amplicon, we summed variant posteriors conditional on sharing and phased variants against the CCT-lowering lead in 1000 Genomes Europeans.

### Chromatin Accessibility, Heritability Share, and Expression Direction

Credible variants from the original 17 shared regions were overlapped with adult corneal open chromatin (four donors; GSE155683).^55^ PIP-weighted peak overlap was compared with 1,000 draws of 17 regions from the original 40. Canonical-correlation anchor transfer assigned 11,947 ATAC barcodes to 21 author-defined RNA classes in Signac.^56^ Logistic regression adjusted for peak counts identified class-enriched peaks. These annotations also supplied cell context for the expanded shared-window set; enrichment testing used the original 17-region set.

LD score regression^57^ estimated genetic correlations with corneal structural traits. Their joint projection onto disease common-SNP heritability was r_Kᵀ R_S^−1^ r_K, with 200-block jackknife intervals. Four prespecified trait sets retained one eye and zone per measurement family. Expression directions combined published risk-expression links^58^ with direct GTEx v8^59^ fibroblast/skin eQTL colocalization (PP4 ≥ 0.8), retaining each source’s criteria. These directions were compared with KC tissue effects by arm, layer, and control source.

### Gene-Level Tissue Synthesis and Control Provenance

Unshrunk KC−control log_2_ fold-changes (log_2_FC) and sampling variances were synthesized across nine arms using metafor multilevel random-effects models, with arm and gene-within-arm intercepts and restricted maximum likelihood (REML).^60^ We reported pooled estimates, 95% confidence intervals (CIs), two-sided Wald *P* values, arm means, and eight source-study omissions, removing both GSE204791 arms together (Supplementary Table S6).

Provenance models included control source and tissue layer, with predictions restricted to observed source–layer combinations. M1 treated living enucleation as a separate class. Alternative classifications M0, M2, and M3, likelihood-ratio tests, source-level permutations, and held-out-source predictions are detailed in Supplementary Methods and Supplementary Table S7.

### Donor and Spatial Analyses

Module scores were equal-weight means of within-arm standardized member expression. CollecTRI^61^ and DoRothEA ABC^62^ regulons with decoupleR’s univariate linear model^63^ estimated AP-1/immediate-early gene (IEG) activity. The six-factor composite contained ATF3, EGR1, FOS, FOSL1, JUN, and JUNB. Hedges’ *g*^64^ was pooled by REML overall and by layer (Supplementary Table S6).

Within study, SMAD7 and SMAD6 expression was regressed on diagnosis and donor-level SMAD3 target output, excluding feedback genes from the score and retaining study covariates. Diagnosis coefficients were pooled across independent studies by DerSimonian –Laird random effects. A further SMAD7 model added the stress/immediate-early module score, with genes shared between the two scores removed from the output score. Same-donor output–matrix models adjusted for diagnosis and SMAD7, removed the eight genes shared with matrix outcome scores from SMAD3 output, and standardized continuous variables within study. Primary arms, covariates, and paired-region analyses are detailed in Supplementary Methods.

Spatial modules were standardized across quality-filtered spots within cohort and averaged within donor. KC−control tests enumerated all 1,287 donor-label assignments in GSE303036 and all 56 in GSE285060; CIs came from donor-level linear models. Keratocyte-composition adjustment^65^ and dendritic-cell (DC)/immune-marker scores are described in Supplementary Methods and Supplementary Table S9.

### Disease Comparisons and Perturbations

Disease-versus-control module and AP-1 effects used Hedges’ *g*. KC−comparator differences included shared-control covariance from GSE241715, weighted by each study’s contribution. FECD repeat-expansion cases formed the primary GSE142538 comparison; other groups remained separate (Supplementary Table S10).

Serum and TGF-β1 responses used library-level ordinary least squares, HC3 standard errors,^66,67^ and exact label permutations. AP-1/IEG scores were sign-inverted so positive changes pointed toward the low KC state; ECM member scores used directions from four whole-cornea/stromal arms. Each score retained its scale (Supplementary Table S11). Published matched-stimulus comparisons in KC- and normal-derived cells^68−70^ were examined by ligand, cell source, and endpoint, distinguishing inhibitory regulators, phosphorylation, matrix output, and survival. Akoto et al.’s covariate-adjusted diagnosis effect^71^ was analyzed as a between-source expression contrast.

### Regional, Severity, and Blood Analyses

Paired central–peripheral expression was modeled with donor, region, and epithelial-marker terms; debrided stroma was analyzed separately. Clinical analyses related epithelial programs to severity, Kmax, and thickness, and compared atopic and nonatopic KC. Blood, RRBS, and m6A analyses and model specifications are provided in Supplementary Methods and Supplementary Table S10.^72−75^

### Mechanical Contrasts and Control-Source References

Forty-two contrasts included corneal and dermal fibroblast stretch, keratocyte time courses, skin expansion, curvature chips, and feeder layers; load-release and handling series were analyzed separately.^12,76−81^ Gene-effect vectors were compared with KC by inverse-variance-weighted cosine similarity, expression-matched gene permutations, and gene-resampling CIs. Duration interactions compared treatment effects against concurrent controls. The original pooled reference synthesized four whole-cornea/stromal arms.

For 29,100 genes measured in at least two tissue arms, M1 meta-regression fitted unshrunk log_2_FC to control source and layer, using sampling variances, cohort/gene random intercepts, and maximum likelihood. Five observed layer–control combinations defined the references. Selection used source and layer before external scoring. The postmortem whole-cornea/stromal reference matched both for stroma and whole sections and was applied by source match to epithelial, endothelial, and immune views. GSE303036 and Du’s published effects^30^ were excluded from fitting. Published comparators were GSE77938 alone, Niu’s ComBat–limma approach,^82^ MetaIntegrator,^83^ DerSimonian–Laird pooling,^84^ and Niu et al.’s released gene-level results.^82^ They were ranked separately from our reference variants.

GSE303036 counts were summed by donor and layer across serial sections before log_2_(CPM+1) transformation. Effects compared equally weighted KC and control donor means; genes were matched by stable Ensembl identifiers. Directional balanced accuracy averaged upward and downward sign accuracy; Spearman correlation assessed effect ranking. Paired bootstrap intervals used 2,000 within-diagnosis donor resamples, identical across methods, with predictions fixed. Bulk comparisons resampled published-table entries. Strong-effect subsets were selected by external absolute effect before intersecting coverage. A second reference combined predictions from six method families, with its configuration selected by balanced accuracy in whole-study holdouts and its predictions frozen before external evaluation. Downstream decisions used identical measured genes for each reference and the independent tissue effects. Implementations, reference definitions, and common-gene sets are detailed in Supplementary Methods.

### Fixed-Case Control Substitution

In GSE241715, the same four KC corneas were compared with seven author-defined normal corneas obtained at living enucleation, five viral, and five bacterial/fungal corneas using log_2_(author-normalized expression + 1). Model and cargo replays held measurable genes and normal-reference inverse-variance weights fixed. We decomposed the change into KC–normal and alternative-control–normal vectors. All 35 three/four splits of the normal donors tested recovery against held-out normal controls. The 42 models and all 1,253 published candidates remained fixed. Model classification used cosine sign; cargo direction followed the published supplementation/inhibition rule. Agreement was evaluated against the specified tissue comparison on common measurable genes. In GSE77938, the replay fixed all 25 KC corneas and varied pathological control class with common-gene weights fixed. The authors’ retained 20 controls^14^ were compared with all 25 and within their five pathological classes. Equations, resampling, and program attribution are detailed in Supplementary Methods.

### Patient-Derived Stromal-Cell Transcriptomes

We analyzed RNA-sequencing matrices from patient-derived bioprinted keratoconus stroma:^85^ three KC donors across two formats and four treatments, and a separate baseline comparison of three KC and three normal donors across both formats. The cultures used passage-3 isolates and received a four-hour, 10-ng/mL TGF-β pulse on day 7; RNA was collected 72 hours after pulse onset. DESeq2 models included donor and format–treatment group; interaction contrasts tested whether the stimulation response differed between formats. Frozen tissue modules and tissue-low gene sets were applied without redefining membership; conventional pathway enrichment and baseline disease-by-format models are specified in Supplementary Methods.

### Published Target Premises and Intervention Outcomes

Forty-six source–gene nominations from nine studies were compared with M1 estimates in the premise’s stated layer and measurement. A nomination was supported when its direction was significant against at least two control classes without a significant opposite estimate, changed use when only the opposite direction was significant, or required another measurement or comparison otherwise, including protein/phosphorylation premises. The published cargo pipeline^15^ was replayed by replacing only differential expression while retaining other evidence and weights. Interventions, systems, controls, readouts, and source coordinates formed 2,769 evidence records and 194 index entries. Each intervention record (an operation in Supplementary Table S14) specifies the perturbation and direction, system and exposure, comparator, and measured endpoint. Expression premises and functional outcomes are separate evidence types. Statements in Niu et al.^82^ and a citing review^86^ were checked against their full texts.

## Results

### Changing Only the Control Reverses Model and Candidate Decisions

We first tested whether changing only the control altered the expression profile used to select models and candidates. POSTN was higher in four KC samples than in normal cornea (log_2_ expression difference, +2.14) but lower than in viral or bacterial/fungal corneas (−4.63 and −2.99; Fig. 1A). With genes and weights held fixed, these substitutions reversed 27 and 21 of 42 model classifications (KC-like or opposite by cosine sign) and supplementation–inhibition directions (supplement if lower in KC, inhibit if higher) for 361 and 385 of 1,090 jointly measurable candidates in the published 1,253-candidate exosome-cargo pipeline (Fig. 1B,C). A classification reversed when the control-state projection had the same sign and greater magnitude than the KC–normal projection (Fig. 1D; Table 2). Because one set of cases cannot support both decisions, these calls belong to the comparison, not to KC.

**Figure 1.**
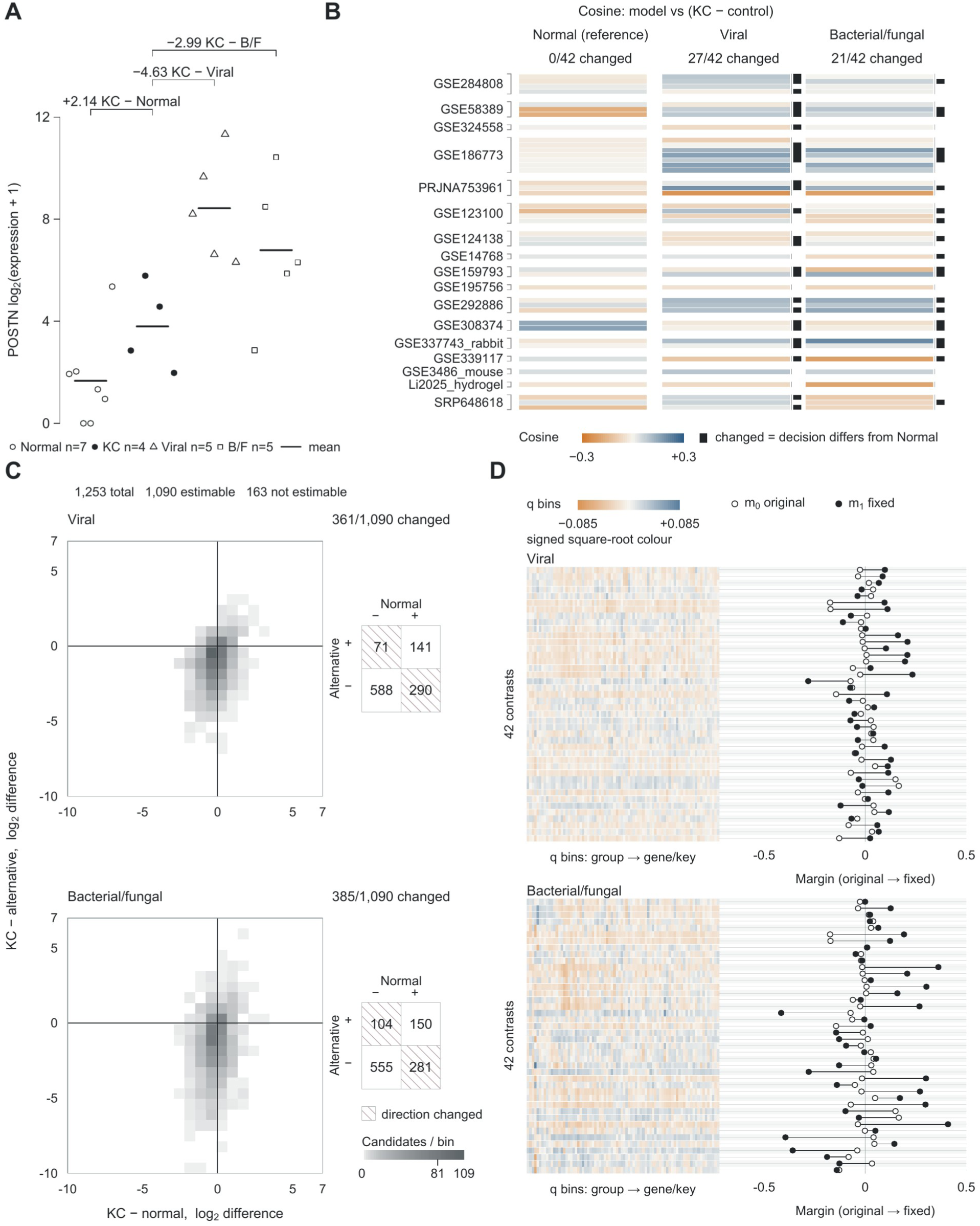
Changing controls with the same keratoconus samples reverses model and candidate decisions. **(A)** POSTN expression in GSE241715: 7 normal, 4 KC, 5 viral, and 5 bacterial/fungal corneas. Points represent samples and horizontal lines group means on the log_2_(normalized expression + 1) scale. The same KC samples give differences of +2.14, −4.63, and −2.99 against the three controls. Open circles, filled circles, open triangles, and open squares denote normal, KC, viral, and bacterial/fungal samples. **(B)** Weighted cosine similarity of 42 experimental contrasts to KC−normal, KC−viral, and KC−bacterial/fungal expression vectors, with genes and weights fixed. Rows align contrasts across controls and are grouped by dataset. Black marks identify classifications reversed from the normal comparison: 27 with viral and 21 with bacterial/fungal controls. **(C)** Paired KC−normal and KC−alternative expression effects for 1,090 jointly estimable candidates from the published 1,253-candidate pool. Gray density cells show candidate counts. The accompanying sign tables identify 361 viral-control and 385 bacterial/fungal-control reversals in hatched cells. The other 163 candidates are outside the jointly estimable set. **(D)** Contribution of the control change to each model decision. Open and filled points show the original margin (m_0_) and the substituted margin on the same denominator (m_1_,fixed), linked within contrast. The latter equals m_0_ − q, where q is the projection of the alternative-control−normal expression difference onto the model vector. A reversal occurs when this projection crosses the original margin. Heatmap rows show signed gene contributions to q, grouped into 80 ordered bins per model; columns do not align genes across models. The two control substitutions give 84 model–control comparisons. Supplementary Methods gives the equation, contribution definitions, and display details. Table 2 and Supplementary Table S12 report decisions and decompositions. KC, keratoconus; BF, bacterial/fungal; log_2_FC, log_2_ fold change.

**Table 2.** Control-dependent decisions and recovery with normal references.

| A. Fixed KC, substituted controls |  |  |  |  |  |
| Replacement control | Model contrasts reversed, n/N | Candidate nominations reversed, n/N |  |  |  |
|---|---|---|---|---|---|
| Viral | 27/42 | 361/1,090 |  |  |  |
| Bacterial / fungal | 21/42 | 385/1,090 |  |  |  |
| **B. Held-out normal controls** |  |  |  |  |  |
| **Old control** | **Models corrected** | **Models newly wrong** | **Candidates corrected** | **Candidates newly wrong** | **Gene agreement, old → normal** |
| Viral | 20 | 4 | 238 | 138 | 58.81% → 77.38% |
| Bacterial / fungal | 17 | 5 | 269 | 133 | 59.59% → 77.37% |
| **C. Independent expression tests** |  |  |  |  |  |
| **External truth / scope** | **Reference / comparator** | **Common genes** | **Reference BA, %** | **Comparator BA, %** | **Difference, pp (95% CI)** |
| GSE303036 stroma; common genes | Control-source reference / MetaIntegrator (nine arms) | 14,304 | 60.60 | 59.33 | 1.27 (0.46, 1.96) |
| GSE303036 stroma; top 200 | Control-source reference / MetaIntegrator (nine arms) | 175 | 84.55 | 78.26 | 6.29 (5.04, 6.29) |
| Du 2025; common genes | Selected normal reference / Niu (ComBat-limma) | 3,427 | 93.09 | 91.64 | 1.45 (0.44, 2.45) |
| Sun 2026; common genes | Selected normal reference / Niu (ComBat-limma) | 2,875 | 92.27 | 89.92 | 2.35 (1.16, 3.48) |
| **D. Recovered external decisions** |  |  |  |  |  |
| **External view** | **Decision object** | **Corrected, n/N** | **Newly wrong, n/N** | **Not estimable, n** |  |
| Du 2025 | Model contrasts | 9/42 | 1/42 | 0 |  |
| Du 2025 | Candidate nominations | 171/872 | 1/872 | 381 |  |
| GSE303036 stroma | Model contrasts | 8/42 | 8/42 | 0 |  |
| GSE303036 stroma | Candidate nominations | 76/1,063 | 150/1,063 | 190 |  |

A retains the same four KC cases. Counts refer to model contrasts or candidate nominations: 42 model contrasts, 1,253 original nominations, and 1,090 commonly estimable nominations.
B reports medians over 35 overlapping splits of seven normal donors into three reference and four referee donors, with four fixed KC cases. The four decision categories sum to 42 within each split; marginal medians need not sum to 42. Quartiles in the machine data use R type 7.
C compares methods within the same external truth, scope, and common gene universe. Balanced accuracy is a percentage; differences and confidence intervals are percentage points. Spatial intervals resample donors within diagnosis; Du and Sun intervals resample published gene rows with paired draws.
D compares the historical whole-cornea/stroma postmortem reference with the selected normal reference. An exactly layer- and source-matched historical route is unavailable in both views. Not-estimable nominations are excluded from the estimable denominator.
Abbreviations: KC, keratoconus; BA, balanced accuracy; CI, confidence interval; pp, percentage points.

The same effect occurred with 25 KC corneas held fixed in GSE77938: substitutions among six pathological control classes reversed 14 of 42 model classifications and 773 of 1,224 candidate directions at least once, and excluding the single Fuchs graft or keeping only the five classes retained by the dataset’s authors^14^ still left 11 model and 47 or 63 candidate reversals (Supplementary Results).

Control source also explained opposing matrix estimates across studies. ECM expression was higher against postmortem donors (+0.359 log_2_FC; 95% CI, 0.066–0.653) and lower against pathological controls (−2.340; −2.806 to −1.874). The opposing matrix conclusions of Kabza et al.^14^ and Shinde et al.^10^ corresponded to these two comparisons (Supplementary Table S7). The dependence was not confined to the stroma: in a keratoconus tissue microarray, epithelial EGFR and 8-OHdG staining was above that of Fuchs endothelial corneal dystrophy corneas and below that of autopsy corneas.^87^

This dependence carried into published candidate selection. Chao’s cargo pipeline^15^ ranked COL1A1 first using GSE77938, whose pathological graft controls were labeled healthy in CorneaMD (corneamd.cn/transcriptomics/4/). Replacing only its differential-expression component with the postmortem reference reversed 286 of 1,210 estimable candidate directions and moved COL1A1 to sixth; the pathological-reference replay reversed none of 1,144 (Supplementary Fig. S1B).

### Stromal Programs Are Downregulated Despite Regional Activation

Because direction depended on the control, we asked which tissue programs remained altered across control sources and where those changes occurred. Across 182 donors, inflammatory, immediate-early, and TGF-β scores were lower in KC (Hedges’ *g*, −0.519 to−0.929), as was inferred AP-1/IEG activity under CollecTRI and DoRothEA. Reductions were strongest in whole cornea/stroma; epithelial arms showed no pooled difference (Fig. 2A). All three programs remained lower against postmortem and pathological controls, and immediate-early and TGF-β reductions persisted against living-source controls. Du’s independent DALK cohort reproduced the lower inflammatory and immediate-early expression despite reduced matrix expression against eye-bank controls (Supplementary Fig. S2A; Supplementary Tables S6 and S7).^30^

**Figure 2.**
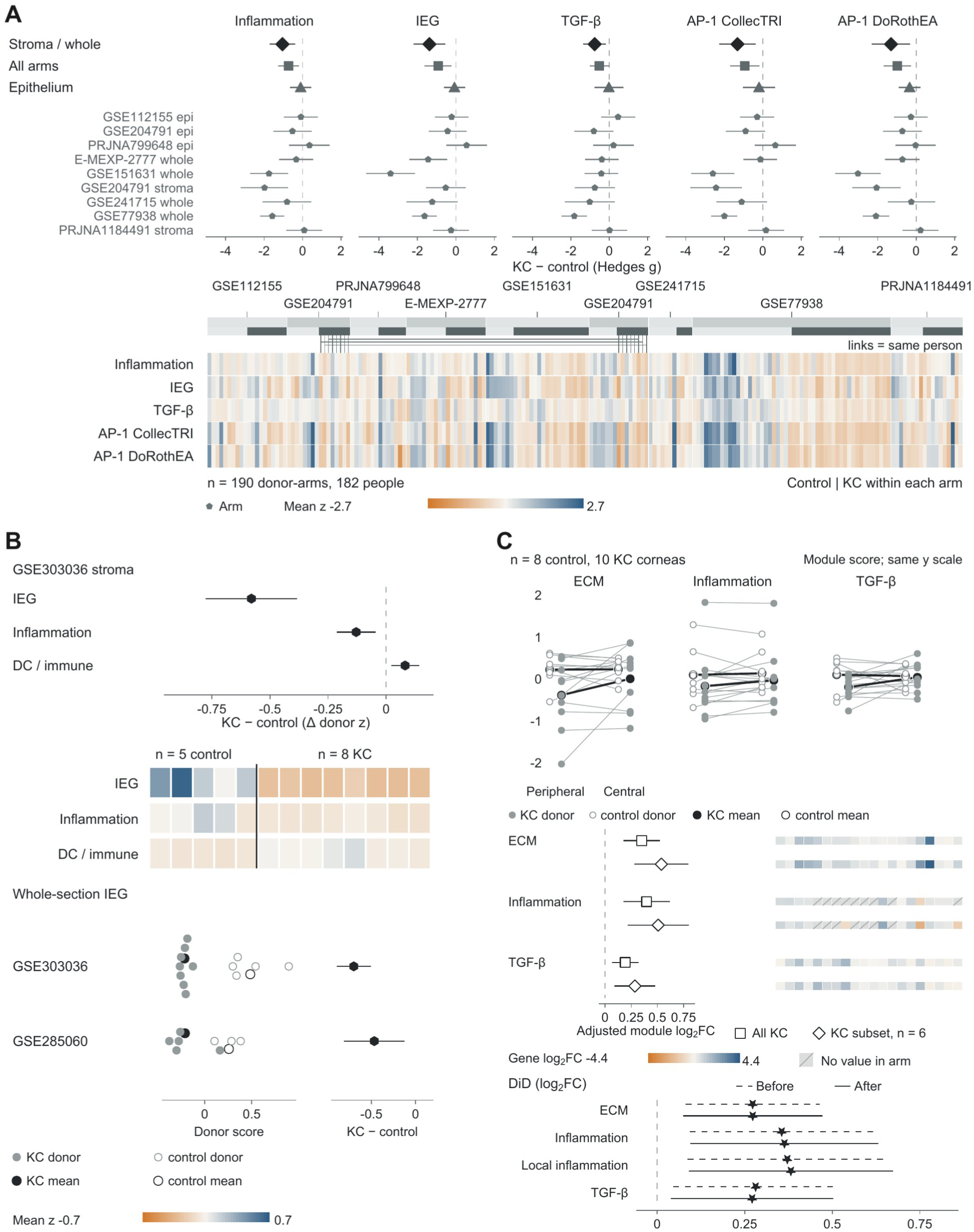
Lower stromal programs coexist with immune-marker increases and regional activation in keratoconus. **(A)** Donor-level inflammation, immediate-early (IEG), TGF-β, and inferred AP-1/IEG activity under CollecTRI and DoRothEA A–C regulons. Dark diamonds, squares, and triangles show pooled Hedges’ *g* for whole cornea/stroma, all arms, and epithelium; lighter pentagons show arm-specific estimates. Intervals are 95% CIs; negative effects indicate lower scores in KC. The heatmap displays 190 donor–arm observations from 182 people (93 KC, 89 controls); lines connect the eight KC donors sampled in both layers. Controls precede KC within each arm. Scores are means of standardized gene expression or factor activity; warm and cool colors denote lower and higher scores. **(B)** GSE303036 stromal donor-mean differences for IEG, inflammation, and DC/immune markers (KC−control: −0.580, −0.129, and +0.081), with 95% CIs and scores for the same 13 donors (8 KC, 5 controls). Below, whole-section IEG scores are shown for GSE303036 and GSE285060 (5 KC, 3 controls). Small points represent donors and larger circles group means; filled and open symbols denote KC and controls. Differences are −0.700 and −0.468, with donor-model CIs and exact donor-label permutation *P* values of 7.8 × 10^−4^ and 0.036. Whole-section and stromal GSE303036 analyses use the same donors. **(C)** Paired peripheral and central module scores in 10 KC and 8 control corneas. Lines join regions within a cornea; larger symbols show group means. The forest displays epithelial-marker-adjusted central−peripheral module log_2_FC and 95% CIs for all 10 KC corneas (squares) and the six-cornea debrided subset (diamonds). Gene strips show 20 ordered positions per module and contrast; hatching denotes missing effects. Below are differences between KC and control regional effects before (dashed) and after (solid) epithelial adjustment, with 95% CIs. Module scores and gene-model log_2_FC have separate scales. Definitions and results are in Supplementary Tables S2, S6, and S8–S10; gene identities and order are in the Figure 2C source-data table. KC, keratoconus; IEG, immediate-early genes; TGF-β, transforming growth factor-β; AP-1, activator protein-1; ECM, extracellular matrix; DC, dendritic cell; DiD, difference-in-differences; CI, confidence interval.

Lower stromal programs coexisted with greater immune-marker abundance. Both spatial cohorts had lower donor-mean immediate-early scores (Δz, −0.700 and −0.468; exact *P* = 7.8 × 10^−4^ and 0.036). Within the same stromal sections, inflammatory and immediate-early scores decreased while DC/immune markers increased (Fig. 2B). Unpooled corneal proteomes also had lower inflammatory and ECM scores, while immune-effector proteins as a class did not differ (Supplementary Fig. S2C).^88^ Cone stroma nevertheless contained increased IL-18 and MMP-9.^89^

Regional activation occurred within this lower overall state. Central ECM, inflammatory, and TGF-β expression exceeded peripheral expression in paired KC corneas, including after epithelial adjustment and in debrided stroma; peripheral inflammation remained below donors (Fig. 2C). Epithelial inflammation increased with severity (Supplementary Table S10). KC also opposed the inflammatory and AP-1 activation of bacterial/fungal keratitis (KC−infection Δ*g*, −3.357 to −4.176; all *P* < 4.3 × 10^−8^).

Global resemblance to KC did not ensure these stromal programs. Acute corneal-cell stretch increased immediate-early, AP-1, and PTGS2 expression and opposed pooled KC tissue (cosine, −0.163; 95% CI, −0.187 to −0.139),^12^ whereas PTGS2 was lower in KC tissue against all three stromal control classes. Week-scale mouse skin expansion likewise became KC-like while immediate-early and AP-1 expression rose (Supplementary Fig. S3C –G).

### Collagen Assembly and Corneal Resistance Define Structural Susceptibility

We asked whether inherited susceptibility lies in the same programs or in the matrix, whose expression direction changed with the control. ECM was the only prespecified module enriched in both KC GWAS and the FinnGen R13 corneal-deformation endpoint (H7_CORNEALDEFORM; joint MAGMA *P* = 1.03 × 10^−11^ to 5.46 × 10^−4^ across Hardcastle, MVP European, and FinnGen; Supplementary Fig. S4A). Among 7,132 public pathways, collagen assembly was the only independently replicated family (Fig. 3A). Inflammatory, immediate-early, type-2 atopy, and TGF-β sets were not enriched. No tested set combined replicated genetic enrichment with a pooled tissue-expression change (Supplementary Tables S3, S6, and S7).

**Figure 3.**
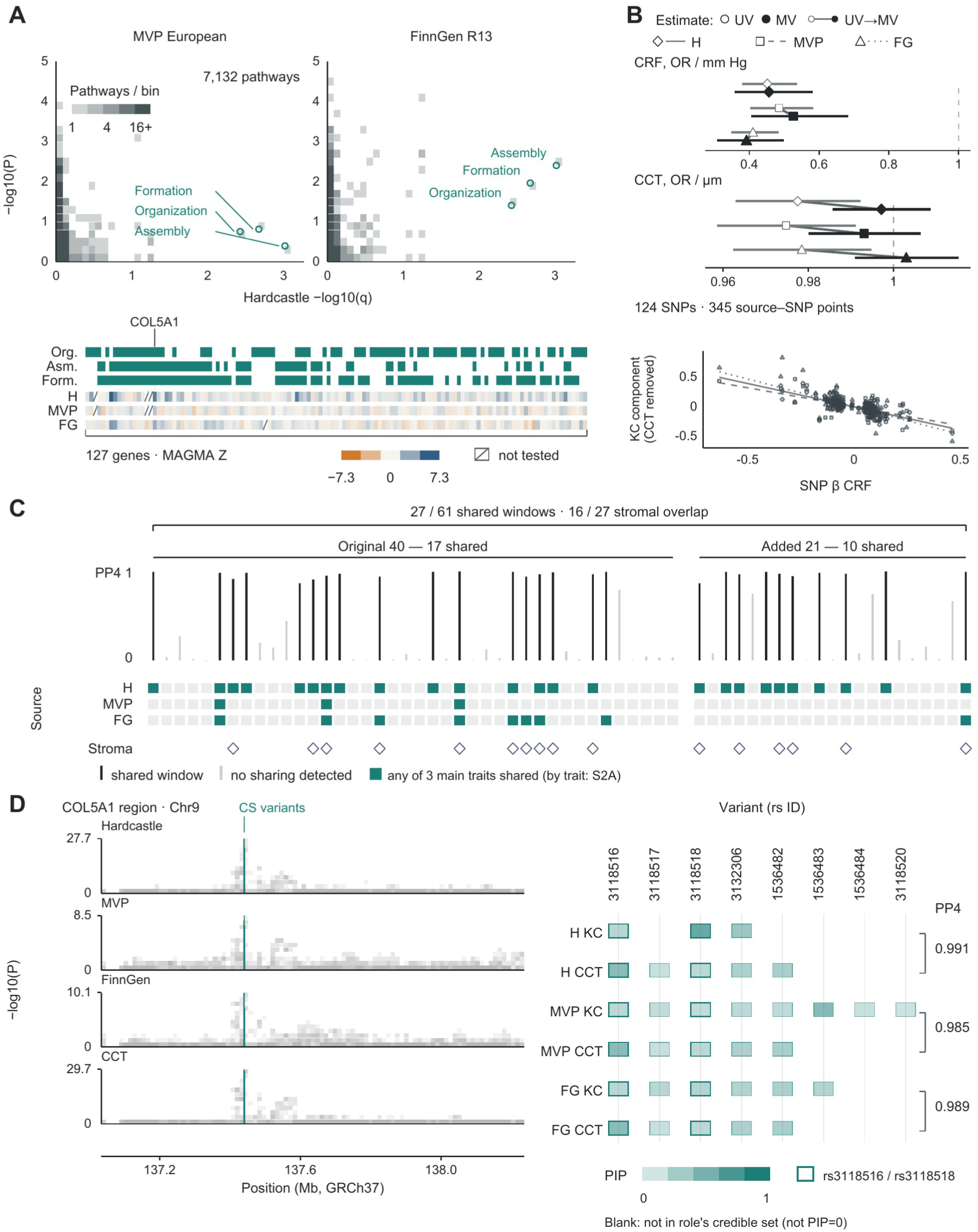
Collagen pathways, corneal resistance, and shared variants identify structural susceptibility to keratoconus. **(A)** Competitive MAGMA results for 7,132 pathways, comparing Hardcastle discovery *q* values with independent-source *P* values in MVP European and the FinnGen R13 corneal-deformation endpoint (H7_CORNEALDEFORM). Discovery required positive enrichment and *q* < 0.05; replication required positive enrichment and *P* < 0.05 in either independent source. Gray cells show pathway counts. Teal circles mark collagen fibril organization, assembly, and formation, which replicate in FinnGen. The three overlapping terms form one descriptive family. Membership strips and gene Z statistics share 127 gene columns; hatching marks untested genes. **(B)** Univariable (open symbols) and joint CRF–CCT (filled symbols) MR estimates with 95% CIs. Diamonds, squares, and triangles denote Hardcastle, MVP, and FinnGen; connecting lines pair each source’s estimates. ORs are per 1 mm Hg higher CRF or 1 µm higher CCT. Below, 345 source–SNP records from 124 distinct SNPs plot CRF effects against the disease component after subtracting the fitted CCT contribution. Lines through the origin use the multivariable CRF coefficients; solid, dashed, and dotted lines denote the three sources. **(C)** Structural sharing in the original 40 regions and 21 additional candidate windows. Seventeen original and ten additional windows are shared; 16 of the 27 have stromal regulatory context. Bar height is the largest PP4 across the three disease sources and updated CCT, CRF, or genetically CCT-residualized CRF. Teal source cells denote sharing and open diamonds stromal context. Windows are ordered by chromosome and start within the original and added groups. Trait-specific results are in Supplementary Figure S5A. **(D)** COL5A1 association tracks for the three disease sources and original CCT (GCST006366), on GRCh37 chromosome 9. Gray density cells show SNP counts by position and −log_10_(*P*); teal lines locate credible-set variants. The matrix shows eight variants across the three primary CCT component pairs, with separate disease and CCT rows. Color represents original PIP; blank cells indicate nonmembership. Outlines mark rs3118516 and rs3118518. Brackets give pairwise PP4: 0.991, 0.985, and 0.989 for Hardcastle, MVP, and FinnGen. Supplementary Figure S5B expands component membership; S5D shows the separate regional accessible-element annotation. Methods and complete estimates are in Supplementary Methods and Supplementary Tables S3–S5. MAGMA, Multi-marker Analysis of GenoMic Annotation; H, Hardcastle; MVP, Million Veteran Program; FG, FinnGen R13 corneal deformation; KC, keratoconus; MR, Mendelian randomization; UV, univariable; MV, multivariable; CRF, corneal resistance factor; CCT, central corneal thickness; CRF⊥CCT, genetically CCT-residualized CRF; OR, odds ratio; SNP, single-nucleotide polymorphism; PP4, posterior probability of a shared causal variant; PIP, posterior inclusion probability; CS, credible set; CI, confidence interval.

Corneal resistance contributed to susceptibility independently of thickness. Genetically determined CRF reduced disease odds across all three sources, and CAUSE favored causal over shared-pleiotropy models, while reverse MR linked disease liability to lower resistance. Joint modeling of CRF and CCT retained the protective CRF effect (ORs, 0.392–0.527 per mm Hg), whereas CCT had no independent effect (*P* = 0.31–0.63; Fig. 3B; Supplementary Fig. S4D). Joint structural-trait projections captured 35.5%–54.3% of common-SNP disease heritability in Hardcastle and 27.4%–38.6% in FinnGen (Supplementary Fig. S4C).

### Shared Structural Signals Extend to Stromal Regulatory Elements

We then traced the shared disease–structure signals to adult corneal regulatory regions and a functionally tested COL5A1 fragment. Seventeen of the original 40 risk regions shared signals with corneal structural traits; adding published leads extended sharing to 27 of 61 windows.^90^ Sixteen shared windows contained credible variants in adult stromal class-enriched peaks (Fig. 3C). The original shared-region set was enriched in keratocyte and stromal stem-cell open chromatin (1.88- and 1.94-fold; *P* = 0.015 and 0.013), with no enrichment in basal or superficial epithelial peaks. Accessible elements occurred at ALDH3A1, FOXO1, SMAD3, and COL5A1 (Supplementary Fig. S5C,D). FNDC3B also shared signals (Supplementary Fig. S5A).

At COL5A1, which encodes collagen V, the primary CCT component was shared across all three disease sources (PP4, 0.985–0.991), with rs3118516 and rs3118518 in both credible sets of each pairing (Fig. 3D; Supplementary Fig. S5B). Five shared variants (rs3118518, rs3132306, rs1536482, rs1536483, and rs1536484) lay within a 938-bp reporter fragment containing 0.56–0.71 of the variant posterior conditional on sharing in each source’s primary CCT component. At the four variants without strand ambiguity, the KC risk allele lay on the CCT-lowering haplotype in every source. That haplotype had 3.9-fold lower reporter activity in hTK keratocytes (7.9 versus 31.1; *P* = 0.004) and lower activity in RITVA cells (0.9 versus 2.9; *P* = 0.02), with no allelic difference in RPE1 cells.^54^ The shared signal therefore maps to a fragment with haplotype-dependent regulatory activity.

### TGF-β Output Is Lower With Weaker Feedback, and KC Cells Still Induce Matrix Production

Latent TGF-β is stored in the stromal matrix, and cells activate it by pulling on it, which works only if the matrix resists the pull.^91−94^ In mouse cornea with half the normal collagen V, collagen was disorganized and myofibroblasts could not activate latent TGF-β mechanically, although they responded to added TGF-β1.^91^ At COL5A1, the haplotype carrying KC risk had lower reporter activity, which predicts less collagen V. If inherited weakness acts through this step, KC stroma should show less TGF-β output, no extra inhibitory feedback, and cells that can still respond.

TGF-β output was lower (Fig. 2A), as was SMAD3 target output without genes shared with the immediate-early score (*g*, −0.620; 95% CI, −1.118 to −0.123; Supplementary Table S6). SMAD7, the inhibitory feedback, was lower at the same donor-level SMAD3 target output across seven independent studies (adjusted KC difference, −0.601 log_2_ units; 95% CI, −0.975 to −0.227; *P* = 0.0016), also after adjusting for the immediate-early score (−0.570; 95% CI, −0.914 to −0.226; Supplementary Table S8), whereas SMAD6 showed no pooled difference (Fig. 4A). Latent-complex storage and activation genes covaried with SMAD3 output. After removing shared genes from the output score and adjusting for diagnosis and SMAD7, output remained positively associated with collagen assembly (standardized slope, 0.570; 95% CI, 0.400–0.741; Fig. 4B).

**Figure 4.**
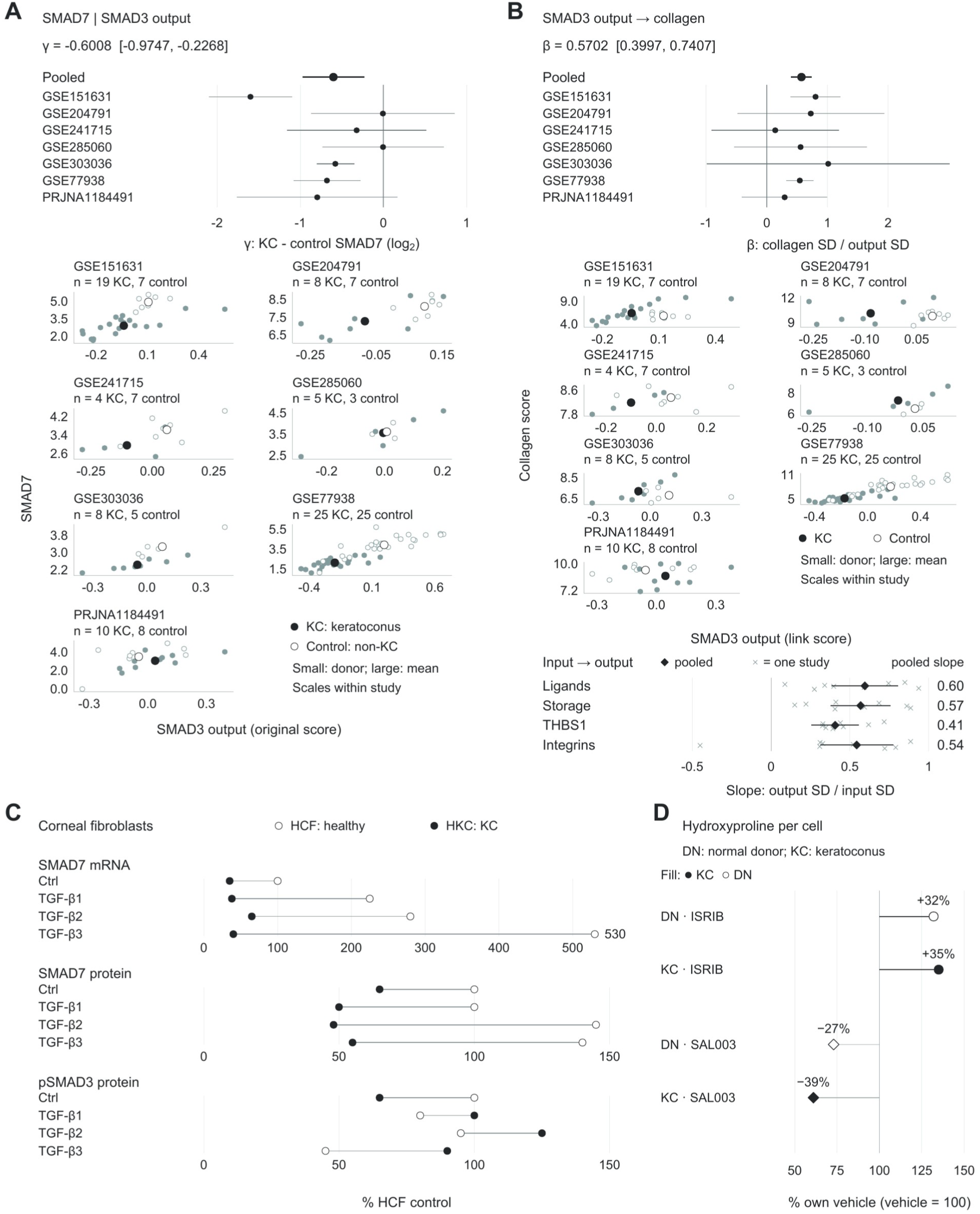
Weaker inhibitory feedback coexists with a positive output–collagen relationship and inducible matrix production. **(A)** Study-specific and pooled KC−control SMAD7 differences conditional on donor-level SMAD3 target output. The seven-study pooled coefficient is −0.6008 log_2_ units (95% CI, −0.9747 to −0.2268). Scatterplots show unadjusted output and SMAD7 expression in 141 records (79 KC, 62 controls); small circles are donors and larger circles group means. Filled and open symbols denote KC and controls. Raw scales are study-specific; the forest shows adjusted diagnosis coefficients. **(B)** SMAD3 output–collagen assembly slopes adjusted for diagnosis, SMAD7, and study covariates. Eight genes shared with the matrix outcome scores were removed from the output score. The seven-study pooled slope is 0.5702 collagen SD per output SD (95% CI, 0.3997–0.7407). Scatterplots show donor-level scores for the same records as A. Below, diamonds and CIs show pooled relationships of ligand, latent-storage, THBS1, and integrin expression with SMAD3 output; crosses show study estimates. These input–output analyses use eight studies and 161 donor records. All slopes describe donor-level RNA associations; pooled estimates in A and B use DerSimonian–Laird random effects. **(C)** Published SMAD7 mRNA, SMAD7 protein, and phosphorylated SMAD3 means after four-week control or TGF-β1, TGF-β2, or TGF-β3 treatment.^68^ Open and filled circles denote normal-derived HCF and KC-derived HKC; lines pair sources under the same treatment. Each endpoint is normalized to its HCF control. The 24 means were digitized from the published figures; the source reports at least three repeats. Extraction precision and reporting details are in Supplementary Methods. **(D)** Published cell-number-normalized hydroxyproline after ISRIB (circles) or SAL003 (diamonds).^70^ Open and filled symbols denote normal donor-derived and KC-derived cultures, each normalized to its own vehicle = 100%. ISRIB values are 132% in normal-derived and 135% in KC-derived cultures; SAL003 values are 73% and 61%, respectively. The source reports five repeats. Published means are shown; numerical error estimates were unavailable. Definitions, model estimates, and published-source values are in Supplementary Tables S2, S8, and S11. KC, keratoconus; TGF-β, transforming growth factor-β; HCF, normal human corneal fibroblasts; HKC, KC-derived corneal fibroblasts; DN, donor-derived normal fibroblasts; pSMAD3, phosphorylated SMAD3; Ctrl, control; SD, standard deviation; CI, confidence interval; mRNA, messenger RNA.

Stimulation studies showed weaker inhibitory feedback alongside retained cellular responses. During four-week TGF-β treatment, normal-derived cells induced SMAD6/7, whereas KC-derived cells did not, including at the SMAD7 protein level. TGF-β3-treated KC cells nevertheless had higher phosphorylated SMAD3 than treated normal cells (Fig. 4C).^68^ A separate three-dimensional construct study also found lower SMAD6/7 and higher phosphorylated SMAD3 in KC-derived cells.^95^ Matrix production and cytoprotection remained inducible: ISRIB increased cell-number-normalized hydroxyproline by 32% in donor-derived and 35% in KC-derived cells (Fig. 4D),^70^ and sulforaphane reduced oxidative challenge–induced caspase 3/7 activity in KC-derived cells.^69^ Serum-induced immediate-early responses in normal-source cells distinguished baseline state from inducibility (Supplementary Fig. S3A,B).^71^

After a short pulse, KC-derived cells in bioprinted stroma retained feedback and matrix responses.^85^ In donor-matched RNA sequencing from three KC donors, TGF-β1 induced SMAD7 in both 2D and 3D cultures (log_2_FC, 2.44 and 0.93; adjusted P < 0.002 for each). The format-by-stimulation interaction favored collagen enrichment in 3D (normalized enrichment score [NES], 1.92; adjusted P = 0.013) and opposed myogenesis enrichment (NES, −1.37; adjusted P = 0.037). Among the 126–200 genes most strongly reduced in each of four patient tissue comparisons, TGF-β1 stimulation in 3D raised 21–59 and lowered 20–46 (adjusted P < 0.05), so recruitment of matrix transcription does not require uniform reversal of the lesion transcriptome (Supplementary Fig. S6; Supplementary Results).

The same constructs increased DNA-normalized procollagen I secretion 2.6-fold after a TGF-β1 pulse in all six KC donors. After withdrawal of the receptor inhibitor SB431542, native type I collagen per construct rose 3.3- to 4.5-fold, with increases in all 12 donors, and eleven days later delayed blockade left more collagen and lower contractile-gene expression than vehicle in 9 of 12.^85^

### Normal Controls Recover Decisions, and Reference Choice Depends on Tissue

Normal controls recovered the decisions that pathological controls altered. Across all 35 three/four splits of seven normal donors, substituting three normal donors for viral or bacterial/fungal controls corrected a median of 20 or 17 model classifications against the held-out normal comparison, while reversing 4 or 5 previously concordant classifications. Median gene-direction agreement increased from 58.81% to 77.38% and from 59.59% to 77.37%, respectively (Fig. 5A,B), and model-score Spearman correlation with the held-out comparison rose from −0.033 to 0.824 and from 0.047 to 0.826.

**Figure 5.**
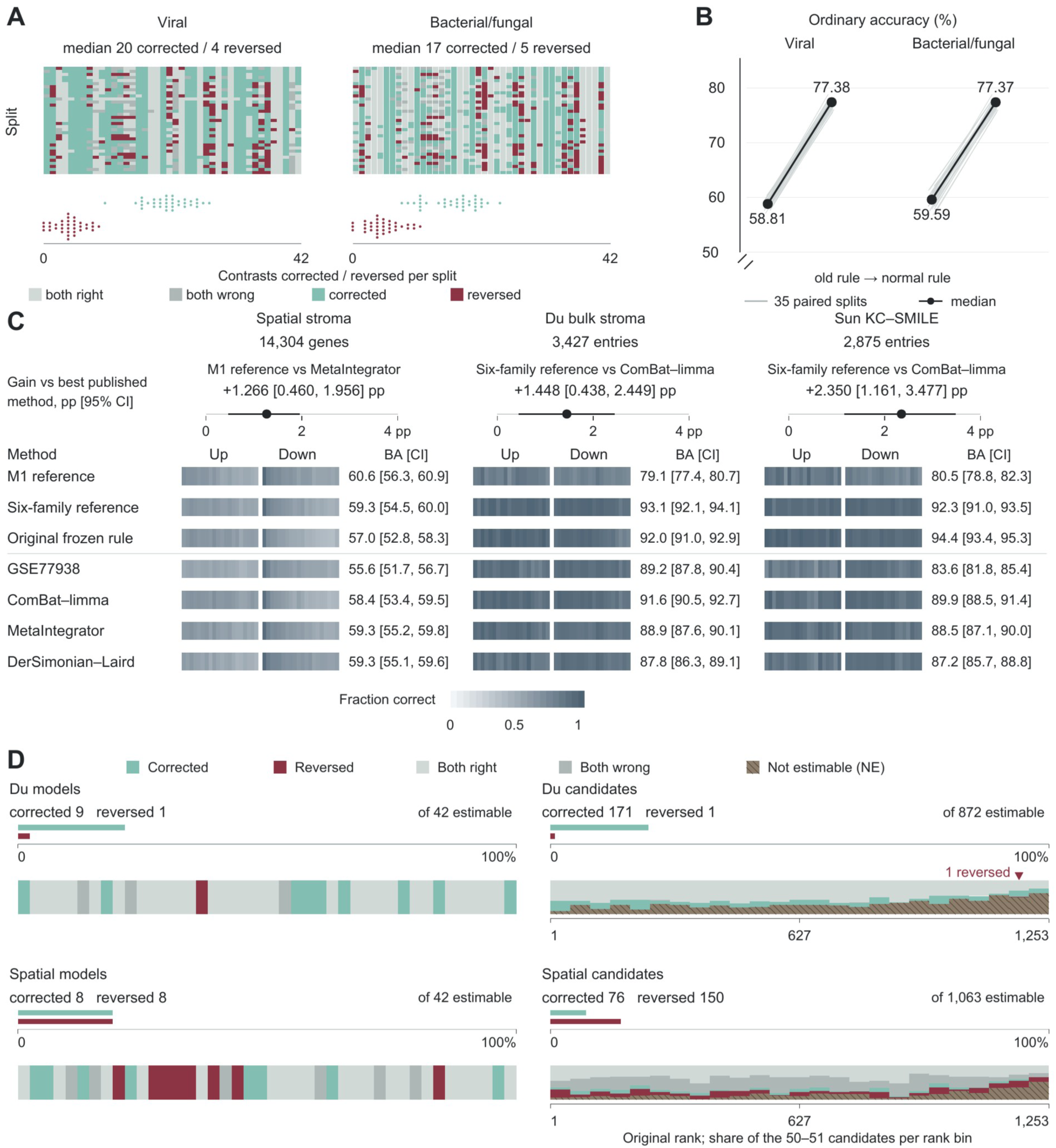
Normal controls recover decisions, and independent tissues favor different references. **(A)** All 35 partitions of seven normal donors into three reference and four held-out donors, with four KC cases fixed. Matrix cells classify each of 42 model contrasts under pathological and selected-normal controls against the held-out-normal comparison. Colors distinguish both correct, both incorrect, corrected, and reversed. Point strips show split-specific counts: median corrected/reversed counts are 20/4 for viral and 17/5 for bacterial/fungal controls. Partitions overlap, and medians are calculated separately. **(B)** Gene-direction agreement in the same paired splits. Gray lines connect pathological- and selected-normal-control values; dark points and lines show medians. Agreement increases from 58.81% to 77.38% for viral controls and from 59.59% to 77.37% for bacterial/fungal controls. **(C)** Directional balanced accuracy (BA) on common entries in independent spatial stroma (14,304 genes; 8 KC, 5 control donors), Du bulk stroma (3,427 entries; 31 KC, 4 eye-bank controls), and Sun’s KC–SMILE bulk comparison (2,875 entries; 10 KC, 10 controls). BA weights upward and downward recall equally. Direction strips summarize correct predictions in 20 equal-count bins ordered by decreasing absolute measured effect. M1 uses the fitted postmortem whole-cornea/stroma reference in all three comparisons; Sun’s measured comparison uses SMILE controls. The six-family reference was selected in whole-study holdouts. Our three reference variants appear above the line, published approaches below. The original frozen reference has higher BA than the six-family reference in Sun. Paired gains over the best published approach are 1.266 percentage points for spatial M1, 1.448 for Du’s six-family reference, and 2.350 for Sun’s six-family reference. Printed 95% CIs use 2,000 donor resamples for spatial data and paired published-entry resamples for bulk tables. Supplementary Figure S1D includes the complete method comparison. **(D)** Changes from postmortem whole-cornea/stroma M1 to the six-family reference, evaluated against independent tissue effects. Models retain all 42 contrasts; candidates retain all 1,253 records in original rank order, with unavailable entries marked NE. Against Du, the update corrects 9 model decisions and reverses 1; among 872 estimable candidates, it corrects 171 and reverses 1. Against spatial stroma, it corrects and reverses 8 models each; among 1,063 candidates, it corrects 76 and reverses 150. Summary bars use estimable denominators. The marker identifies the single reversed Du candidate, HEPHL1. Binning and display-width details are in Supplementary Methods. Table 2 and Supplementary Table S13 give all counts and paired comparisons. KC, keratoconus; BA, balanced accuracy; M1, model 1; SMILE, small-incision lenticule extraction; pp, percentage points; NE, not estimable; CI, confidence interval.

The source- and layer-based reference (M1) fits expression effects to control source and tissue layer, and each observed layer–control combination defines a reference, chosen before external scoring. In independent spatial stroma it outperformed published approaches. Across 14,304 common genes, directional balanced accuracy was 60.60%, compared with 59.33% for the best published approach (gain, 1.27 percentage points; 95% CI, 0.46–1.96). Among the 172 jointly measurable genes in the 200 strongest stromal effects, balanced accuracy was 84.5%, compared with 80.1% for Niu’s released gene-level results (difference, 4.4 percentage points; 95% CI, 3.1–4.5). It also led in endothelium, and ranked stromal, endothelial, and immune effects better than every published approach (Fig. 5C; Supplementary Table S13).

The six-family reference combines predictions from six method families, configured in whole-study holdouts and frozen before external evaluation. It outperformed published approaches in both bulk comparisons. In Du’s published differential-expression table, the source- and layer-based reference scored 79.1%, compared with 91.6% for ComBat–limma. The six-family reference reached 93.09% across 3,427 common entries (gain over the best published approach, 1.45 points; 95% CI, 0.44–2.45). Its gain reproduced in the KC–SMILE comparison (2.35 points; 95% CI, 1.16–3.48; Fig. 5C; Supplementary Fig. S1D).^31^

Reference choice depended on the tissue. Against Du’s measured effects, the six-family update corrected nine model classifications and 171 of 872 candidate directions, reversing one previously correct decision in each analysis. In spatial stroma, the same update corrected 76 of 1,063 candidate directions but reversed 150 (Fig. 5D; Table 2). The source- and layer-based reference therefore served spatial stroma, and the six-family reference bulk tables.

### Functional Benefit Can Oppose Reversal of Disease Expression

Choosing an intervention also requires its functional effect, so we first compared each published target premise with the reference estimate for its tissue and readout. Forty-one of 46 published nominations required a different comparison, measurement, or use: 36 required another measurement or comparison, and five changed use because only the opposite direction was supported. Epithelial AP-1 hubs agreed with their source comparison,^11^ while JUN and EGR1 were lower across whole-cornea/stromal control classes. Epithelial-expression and tear MMP-9 findings supported the rationale for topical cyclosporine treatment^96^ and a subsequent 362-eye cyclosporine–trehalose cohort,^97^ while MMP9 transcripts were lower in every measured KC tissue arm.

The same distinction mattered for other proposed targets. NEDD8’s upregulation premise was absent in whole-cornea/stromal comparisons, while its inflammatory rescue followed acute stretch.^98^ CCN1 and ATF4 had protein-based premises despite lower transcripts^99,100^ (Table 3; Supplementary Table S14). Niu et al.^82^ fitted a ten-gene nomogram to KC-versus-control labels in GSE151631, GSE112155, and GSE77938 but described it as predicting KC progression; a later review restated the genes as relating to prognosis.^86^

**Table 3.**
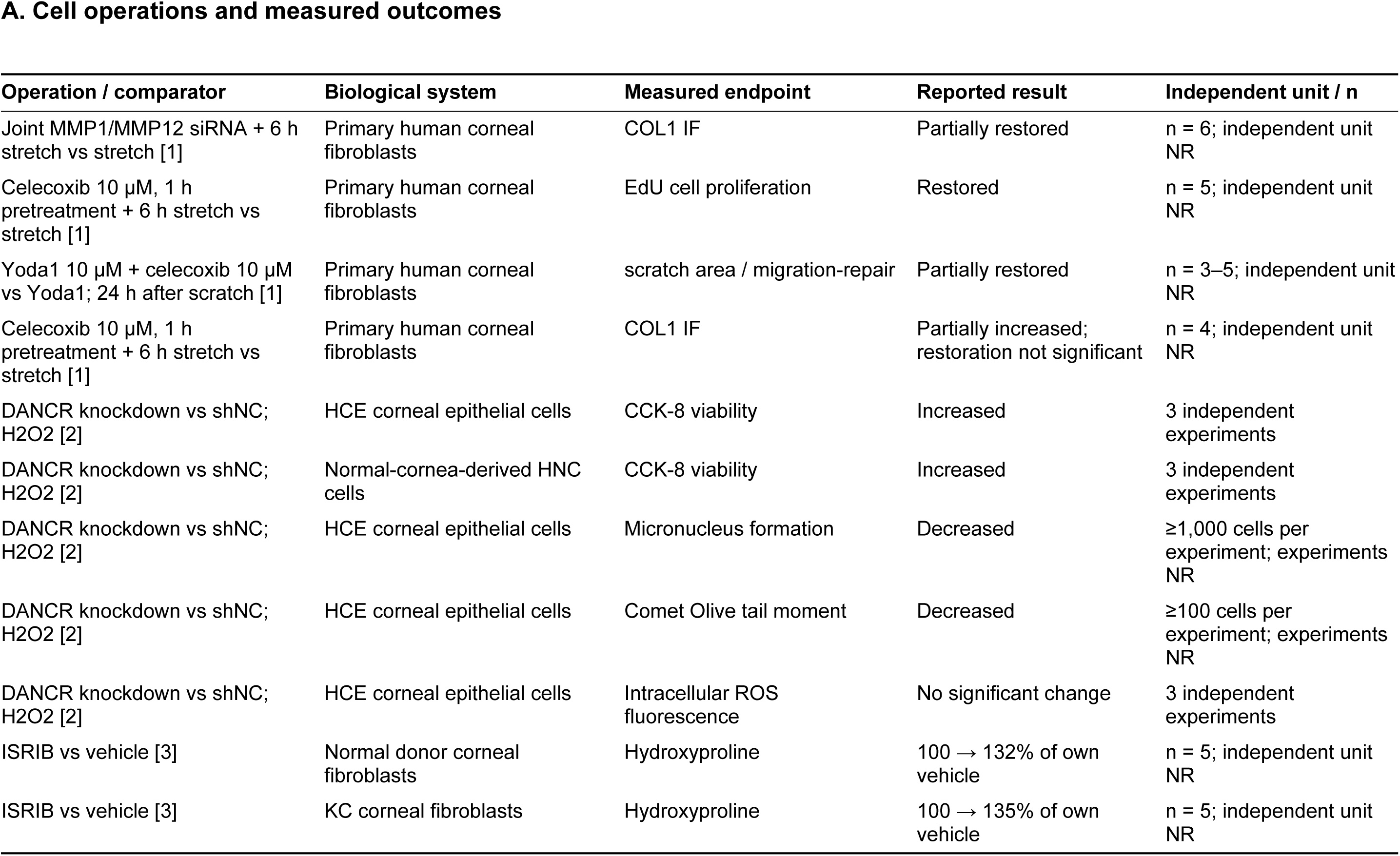
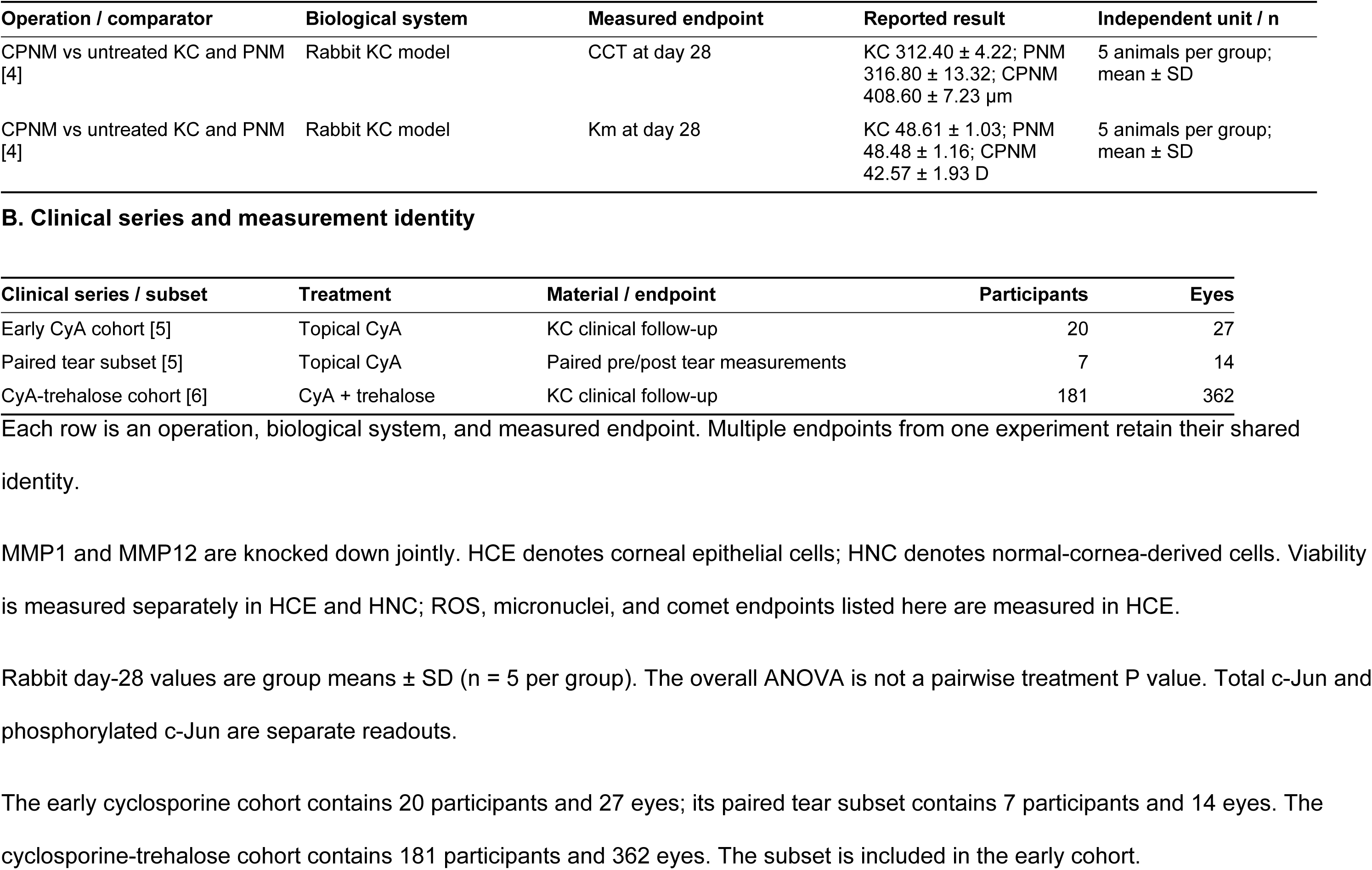
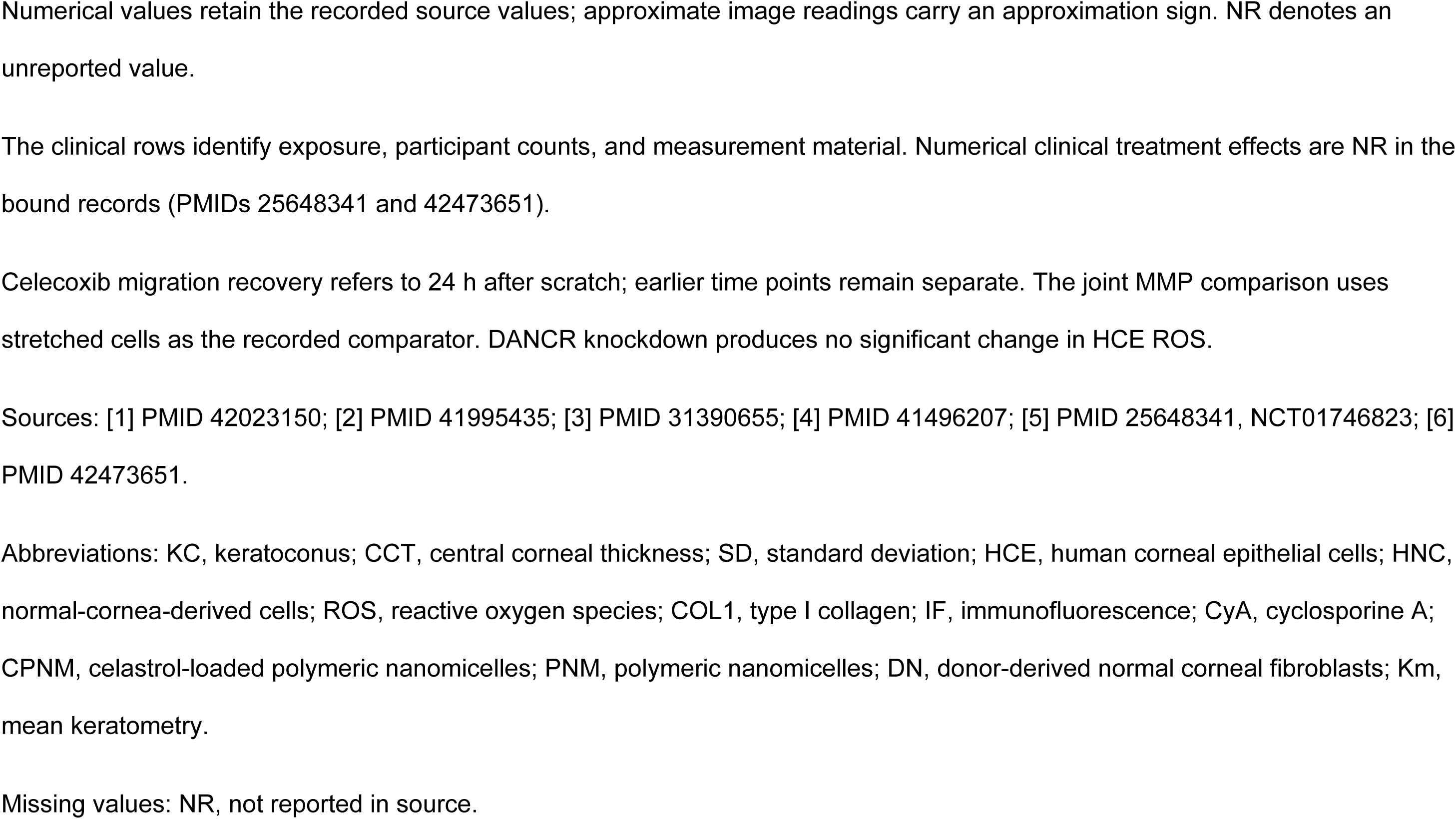
Published operations and the identity of their measured endpoints.

| **A. Cell operations and measured outcomes** |  |  |  |  |
| **Operation / comparator** | **Biological system** | **Measured endpoint** | **Reported result** | **Independent unit / n** |
|---|---|---|---|---|
| Joint MMP1/MMP12 siRNA + 6 h stretch vs stretch [1] | Primary human corneal fibroblasts | COL1 IF | Partially restored | n = 6; independent unit NR |
| Celecoxib 10 µM, 1 h pretreatment + 6 h stretch vs stretch [1] | Primary human corneal fibroblasts | EdU cell proliferation | Restored | n = 5; independent unit NR |
| Yoda1 10 µM + celecoxib 10 µM vs Yoda1; 24 h after scratch [1] | Primary human corneal fibroblasts | scratch area / migration-repair | Partially restored | n = 3–5; independent unit NR |
| Celecoxib 10 µM, 1 h pretreatment + 6 h stretch vs stretch [1] | Primary human corneal fibroblasts | COL1 IF | Partially increased; restoration not significant | n = 4; independent unit NR |
| DANCR knockdown vs shNC; H2O2 [2] | HCE corneal epithelial cells | CCK-8 viability | Increased | 3 independent experiments |
| DANCR knockdown vs shNC; H2O2 [2] | Normal-cornea-derived HNC cells | CCK-8 viability | Increased | 3 independent experiments |
| DANCR knockdown vs shNC; H2O2 [2] | HCE corneal epithelial cells | Micronucleus formation | Decreased | ≥1,000 cells per experiment; experiments NR |
| DANCR knockdown vs shNC; H2O2 [2] | HCE corneal epithelial cells | Comet Olive tail moment | Decreased | ≥100 cells per experiment; experiments NR |
| DANCR knockdown vs shNC; H2O2 [2] | HCE corneal epithelial cells | Intracellular ROS fluorescence | No significant change | 3 independent experiments |
| ISRIB vs vehicle [3] | Normal donor corneal fibroblasts | Hydroxyproline | 100 → 132% of own vehicle | n = 5; independent unit NR |
| ISRIB vs vehicle [3] | KC corneal fibroblasts | Hydroxyproline | 100 → 135% of own vehicle | n = 5; independent unit NR |
| CPNM vs untreated KC and PNM [4] | Rabbit KC model | CCT at day 28 | KC 312.40 ± 4.22; PNM 316.80 ± 13.32; CPNM 408.60 ± 7.23 µm | 5 animals per group; mean ± SD |
| CPNM vs untreated KC and PNM [4] | Rabbit KC model | Km at day 28 | KC 48.61 ± 1.03; PNM 48.48 ± 1.16; CPNM 42.57 ± 1.93 D | 5 animals per group; mean ± SD |
| **B. Clinical series and measurement identity** |  |  |  |  |
| **Clinical series / subset** | **Treatment** | **Material / endpoint** | **Participants** | **Eyes** |
| Early CyA cohort [5] | Topical CyA | KC clinical follow-up | 20 | 27 |
| Paired tear subset [5] | Topical CyA | Paired pre/post tear measurements | 7 | 14 |
| CyA-trehalose cohort [6] | CyA + trehalose | KC clinical follow-up | 181 | 362 |

Each row is an operation, biological system, and measured endpoint. Multiple endpoints from one experiment retain their shared identity.
MMP1 and MMP12 are knocked down jointly. HCE denotes corneal epithelial cells; HNC denotes normal-cornea-derived cells. Viability is measured separately in HCE and HNC; ROS, micronuclei, and comet endpoints listed here are measured in HCE.
Rabbit day-28 values are group means ± SD (n = 5 per group). The overall ANOVA is not a pairwise treatment P value. Total c-Jun and phosphorylated c-Jun are separate readouts.
The early cyclosporine cohort contains 20 participants and 27 eyes; its paired tear subset contains 7 participants and 14 eyes. The cyclosporine-trehalose cohort contains 181 participants and 362 eyes. The subset is included in the early cohort.
Numerical values retain the recorded source values; approximate image readings carry an approximation sign. NR denotes an unreported value.
The clinical rows identify exposure, participant counts, and measurement material. Numerical clinical treatment effects are NR in the bound records (PMIDs 25648341 and 42473651).
Celecoxib migration recovery refers to 24 h after scratch; earlier time points remain separate. The joint MMP comparison uses stretched cells as the recorded comparator. DANCR knockdown produces no significant change in HCE ROS.
Sources: [1] PMID 42023150; [2] PMID 41995435; [3] PMID 31390655; [4] PMID 41496207; [5] PMID 25648341, NCT01746823; [6] PMID 42473651.
Abbreviations: KC, keratoconus; CCT, central corneal thickness; SD, standard deviation; HCE, human corneal epithelial cells; HNC, normal-cornea-derived cells; ROS, reactive oxygen species; COL1, type I collagen; IF, immunofluorescence; CyA, cyclosporine A; CPNM, celastrol-loaded polymeric nanomicelles; PNM, polymeric nanomicelles; DN, donor-derived normal corneal fibroblasts; Km, mean keratometry.
Missing values: NR, not reported in source.

Neither genetic-risk expression nor disease expression specified the effect of an intervention. Risk-expression and tissue-expression directions differed for 12 of 13 genes in at least one KC arm (Fig. 6A). At FNDC3B, lower expression accompanied risk and seven of nine tissue comparisons, yet knockdown reduced stress fibers in embryonic fibroblasts^101^ and increased TGF-β1-induced stress fibers in HeLa cells,^102^ so the consequence depended on cellular context.

**Figure 6.**
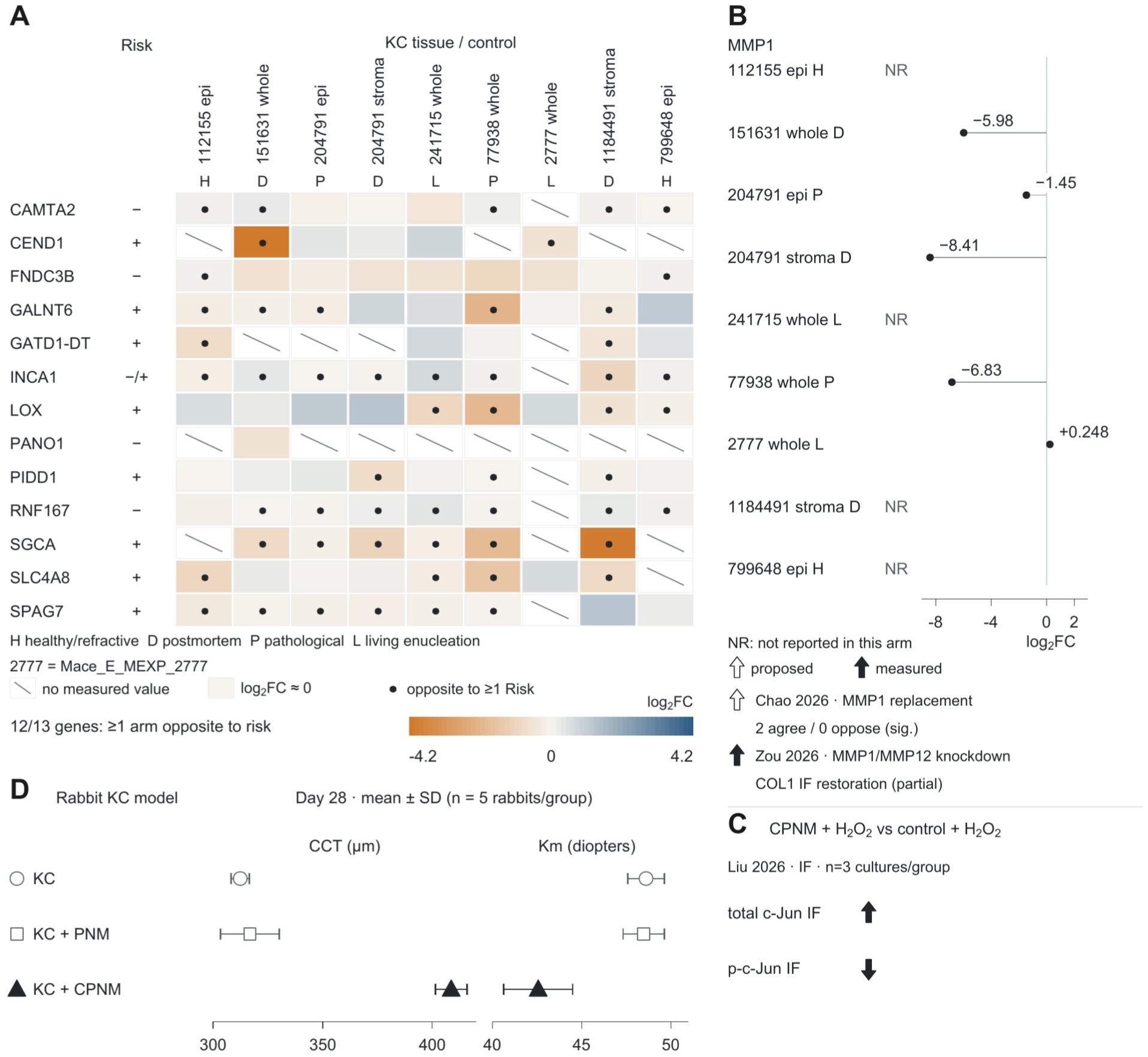
Risk-expression direction, tissue state, and functional repair outcomes are separable. **(A)** KC−control log_2_FC for 13 genes across nine tissue arms. Of 117 positions, 92 have measured effects and 25 are missing (hatched). Dots mark effects opposite to at least one accepted risk-allele expression direction; 12 genes have such an effect in at least one arm. The risk column retains both accepted directions for INCA1. Warm and cool colors denote lower and higher KC expression. Source codes identify healthy/refractive (H), postmortem donor (D), pathological (P), and living-enucleation (L) controls. GSE204791’s eight KC donors contribute both layers. **(B)** MMP1 tissue expression, replacement-cargo nomination, and response to joint MMP1/MMP12 knockdown. Five arms have measured log_2_FC; four are not reported (NR). Chao et al.^15^ nominated MMP1 to supplement low matrix expression. Two of three matching layer–control estimates significantly support that direction and none significantly opposes it. During six-hour corneal-cell stretch, joint MMP1/MMP12 knockdown partly restores collagen I immunofluorescence.^12^ Open arrows mark the nomination and filled arrows the measured outcome. **(C)** Total and phosphorylated c-Jun immunofluorescence in rabbit-derived cultures pretreated with celastrol-loaded nanomicelles (CPNM) at 1 µg/mL celastrol for 24 hours and then H_2_O_2_ (200 µg/mL, 1 hour), compared with control plus H_2_O_2_.^13^ Total c-Jun increases while phospho-c-Jun decreases. The source reports three independently established cultures from three donor rabbits per group. Arrows show reported directions. **(D)** Day-28 central corneal thickness (CCT) and mean keratometry (Km) in the published rabbit ectasia model induced by epithelial removal and type II collagenase.^13^ Symbols and error bars show means ± standard deviation (SD), using the five rabbits per group reported in the source Table 1. Open circles, open squares, and filled triangles denote untreated KC, empty nanomicelles (PNM), and CPNM. Culture readouts in C and rabbit outcomes in D have separate experimental units. Full values and source reporting details are in Supplementary Methods. Table 3 and Supplementary Table S14 identify interventions, endpoints, and source records. KC, keratoconus; epi, epithelium; stroma, corneal stroma; whole, whole cornea; log_2_FC, log_2_ fold change; siRNA, small interfering RNA; IF, immunofluorescence; CCT, central corneal thickness; SD, standard deviation.

The intervention-outcome index (Supplementary Table S14, where interventions are termed operations; 2,769 evidence records, 194 index entries) links each perturbation to its system, comparator, and measured endpoint. Linked to the 46 nominations, it held intervention records on the target itself for three (two LOX nominations, where recombinant LOX raised tendon tensile modulus without increasing collagen content,^103^ and MMP1), records only as readouts of other interventions for 22, and no record for 21. Corneal interventions directly separated transcript reversal from functional benefit. Low MMP1 expression prompted nomination as replacement cargo,^15^ yet combined MMP1/MMP12 knockdown partly restored collagen I after corneal-cell stretch.^12^ DANCR knockdown extended its low disease-expression direction while reducing oxidative DNA damage and improving survival in non-KC corneal epithelial cells.^104^ Celastrol increased total c-Jun while lowering phospho-c-Jun and improving rabbit corneal curvature and thickness.^13^ Each intervention therefore requires a measured functional endpoint; reversal of the disease transcript is a separate outcome (Fig. 6B–D; Table 3; Supplementary Table S14).

## Discussion

Read against inherited stromal weakness, gene-expression differences between KC and control corneas describe the state of the diseased stroma and not its cause. This overturns two assumptions behind current use. Direction does not belong to KC, because the same corneas gave opposite model and candidate decisions against different controls. The state is not the cause, because the lower inflammatory, immediate-early, and TGF-β programs that persisted across controls had no genetic enrichment, whereas inherited risk centered on collagen assembly and resistance to deformation. Expression therefore needs a defined comparison before it guides a model and a measured functional effect before it guides an intervention.

Genetic studies had implicated collagen matrix integrity^3,6^ and CRF colocalization stromal regulation;^58^ we identify resistance to deformation as the thickness-independent protective component. This weakness can reach the signaling state through TGF-β activation: cells release latent TGF-β from the matrix by pulling on it, so a matrix with low resistance releases less active TGF-β, as in mouse cornea with half the normal collagen V.^91^ The COL5A1 risk haplotype has lower reporter activity, which predicts the same weakness in KC. If the inherited weakness acts through the same step, KC tissue should show less active TGF-β acting on cells that can still respond and are not held back by extra braking, and it did. TGF-β programs, which active TGF-β drives, were lower; SMAD7, the inhibitory feedback, was lower, not higher, than the SMAD3 target output predicted; and stimulated KC-derived cells increased matrix production.^68,70,95^ Loading did not lower the programs: stretch and skin expansion raised immediate-early and AP-1 expression. The low signal is therefore the result of a weak matrix, not a cause of the disease.

The first remedy is to state the comparison. Direction exists only against a control, so a reference specified by control source and tissue layer turns disagreement between studies into a measurable difference of comparison. Shinde’s higher and Kabza’s lower matrix expression corresponded to comparisons against postmortem donors and pathological grafts, respectively,^10,14^ as pooled ECM expression reversed between these controls. AP-1 depends on the layer in the same way: epithelial hubs agreed with their own comparison,^11^ whereas JUN and EGR1 were lower across whole-cornea and stromal controls. Local inflammatory findings^9,29^ reflect regional activation within a lower overall state. Such references predicted direction more accurately than published methods in independent tissue, and against them 41 of 46 published nominations needed another comparison, measurement, or use.

The second remedy is to choose interventions by what they do. Because the state is not the cause, reversing an expression change need not help, and published interventions separate the two. Low MMP1 expression prompted nomination as replacement cargo,^15^ yet MMP1/MMP12 knockdown partly restored collagen I after stretch.^12^ The intervention-outcome index records such outcomes by system and endpoint; it holds an intervention on the target itself for only 3 of the 46 nominations, which therefore still need measured outcomes, not expression direction. Interventions that raise matrix resistance have the record that resistance weakness predicts: corneal cross-linking halted progression in randomized trials,^105,106^ and recombinant LOX raised tensile modulus without increasing collagen content.^103^

These findings change what KC expression data can be used for. A model is tested against a reference matched to control source and tissue layer, and a target is chosen from the measured outcomes of named interventions. Because inherited risk lies in matrix resistance and KC-derived cells retain matrix production, the repair objective is organized, load-bearing matrix, judged by matrix organization, resistance to deformation, corneal shape, optical quality, and progression rather than by reversal of an expression change. In this account KC is a disease of the matrix’s resistance to deformation, and expression records how the stroma responds to that weakness; treatments should therefore be designed and judged by their effect on the matrix.

## Acknowledgments

The authors thank the investigators and consortia who deposited the public datasets listed in Supplementary Table S1.

## Use of artificial intelligence

The authors used Claude Opus 5.5 (Anthropic) and Astra 6 (OpenAI) for language editing and coding assistance between June 2026 and November 2026. The authors reviewed and verified the outputs and take full responsibility for the content of this work.

## Author Contributions (CRediT)

Hanyu Jiang: Conceptualization, Data curation, Formal analysis, Investigation, Methodology, Software, Validation, Visualization, Writing – original draft, Writing – review & editing.

Fei Gao: Conceptualization, Methodology, Writing – review & editing.

Wei Zhang: Conceptualization, Methodology, Writing – review & editing.

Ying Jie: Supervision, Writing – review & editing.

Ying Li: Supervision, Writing – review & editing.

Yang Jiang: Supervision, Writing – review & editing.

All authors read and approved the final manuscript.

## Data and Code Availability

The core analysis used public source data. GWAS summary statistics are available under GCST90013442, GCST90477661, GCST90480060, GCST006366, GCST90000654, GCST90100568, and FinnGen R13 endpoint H7_CORNEALDEFORM. Tissue and disease transcriptomes are available under GSE151631, GSE77938, GSE112155, GSE241715, GSE204791, E-MEXP-2777, PRJNA1184491, PRJNA799648, GSE58291, GSE142538, GSE171830, and GSE155776. Perturbation data are GSE260476 and GSE287915; mechanical, load-release, feeder, and handling series are SRP648618, GSE58389, GSE124138, GSE284808, GSE186773, PRJNA753961, GSE308374, GSE262420, GSE310514, GSE324558, and GSE337743; adult corneal chromatin accessibility is GSE155683; methylome data are GSE107189; m6A data are GSE181618; and spatial data are GSE303036 and GSE285060. Independent bulk validation used the published Supplementary Table S4 of Du et al.^30^ and Supplemental Table S5 of Sun et al.^31^ Culture effect-table inputs and the expanded published-lead window list are identified in Supplementary Methods. Public-pathway membership came from MSigDB 2026.1.Hs. Dataset identities and analytic subsets are specified in Supplementary Table S1, and derived estimates are listed in Supplementary Tables S1–S16. Code and derived data are archived at Zenodo.

---

**Supplementary Figure S1. Complete model projections, candidate replays, and independent reference comparisons retain their object-specific outcomes. (A)** Signed weighted cosine similarity of 42 experimental contrasts to five KC−control reference vectors, grouped into 17 datasets. The reference codes identify the control comparison: P, postmortem whole cornea/stroma; D, pathological whole cornea/stroma; E, living-enucleation whole cornea/stroma; R, healthy/refractive epithelium; K, pathological epithelium. Filled dots identify rank 1 within each reference; intervals are not shown. The irradiated 3T3-J2 feeder contrast GSE324558 LQ−SQ ranks 9, 6, 9, 41, and 4, respectively, among 42 contrasts. References and contrasts reuse inputs. Supplementary Table S12 reports projections. **(B)** Original versus replayed candidate ranks after replacing only the published cargo pipeline’s differential-expression component. All other evidence and weights are retained. Against the postmortem reference, 286 of 1,210 estimable candidates change supplementation/inhibition direction, with 43 NE; against the pathological reference, 0 of 1,144 change, with 109 NE. Each begins from 1,253 candidate records. Ranks are within each estimable subset. Density cells count candidates in bins of at most 42 rank positions, with common square-root shading on a 0–35 count scale. Individual flip points are overlaid. The outlined COL1A1 point moves from rank 1 to rank 6 in the postmortem subset; the diagonal marks equal ranks. Numerical summaries are in Supplementary Table S12. **(C)** Candidate decisions across the 35 overlapping partitions of seven normal donors (three selected, four held out), with four KC cases fixed. Filled and open donor-strip cells identify selected and held-out donors. The two matrices retain 87,710 records (35 splits × 2 old-control groups × 1,253 candidates), ordered by the original normal-comparison rank. Each of 114 bins contains 11 original rank positions, except the last with 10. Color represents (corrected − reversed)/all candidate records in the bin; both-correct, both-incorrect, and NE records remain in that denominator. Teal indicates a positive signed proportion and brown a negative proportion. Separate NE strips show the mean of the 35 split-specific NE proportions per bin. Median corrected/reversed/NE counts are 238/138/199 for viral and 269/133/197 for bacterial/fungal controls; separately computed medians need not sum to 1,253. Supplementary Table S13 contains split summaries. **(D)** Independent directional BA for five published methods and eight reference entries on GSE303036 stroma, Du bulk, and Sun bulk truth. Two pairs with identical predictions share rows, giving 11 displayed rows for 13 method entries. The merged rows represent Original frozen rule and the field-selected original/six-family references. GSE77938, ComBat–limma, MetaIntegrator, and DerSimonian–Laird identify their corresponding single-arm, batch-adjusted, and meta-analysis predictions; default denotes the default ComBat implementation, field denotes field-selected variants, and affine denotes affine variants. M1 is the fitted meta-regression reference with four control-source classes, living enucleation separate, and additive tissue-layer terms. The spatial, Du, and Sun columns all use its fitted postmortem whole-cornea/stroma route; Sun truth is living SMILE-derived stroma and retains that historical prediction route. All displayed direct M1 rows are fitted predictions. Each truth-direction strip retains 60 ordered equal-count rank bins, ordered by decreasing absolute truth log_2_FC; color is correct/n. The common pools contain 14,304 spatial genes (8,420 up/5,884 down), 3,427 Du entries (715/2,712), and 2,875 Sun entries (565/2,310). BA averages the two direction-specific recalls equally; equal strip widths do not imply equal entry counts. Bold rows identify reference anchors. This panel prints point estimates; spatial CIs elsewhere resample donors and bulk CIs resample published entries. Complete scores and paired comparisons are in Supplementary Table S13. KC, keratoconus; BA, balanced accuracy; M1, model 1; NE, not estimable; CI, confidence interval; SMILE, small-incision lenticule extraction; log_2_FC, log_2_ fold change.

**Supplementary Figure S2. Control source, spatial location, protein measurements, and infection comparisons resolve the tissue state. (A)** M3 class estimates and 95% CIs for ECM, IEG, inflammation, and TGF-β, with member-gene KC−control log_2_FC across nine arms. M3 omits the two living-enucleation arms and fits control provenance and tissue layer in the remaining seven arms. Circles, squares, and diamonds show postmortem whole-cornea/stroma, pathological whole-cornea/stroma, and healthy/refractive epithelial estimates. These are fitted class estimates. The four 20-gene × 9-arm matrices contain 720 positions, with 616 finite effects; hatching denotes missing effects. H/P/D denote the original healthy/refractive, pathological, and postmortem model classifications; epi/st denote epithelium and whole cornea/stroma. These matrix classifications retain the original assignments of living-enucleation arms, whereas M3 estimates omit those arms. Gene identities and row order are listed in the Supplementary Figure S2A source-data table. Definitions, tissue summaries, and class estimates are in Supplementary Tables S2, S6, and S7. **(B)** IEG and DC/immune maps for all 10,876 retained GSE303036 tissue spots from 13 donors (5 controls, 8 KC), with donor means below. Columns identify the same donors in both program maps; spots and sections are nested within donors. Faint outlines identify epithelial spots. Each program uses a symmetric signed-square-root color scale, sign(z)√(| z|/maximum |z|), shared between its maps and donor-mean strip; color-bar labels report the original z scores. A1 and A3 are rotated together for display while preserving within-section distances. Scores and donor-level effects are in Supplementary Table S9. **(C)** Unpooled mixed epithelial/stromal TMT proteomics from five KC and five control donors. Forest points and lines show KC−control class effects on a log_2_ scale and Welch 95% CIs. Small circles are donor class scores; larger diamonds are group means; filled and open symbols indicate KC and controls. The heatmap contains 93 source protein rows/accessions across ten channels: 9 ECM, 2 inflammation, and 82 immune-effector rows. Gray intensity shows log_2_ abundance. Same-symbol proteins are averaged within channel for the donor class score while source protein rows remain separate in the heatmap. Protein row/accession identities, order, and channel keys are listed in the Supplementary Figure S2C source-data table; N/K channel labels identify control/KC samples. This protein immune-effector set differs from the DC/immune transcript marker set. Supplementary Table S6 contains protein summaries and source-member records. **(D)** IEG, inflammation, and CollecTRI AP-1 scores for GSE241715 (7 normal, 4 KC, 5 viral, 5 bacterial/fungal samples), with six KC−infection differences in Hedges’ *g*. Open triangles and filled squares identify viral and bacterial/fungal comparisons. Intervals are 95% CIs incorporating shared-normal-control covariance. The IEG viral interval crosses zero (Δg, −0.453; 95% CI, −1.574 to 0.668); each comparison retains its own estimate. These samples recur in the control-substitution analysis. Supplementary Table S10 contains disease comparisons. KC, keratoconus; ECM, extracellular matrix; IEG, immediate-early genes; TGF-β, transforming growth factor-β; DC, dendritic cell; AP-1, activator protein-1; CollecTRI, Collection of Transcriptional Regulatory Interactions; CI, confidence interval; log_2_FC, log_2_ fold change; M3, control-provenance model 3 excluding living-enucleation controls; TMT, tandem mass tag.

**Supplementary Figure S3. Stimulation and loading distinguish global disease similarity from immediate-early responses. (A)** Human stromal-cell responses in GSE260476: four libraries each under SF, TGF-β1, and 10% serum conditions. The same SF libraries enter both treatment comparisons. The upper matrix shows six AP-1/IEG member-gene expression values per library; the lower matrix shows CollecTRI and DoRothEA ABC activity, member SSM, and member-rank scores. Each readout retains its own scale. Triangles and squares show serum−SF and TGF-β1−SF effects with HC3 95% CIs; shapes identify treatment. AP-1/IEG scores are sign-inverted toward the low-activity KC signature: positive Δ is KC-directed, so the negative serum CollecTRI change (−1.687) represents movement away from the low-activity state. Library is the recorded unit; between-condition donor pairing is unavailable. **(B)** Rabbit stromal-cell responses in GSE287915: eight libraries each under SF, FBS, and TGF-β1 conditions, with the same SF group reused. Gray points show library scores; dark circles, triangles, and squares show SF, FBS, and TGF-β1 means. Treatment effects and HC3 95% CIs share the corresponding row baseline. AP-1/IEG SSM and rank scores are sign-inverted toward low KC activity; ECM member scores instead weight each gene by its pooled KC−control expression direction. Positive Δ is directed toward the respective KC program. Readouts have separate scales. Human serum and rabbit FBS retain their source treatment definitions. Supplementary Table S11 contains the stimulus effects and inference; HC3 residual degrees of freedom are 6 in human and 14 in rabbit. **(C)** Global cosine similarity to pooled KC tissue and six-member AP-1 changes under acute human corneal-cell stretch (SRP648618, F−C and EF−C) or mouse back-skin expansion (GSE186773, E−NE at PI0, PI7, PI14, and PI21). C denotes static control; F and EF denote 0.1-Hz and 0.5-Hz stretch. The general human stretch protocol reports 15% strain for 6 hours at 0.5 Hz; the F arm’s amplitude and duration are not separately established. Dark marks and intervals show cosine with gene-bootstrap 95% CIs, using 12,456 shared genes per acute contrast and 14,266 per mouse contrast. Red human squares show RNA-sequencing model AP-1 effects with their model CIs; mouse circles show mean microarray log_2_ signal changes, for which source intervals are unavailable. Cosine and AP-1 change have separate axes, and the two biological systems retain separate comparisons. **(D)** Individual-library six-gene AP-1 mean log_2_ expression in the acute human C/F/EF groups, with three libraries per group. **(E)** AP-1 mean RMA-normalized log_2_ microarray signal in mouse expanded and nonexpanded skin, with one array per group at each PI day. Open and filled points identify NE and E; lines join same-period values. **(F)** Sixteen mouse E−NE log_2_FC values for FOS, EGR1, JUN, and JUNB over the four periods. Warm and cool colors indicate decreases and increases. This four-member display retains the members available in the listed-effects source; ATF3 and FOSL1 are absent. MMP1 is absent at all periods and is not assigned zero. **(G)** All 12,456 shared gene effects in human EF−C versus pooled KC−control. Gray cells count genes in a 40 × 40 grid using square-root count shading (251 occupied bins; 1–1,172 genes per occupied bin). Red diamonds identify ATF3, EGR1, FOS, FOSL1, JUN, and JUNB. Bins summarize gene records. Supplementary Table S12 and its associated data-release files index the experimental-model comparisons. AP-1, activator protein-1; IEG, immediate-early genes; ECM, extracellular matrix; KC, keratoconus; TGF-β1, transforming growth factor-β1; SF, serum-free; FBS, fetal bovine serum; HC3, heteroskedasticity-consistent standard errors of type 3; CI, confidence interval; SSM, standardized-mean score; rank, mean member rank among analyzable genes within each library; CollecTRI, Collection of Transcriptional Regulatory Interactions; DoRothEA, Discriminant Regulon Expression Analysis; ABC, confidence levels A–C; RMA, robust multi-array average; log_2_FC, log_2_ fold change; E/NE, expanded/nonexpanded skin; PI, days after full expander injection.

**Supplementary Figure S4. Gene membership, deletion analyses, structural-trait projection, and MR instruments resolve the genetic signal. (A)** Membership of collagen fibril organization, assembly, and formation (79, 61, and 90 genes; union, 127; three-way intersection, 28), with source-specific competitive MAGMA coefficients and gene Z statistics. The separate six-module display contains 96 distinct mapped genes and 114 memberships, with joint coefficients for Hardcastle, MVP European, FinnGen R13, CCT, CRF, and CRF⊥CCT. Each narrow column is an Entrez gene; the two gene universes have separate ordering and index scales. Gene order is provided in the Source Data. Gray membership cells identify included genes, and hatching identifies genes untested in a source. Public-pathway Z statistics are available for 124, 124, and 126 genes by disease source; all 96 mapped fixed-module genes have Z statistics in each source. Coefficients and gene Z statistics retain separate scales. Supplementary Tables S2 and S3 provide definitions and enrichment results. **(B)** Marginal ECM enrichment after each of 19 single-gene deletions and 14 physical-locus deletions in each of three disease sources (57 and 42 deletion analyses). Locus labels identify all genes removed together; outlined cells mark multigene loci. The separate Full cells show each source’s undeleted marginal ECM model. Color encodes log_10_[(0.05/6)/*P*]; zero marks the six-module Bonferroni threshold, and every deletion remains below that *P*-value threshold. Genes share loci and analyses share input data. Supplementary Table S3 contains the deletion results. **(C)** Joint structural-trait projections onto disease common-SNP heritability for right- and left-eye 3-mm and 6-mm measurement families. Squares and lines show the original R^2^ projections and 200-block jackknife 95% CIs; pale ticks show the 200 block-deletion estimates for each source–family combination (2,400 values). The final combinations include CCT, CRF, and five corneal measurement families. Eye, measurement-family, and disease-source estimates share inputs. Intervals retain their original endpoints, including values outside 0–1. Results are in Supplementary Table S5. **(D)** Multivariable CRF and CCT components for Hardcastle, MVP, and FinnGen. Each point is a harmonized source–SNP record (101, 124, and 120 per exposure), plotting the selected exposure’s effect against β_KC minus the fitted contribution of the other exposure. Lines through the origin use the corresponding no-intercept multiplicative random-effects IVW coefficient. The two exposure rows reuse each source’s instruments and have separate coordinate scales; within each exposure row, the y-axis scale is shared across disease sources. CRF ORs per mm Hg are 0.456 (95% CI, 0.358–0.582), 0.527 (0.405–0.684), and 0.392 (0.308–0.499); CCT ORs per µm are 0.997 (0.986–1.009), 0.993 (0.980–1.006), and 1.003 (0.991–1.015). Complete estimates are in Supplementary Table S4. KC, keratoconus; ECM, extracellular matrix; IEG, immediate-early genes; NRF2, nuclear factor erythroid 2-related factor 2; Corneal inflam., corneal inflammation; MAGMA, Multi-marker Analysis of GenoMic Annotation; MVP, Million Veteran Program; MR, Mendelian randomization; MVMR, multivariable Mendelian randomization; IVW, inverse-variance weighted; CRF, corneal resistance factor; CCT, central corneal thickness; CRF⊥CCT, genetically CCT-residualized CRF; SNP, single-nucleotide polymorphism; OR, odds ratio; CI, confidence interval; R/L, right/left eye.

**Supplementary Figure S5. Structural sharing extends from risk windows to credible components and corneal accessible elements. (A)** PP4 for 61 candidate windows across three disease sources and four trait/version channels: updated CCT (GCST90000654; Choquet 2020), CRF, CRF⊥CCT, and original CCT (GCST006366). Genome-ordered windows are split into 31 and 30 rows, with chromosome 10 continuing across the split. Cell color shows PP4 from 0 to 1; borders indicate PP4 ≥ 0.8, corner circles indicate SuSiE, and the black track identifies sharing in any of the three primary channels, excluding the original-CCT comparison channel. These channels yield 27 of 61 shared windows, including 17 of the original 40. Windows can overlap. Source-specific posteriors and methods are in Supplementary Table S5. **(B)** COL5A1 credible-set memberships for five shared component pairs, ordered by GRCh38 position. The 39 distinct variants contribute 81 component–role–variant PIP records. K and T identify KC and structural-trait credible-set roles. Teal cells show original PIP; hatched gray cells denote variants outside that role’s credible set. Outlines identify the three primary CCT L1/L1 pairs; their eight-variant union is bracketed. The Hardcastle, MVP, and FinnGen primary-component unions contain 5, 8, and 6 variants, respectively. Displayed PP4 values are FinnGen CCT L1/L1, 0.989; Hardcastle CCT L1/L1, 0.991; Hardcastle CCT L3/L3, 0.846; MVP CCT L1/L1, 0.985; and MVP CRF L1/L3, 0.863. L1 and L3 retain the fitted component identifiers. PP4 is the component-pair posterior; cell color represents variant PIP. **(C)** KC credible-set PIP fractions overlapping accessible peaks in 21 author-defined cell classes and a separate all-deposited-peaks aggregate. The shared-region field contains 17 × 22 values and the background field 40 × 22 values. The 17 regions are included in the 40, but each field retains its own signal inputs and aggregation: shared pairs are grouped together, whereas the background is averaged by source groups. Thus the same region–class key can have different values in the two fields, and columns have independent region ordering. The two highlighted stromal classes are selected displays: keratocytes have 1.88-fold enrichment (empirical *P* = 0.015), and stromal stem cells 1.94-fold enrichment (*P* = 0.013). All class results, including other enriched classes, are in Supplementary Table S5, block D. Empirical *P* values use 1,000 draws of 17 regions from the original 40; stored null intervals describe the permutation distribution. **(D)** Recorded accessible-peak matches across 199 locus–SNP columns from 12 loci and 21 cell classes. Slate cells identify at least one peak match: 683 occupied positions derive from 1,165 source rows, 373 signal–SNP identities, and 96 peaks. Twenty classes have matches; the red-blood-cell row has no recorded match. Empty cells denote absence of a recorded match. Block numbers identify the locus names below the matrix. The ALDH3A1 inset shows rs4646785 at GRCh38 chr17:19,748,589 (PIP, 0.955), within peak chr17:19,747,406–19,749,213, matched in corneal stromal keratocytes and stromal stem cells. The regional COL5A1 matches are distinct from the eight primary-component variants in B. Supplementary Table S5 summarizes regulatory context. H/M/F, Hardcastle/MVP European/FinnGen R13; MVP, Million Veteran Program; KC, keratoconus; CCT, central corneal thickness; CRF, corneal resistance factor; CRF* or CRF⊥CCT, genetically CCT-residualized CRF; PP4, posterior probability that two traits share a causal variant; PIP, posterior inclusion probability; SuSiE, sum of single effects; SNP, single-nucleotide polymorphism; GWAS, genome-wide association study; ATAC, assay for transposase-accessible chromatin; GRCh38, Genome Reference Consortium human genome build 38; C., corneal; Conj., conjunctival; L., limbal.

**Supplementary Figure S6. KC-derived cells retain feedback and matrix induction across culture formats without uniform reversal of tissue-low genes.** All panels use the same three KC donors (1206, 1207, and 1208) and 24 libraries across two formats and four treatments. Ligand contrasts share vehicle controls. The stimulus experiment contains no normal-donor arm. **(A)** DESeq2 model estimates of TGF-β1-induced SMAD7 expression relative to vehicle in 2D and bioprinted 3D culture: log_2_FC, 2.44 and 0.93; Benjamini–Hochberg-adjusted *P*, 1.61 × 10^−20^ and 0.00108, respectively. DESeq2 fits donor and format–treatment terms, with two-sided Wald inference. Thin paired lines separately show each donor’s 2D and 3D log_2_ induction ratio. Open and filled circles identify 2D and 3D. Individual ratios and fitted coefficients retain their separate axes. **(B)** GSEA for the format-by-stimulation contrast, (3D TGF-β1−3D vehicle) − (2D TGF-β1−2D vehicle). NABA_COLLAGENS has NES 1.919 and *q* = 0.0134; HALLMARK_MYOGENESIS has NES −1.367 and *q* = 0.0369. The *q* values are the original adjusted gene-set *P* values. Rug marks show Wald statistics for all finite-ranked members (35 and 152), rather than only leading-edge members. Below, the left display joins the 2D and 3D DESeq2 coefficients for nine predefined collagen-assembly and five contractile-myofibroblast genes. The right display joins, for each of the two modules, every donor’s TGF-β1-minus-control difference in mean log_2_(CPM+1) expression of the module members within 2D and within 3D, with D1/D2/D3 identifying donors 1206/1207/1208. The left and right displays use different quantities and axes, and neither corresponds to the count-based donor ratios in A. These predefined modules differ from the two GSEA sets. Filled gene symbols identify 3D; open symbols identify 2D. **(C)** 3D TGF-β1 responses of genes selected for low expression in four patient tissue comparisons, ordered by tissue rank. Color represents model log_2_FC with signed-square-root shading; labels give the original log_2_FC scale. Upper and lower short bars identify significant increases and decreases, respectively, at adjusted *P* < 0.05; unmarked tested cells are NS. Hatched gray cells have no model result. Up/down/tested-NS/no-model counts are 21/46/74/59 for GSE204791 stroma (200 members), 25/35/84/56 for PRJNA1184491 stroma (200), 29/20/73/4 for GSE303036 stroma (126), and 59/26/61/54 for GSE77938 (200). These are 726 gene–cohort memberships, with genes allowed to recur across sets. The blank extent after the 126-member set is outside its membership. Supplementary Table S11, blocks D–F, reports culture effects, GSEA, and member-direction counts. KC, keratoconus; 2D/3D, two-dimensional/three-dimensional culture; TGF-β1, transforming growth factor-β1; DESeq2, differential expression analysis for sequence count data, version 2; log_2_FC, log_2_ fold change; GSEA, gene set enrichment analysis; NES, normalized enrichment score; NS, not statistically significant; *q*, multiple-testing-adjusted gene-set *P* value.

**Tables**

Tables 1–3 are main tables; Supplementary Tables S1–S16 accompany the manuscript.

- **Table 1**. Human tissue cohorts and independent expression tests.
- **Table 2**. Control-dependent decisions and recovery with normal references.
- **Table 3**. Published operations and the identity of their measured endpoints.
- **Supplementary Table S1.** Complete data-source and analysis-unit inventory.
- **Supplementary Table S2.** Gene-set definitions, symbol resolution, and measurement coverage.
- **Supplementary Table S3.** Genetic pathway enrichment and contribution of collagen loci.
- **Supplementary Table S4.** Corneal biomechanics and keratoconus: Mendelian-randomization estimates.
- **Supplementary Table S5.** Shared genetic signals, credible variants, and corneal regulatory context.
- **Supplementary Table S6.** Tissue gene and program effects across all study arms.
- **Supplementary Table S7.** Control-source definitions and reference-specific expression estimates.
- **Supplementary Table S8.** Regulatory activity, conditional feedback, and within-donor collagen relationships.
- **Supplementary Table S9.** Spatial donor states, regional coverage, and immune readouts.
- **Supplementary Table S10.** Disease, within-cornea, clinical, and blood comparisons.
- **Supplementary Table S11.** Inducible programs, handling effects, and patient-derived culture responses.
- **Supplementary Table S12.** Fixed-case control substitution and model/candidate replay.
- **Supplementary Table S13.** Normal-reference selection and independent validation.
- **Supplementary Table S14.** Published decision premises, risk-direction concordance, and operation index.
- **Supplementary Table S15.** DNA methylation and m6A measurements in their assayed universes.
- **Supplementary Table S16.** Reusable data release and scientific result index.

## Code key for the supplementary tables

Module codes: FULL_ECM = ECM/proteoglycan; FULL_STRESS = stress/immediate-early; INFLAMMATION = general inflammation; CORNEAL_LOCAL_INFLAM = corneal-local inflammation; OXIDATIVE_STRESS_NRF2 = oxidative stress/NRF2; ATOPIC_SYSTEMIC = atopic/systemic; TGFB_SIGNALING and PURE_ATOPY_CANONICAL = the two attribution follow-up sets. Source/LD-context codes: HARDCASTLE_EUR = Hardcastle European; GCST90477661 and MVP_EUR = MVP European; AA_AFR and AA_EUR = MVP mixed-ancestry under AFR and EUR LD; FINNGEN_R13 = FinnGen release 13. Universe code: U-MHC-full = the unified gene universe, MHC region included. Public pathways use the source-specific MAGMA gene universes described in Supplementary Table S3.

