## Supplementary Methods for "Reinterpreting Keratoconus Gene Expression in Light of Inherited Stromal Weakness"

Numbered citations use the reference list of the main article; citations marked S1–S5 are listed under Supplementary References at the end of this document.

#### Study Design and Data Provenance

The core computational analysis reanalyzed public, deidentified data. The culture analysis used RNA-sequencing matrices from patient-derived bioprinted stroma,[85] as described under Patient-Derived Culture Transcriptomes. No new human samples were collected and no animals were used in this analysis. Ethical approval and participant consent were obtained by the source studies, and the rabbit series was derived by its depositors from commercially supplied rabbit eyes.[34] Control-tissue provenance and the definitions of corneal regions, tissue layers, and clinical measurements were obtained from the source publications. The dataset inventory and analysis-specific sample units are given in Supplementary Table S1.

#### Genetic Data Sources and Effect Scales

The three independent disease sources were two European keratoconus (KC) GWAS—Hardcastle et al. (GCST90013442; 2,116 cases and 24,626 controls)[3] and MVP European (GCST90477661; 573 cases and 450,622 controls)[16]—and the FinnGen R13 corneal-deformation endpoint (H7_CORNEALDEFORM; 921 cases and 471,319 controls).[17] The overlapping MVP mixed-ancestry GWAS (GCST90480060; 940 cases and 572,131 controls) shared its European participants with GCST90477661 and remained an ancestry and linkage-disequilibrium (LD) reference sensitivity analysis for the prespecified modules, using African and European reference panels.

The structural-trait GWAS were central corneal thickness (CCT; GCST006366; N = 17,803)[18] and corneal resistance factor (CRF; GCST90100568; N = 76,029)[19]. CRF was measured with the Ocular Response Analyzer, averaged across both eyes, and analyzed on its original millimeters-of-mercury scale. The source CRF GWAS excluded KC, other corneal diseases, and previous ocular surgery. The larger CCT meta-analysis (GCST90000654; N = 44,039)[20] was used for the primary whole-risk-region colocalization census, with GCST006366 analyzed alongside it. That meta-analysis included 32,292 European, 9,623 East Asian, 1,367 Hispanic/Latino, and 757 African American participants. The prespecified structural-trait module tests, CCT-residualized CRF construction, and Mendelian randomization (MR) used GCST006366 for CCT. CCT effects are in micrometers and CRF effects in millimeters of mercury.

#### Bulk Transcriptome Inputs and Control Provenance

Nine analytic arms — dataset-by-tissue case–control contrasts — were drawn from eight source studies. Six were Gene Expression Omnibus (GEO) or ArrayExpress series: GSE151631 (whole cornea),[10] GSE77938 (whole cornea),[14] GSE112155 (epithelium),[22] GSE241715 (whole cornea; also providing viral and bacterial/fungal keratitis arms),[23] GSE204791 (stromal and epithelial arms of one study),[24] and E-MEXP-2777 (whole cornea).[7] Two RNA-sequencing cohorts were reprocessed from the European Nucleotide Archive/Sequence Read Archive: PRJNA1184491 (stroma)[25] and PRJNA799648 (epithelium).[11] The reprocessed cohorts were quantified with salmon[72] against GENCODE v44,[73] imported with tximport,[74] and tested with DESeq2.[75] For the array series E-MEXP-2777, duplicate probes for one gene were represented by the highest-variance probe before limma empirical-Bayes testing; in the other deposited differential-expression tables, duplicate symbols were resolved to the row with the largest absolute test statistic. Control-tissue provenance was read from each source publication's Methods — neither repository records that field — and classified as postmortem donor, healthy/refractive, pathological, or living enucleation (Supplementary Table S1). The E-MEXP-2777 and GSE241715 control corneas came from eyes enucleated in living patients for melanoma. For the shared-platform comparisons in GSE241715, per-sample module and marker scores were the unweighted mean of gene-wise $z$ scores of $\log_2$ expression standardized across all 21 samples. The stress-module score used the 19 module genes detected in this cohort (*NPAS4* is absent); marker scores used keratocyte (*NNMT*, *KERA*, *ALDH3A1*, *ALDH1A1*, *LUM*, *DCN*, *CD34*, *PTGDS*), immune-infiltration (*PTPRC*, *S100A8*, *S100A9*, *MMP9*, *CXCL8*, *ITGAM*, *LYZ*), and myofibroblast (*ACTA2*, *TAGLN*, *COL1A1*, *FN1*) sets. Group effects were disease-group mean minus control mean, and differential-expression burden was genes at Benjamini–Hochberg (BH)-adjusted $P < 0.05$ over genes tested.

These nine-arm gene-effect analyses used the regional aggregation and study-specific case–control definitions of each study. The donor-score analysis used the donor counts and within-donor aggregation specified below; the disease-comparison and regional analyses used their separately stated sample definitions.

#### Perturbation-Series Inputs

Two public corneal stromal perturbation datasets were used. GSE260476 profiles human corneal fibroblast states (serum-free quiescent, 10% serum proliferative, and transforming growth factor-β (TGF-β) myofibroblast; 4 independent libraries per state, 72 h).[32,33] States were assigned from the sample-level GEO treatment fields (A libraries untreated/serum-free, B libraries TGF-β1-treated, C libraries 10% serum-treated); the associated proliferation paper[33] labels A as proliferating and C as quiescent, the reverse of the GEO fields, and we retained the GEO mapping, which agrees with the markedly higher cell-cycle transcript counts in C. The deposit provides DESeq2-normalized gene counts, analyzed as $\log_2(\text{count} + 1)$ after requiring a normalized count ≥ 1 in at least four of the twelve libraries and nonzero variance; no cross-condition donor map is available, so libraries are the inference units. GSE287915 profiles rabbit corneal stroma (serum-free, fetal bovine serum [FBS], and TGF-β1; 8 libraries per group).[34] Rabbit counts were mapped to unique one-to-one human orthologs of BioMart confidence 1, normalized as counts per million (CPM) against the original library totals, transformed as $\log_2(\text{CPM} + 1)$, and retained at CPM ≥ 1 in at least eight libraries with nonzero variance; 15 of the 20 extracellular-matrix (ECM) genes and five of the six activator protein-1 (AP-1)/immediate-early gene (IEG) genes (*JUNB* unavailable) were analyzable. The public matrix carries no animal identifiers, so libraries are the inference units.

#### Within-Dataset Contrast-Definition Analysis

GSE142538 (Fuchs endothelial corneal dystrophy [FECD] endothelium)[27] also provided a within-dataset demonstration of contrast definition. Three groups — pre-symptomatic repeat-expansion carriers, advanced repeat-expansion FECD, and advanced non-expansion FECD — were each compared with the same controls; the first comparison draws both sides from the same tissue source, the other two couple disease state with tissue source. Differential-expression burden is the number of genes at BH-adjusted $P < 0.05$ over the number tested in each contrast. This contrast-definition analysis is reported separately from the FECD module-score synthesis described below (Supplementary Table S10, block E).

### Additional Molecular and Spatial Datasets

Additional datasets comprised the reduced-representation bisulfite sequencing (RRBS) methylome GSE107189 (5 KC, 5 control),[35] $N^6$-methyladenosine (m6A) epitranscriptome GSE181618 (5, 5; Arraystar array, within-sample immunoprecipitate [IP]/supernatant contrast), and two Visium spatial cohorts, GSE303036 (8 KC, 5 control)[8] and GSE285060 (5, 3).[29]

### Prespecified Gene Modules and Testing Families

Six mechanism modules were prespecified before testing, with membership listed in Supplementary Table S2: ECM/proteoglycan, stress/immediate-early, general inflammation, corneal-local inflammation, oxidative stress/nuclear factor erythroid 2-related factor 2 (NRF2), and a literature-derived atopic/systemic set. The atopic/systemic set was assembled from genes prioritized in published KC germline analyses, including *SMAD3* and *KLF5*. These modules operationalize the nominated KC mechanisms. Inflammation enters as two sets (general and corneal-local), type-2 atopy as the literature-derived set, and the keratocyte stress program as the stress/immediate-early module; TGF-β signaling is evaluated separately as an attribution follow-up (below). The prespecified definitions hold 20 genes per module and 19 in the atopic/systemic set, 119 memberships in all; the tissue and methylome layers use these full definitions, and each layer reports the members it covers. Germline member counts are the subset analyzable by Multi-marker Analysis of GenoMic Annotation (MAGMA) in the unified gene universe (below), which admits a gene only where all five source contexts return a result. Five memberships fail that requirement, all of them outside the autosomes: *BGN*, *TIMP1* (in both inflammation sets) and *G6PD* lie on the X chromosome and are returned by FinnGen alone, and the pseudoautosomal *CRLF2* is returned by no source. The ECM module therefore enters MAGMA with 19 genes and the atopic set with 18 after alias resolution (*PCNXL3* maps under its current symbol *PCNX3*; Supplementary Table S2). Joint deletion of *SMAD3* and *KLF5* leaves 16 genes; the literature-prioritized subset holds 6. These six sets form the prespecified family of the joint conditional model. Two additional mechanism-layer sets defined independently of any KC GWAS result — canonical type-2 atopy and TGF-β signaling (with and without *SMAD3*) — were evaluated marginally as attribution follow-ups (Supplementary Table S3, blocks C and F). ECM shares no member with any other module; the only overlaps are inflammation with corneal-local inflammation (17 mapped genes, 18 in the prespecified definitions, the difference being *TIMP1*) and inflammation with the atopic set (*IL17A*).

The prespecified module analyses used a single alias-resolved gene universe (U-MHC-full), constructed by mapping each input symbol to its HUGO Gene Nomenclature Committee–approved symbol[36] and corresponding Entrez identifier while retaining major histocompatibility complex (MHC)-region genes. Mapped module membership was restricted to Entrez identifiers with MAGMA gene results in all five source/LD contexts, with duplicate set–identifier pairs removed. All marginal and joint prespecified-module results were computed in this universe.

Germline enrichment used MAGMA v1.10[37] competitive gene-set tests against the genomic background, conditioning on gene size, gene density, inverse minor-allele count, and their logs. Gene windows were 35 kb upstream / 10 kb downstream on NCBI37.3 gene locations, with 1000 Genomes[38] LD panels (EUR for the European analyses and FinnGen; ancestry-aware AFR primary and EUR sensitivity for the mixed meta-analysis). Competitive tests were used because they ask whether a set carries more association than matched background genes, the question a mechanism claim requires.[39] Marginal models tested each module separately; the joint conditional model entered all six module-membership indicators simultaneously with the same gene-level covariates. Each GWAS was corrected for the number of sets tested in it: α = 0.05/6 = $8.33 \times 10^{-3}$ for the six-module marginal and joint tests and α = 0.05/5 = 0.01 for the five-set attribution ladder. For the endophenotype extension, the same module definitions, annotation settings, covariates, and marginal and six-module joint models were applied to CCT, CRF, and CRF⊥CCT (Genetic Data Sources and Effect Scales), with enrichment assessed at α = 0.05/6 within each channel and model. Ordering used a three-module joint model (ECM, stress/immediate-early, and NRF2 indicators, with each contrast obtained by sum-and-difference reparameterization) within each channel, without an intersection–union test across channels, because the three channels are related rather than independent replications. ECM deletions were evaluated marginally and in joint models that replaced the intact ECM set with one deletion set while retaining the other five modules. U-MHC-full was primary and the MHC-excluded universe a sensitivity analysis. Deletion analyses are evaluated at the same per-GWAS threshold.

#### Germline Ordering and Robustness

Two germline ordering contrasts were prespecified — ECM − AP-1/stress and ECM − NRF2, each the difference of joint module coefficients (Δβ) with its standard error from the joint fit — and evaluated as intersection–union tests (IUT) [S1, S2]. Each contrast must hold one-sided in both independent European sources, giving a contrast-level two-source IUT *P* (the maximum of its two source *P* values); the family-level ordering IUT *P* is the maximum across the four components of both contrasts. All four component *P* values are reported, and the component selection is by layer, not by result. Deletion analyses tested the persistence of the marginal ECM enrichment in the unified universe, not a re-estimation of the ordering. In KC, leave-one-gene-out comprised 95 deletions across five source/LD contexts: 76 across the other four backgrounds and 19 in FinnGen; for leave-one-physical-locus-out, the 19 ECM genes were merged into 14 loci by chromosomal adjacency with an inter-gene gap ≤ 1 Mb. Calibration covered the two contrasts in four contexts — Hardcastle, MVP European, and the mixed-ancestry analysis under AFR and EUR LD — with 1,000 matched random gene sets per cell (seed 20260824): 19-gene sets were drawn without replacement from genes outside all six modules, preserving the ECM module's counts in 5 × 5 × 5 quantile strata of log gene length, log single-nucleotide polymorphism (SNP) density, and log(NSNP/NPARAM), the ratio of SNP count to the MAGMA gene-parameter count; the same 1,000 sets were evaluated in all four contexts. Empirical *P* values are two-sided with add-one correction (twice the smaller tail).

#### Bidirectional Mendelian Randomization

Bidirectional two-sample MR evaluated CRF and CCT against each of the three KC sources, giving 12 exposure–outcome directions. Inverse-variance weighting (IVW) was the main estimator. MR-Egger[43], weighted-median[44], weighted-mode[45], and MR-PRESSO estimates[46] were obtained for the same directions using TwoSampleMR and MRPRESSO.[42] The weighted-mode analysis used 1,000 bootstrap samples, phi = 1, and a fixed seed. MR-PRESSO used 10,000 simulations and SignifThreshold = 0.05. Instrument strength was summarized by the mean and minimum F statistic, calculated for each instrument as $(\beta\_exposure/SE\_exposure)^2$.

The 95% confidence intervals for TwoSampleMR estimators used the software's normal approximation, while MR-Egger and weighted-mode *P* values used its t tests. MR-PRESSO confidence intervals used the t distribution with the fitted regression's residual degrees of freedom. Cochran Q, the MR-Egger intercept, and the MR-PRESSO global and outlier tests were reported with the corresponding instrument counts. Estimates that could not be obtained with the available number of instruments were retained as unavailable. The full method-specific results and diagnostic statistics are provided in Supplementary Table S4 (blocks A through E_variants).

Steiger analyses used a KC prevalence of 0.00138, liability-scale correlations for KC, and effective sample sizes. The overall direction test, the number of variants supporting the specified direction, the variant-level prevalence sensitivity results, and estimates after Steiger filtering were reported separately. For the KC-locus exclusion analysis, all variants associated with KC at $P < 5 \times 10^{-8}$ in any of the three disease sources were expanded by ±500 kb on the standardized GRCh38 coordinates. CRF and CCT instruments inside these intervals were removed, and the IVW model was refitted.

Forward effects are reported as odds ratios for KC per 1 mm Hg higher CRF or 1 µm higher CCT. Reverse effects are changes in CRF or CCT per unit increase in KC genetic log odds. These scales were preserved in plots and tables.

#### CAUSE Model Comparisons

CAUSE compared causal and sharing models in all 12 directions.[47] The four Hardcastle directions and four directions each for MVP European and FinnGen were fitted with the same settings. Nuisance parameters were estimated from up to 1,000,000 randomly sampled variants using seed 20260821, mixSQP, null_wt = 10, and n_iter = 20. Model fitting used the CEU LDshrink reference with allele frequency ≥ 0.05, an exposure-association threshold of $P < 0.001$, and LD pruning at $r^2$ = 0.1.

The prior on the proportion of variants with a shared effect was Beta(1,10), with max_q = 1. The initial grids contained 21 values each for γ and η and 11 for $q$, giving a 21 × 11 sharing grid and a 21 × 21 × 11 causal grid. sigma_g used the default estimation procedure, and force was FALSE. Results are reported as γ with a 95% credible interval, and as exp(γ) for forward odds ratios. The expected log pointwise predictive-density difference was calculated as sharing minus causal. The accompanying one-sided $P$ value tests whether the causal model fits better than the sharing model; it is not a test of γ against zero. All comparisons are included in Supplementary Table S4.

#### Multivariable Mendelian Randomization

CRF and CCT were entered together using the union of their instruments.[48] This set was reclumped with European LD at $P < 5 \times 10^{-8}$, $r^2 = 0.001$, and a 10,000-kb window. After allele harmonization across CRF, CCT, and each KC outcome, no-intercept multiplicative random-effects IVW was fitted with MendelianRandomization::mr_mvivw, using model = "random" and distribution = "normal".[49] The fitted effects, 95% confidence intervals, $P$ values, instrument counts, and Cochran Q were reported for each disease source. CRF effects remained on the per-mm Hg scale and CCT effects on the per-µm scale (Supplementary Table S4).

#### Whole-Risk-Region Colocalization Census

Genome-wide significant KC variants were selected independently within each disease source at $P < 5 \times 10^{-8}$. LD clumping used 1000 Genomes European LD[38], a ±500-kb window, and $r^2 < 0.1$ between lead variants. Each lead defined a ±500-kb physical interval. Identical or overlapping intervals were merged across sources, producing 40 nonoverlapping risk regions. Six regions overlapped the 14 prespecified ECM intervals and 34 did not. Two lead variants absent from the LD panel, rs116576225 in MVP and rs1704547735 in FinnGen, were retained as physical risk regions after checking their rsID positions; reference-matched variants within their intervals were used for colocalization. Gene labels were positional annotations.

The primary census paired each KC source with the larger CCT study (GCST90000654; N = 44,039), CRF (GCST90100568; N = 76,029), and the unfiltered CCT-residualized CRF summary statistics (N = 17,803). The CCT study (GCST006366; N = 17,803) was analyzed in parallel. For the larger CCT study, β and two-sided *P* values were available; standard errors were reconstructed as $|\beta/\Phi^{-1}(1-P/2)|$ and allele frequencies were taken from the LD reference. Its GRCh38 summary statistics were matched to the GRCh37 reference by rsID and allele identity.

Each KC–trait comparison used its own pairwise intersection of aligned variants. Variants required reference minor-allele frequency ≥ 0.01, reference missingness ≤ 0.05, and an effect-allele-frequency discrepancy ≤ 0.20 against the reference; Hardcastle was exempt from the effect-allele-frequency check. A comparison required at least 100 common variants and 5% reference coverage. SuSiE fits[50],[51] used susieR 0.12.35 through coloc::runsusie with package-default fit settings, R 4.5.0, seed 1, and 95% credible sets. For binary KC outcomes, SuSiE used effective sample size, 4/(1/N_cases + 1/N_controls); quantitative traits used their GWAS sample sizes.

Every available signal-component pair was evaluated with coloc.susie in coloc 5.2.3[52], using p1 = p2 = $10^{-4}$ and p12 = $5 \times 10^{-6}$. When credible sets or usable signal pairs were absent, coloc.abf used the same priors[53] and the recorded total sample size and case fraction for the binary phenotype. All PP0–PP4 values were retained. PP4 represents a shared causal variant and PP3 distinct causal variants. A region–source–trait comparison was called shared when any valid signal pair, or its ABF analysis, reached PP4 ≥ 0.8. A high PP3 from another pair did not replace the posterior of a shared pair.

The region-level numerator counted each physical region once when sharing was detected in any disease source and any primary structural channel. Source-specific proportions used the significant risk regions belonging to that source as the denominator. Wilson 95% intervals summarized the region-level proportions. The primary matrix contains 40 regions × 3 KC sources × 3 structural channels = 360 comparisons; inclusion of the GCST006366 CCT analysis gives 480 comparisons. The main figure displays the largest valid PP4 across the three KC sources for each region and primary channel. Every displayed maximum retains its originating source, method, trait version, signal pair, and the PP3 from that same pair. The complete source-specific census and all signal pairs are provided in Supplementary Table S5 and Supplementary Fig. S5A.

The 14-locus ECM analysis used its own intervals, aligned variants, and SuSiE fits. Its 110 signal-pair posteriors and five locus–trait combinations were reproduced separately from the 40-region census. Its 126 locus–source–trait cells were also completed with ABF when the SuSiE comparison was unavailable. The two analyses carry separate input-version labels. Signal-resolved *FNDC3B* comparisons are reported with their KC and structural-trait component identities; regional ABF results are recorded under their own method.

**Component-Resolved COL5A1 Sharing**

The component analysis used the saved SuSiE fits, trait-specific credible sets, posterior inclusion probabilities, and coloc posteriors. Shared components used the PP4 ≥ 0.8 criterion. Disease and structural-trait credible-set membership was reported separately for every source–trait–component pair. No cross-trait PIP was constructed. The primary CCT pair was evaluated separately from the additional Hardcastle CCT and MVP CRF pairs. Variant matching used genome build, chromosome, position, and alleles; mapped GRCh38 positions were intersected with the adult corneal ATAC peaks and stromal-class-enriched peak sets. Total accessible peaks and class-enriched peaks remained distinct annotations.

Reporter-assay constructs were taken from a doctoral thesis on COL5A1 regulation.[54] Each construct's primer pair was searched against the complete GRCh38 reference; each of the four constructs matched exactly one same-chromosome, opposite-orientation pair 700–1,300 bp apart, giving amplicon lengths of 840, 938, 924, and 957 bp for regions 1–4. Variant posteriors were recomputed from the saved SuSiE fits with the coloc.susie call (coloc 5.2.3, susieR 0.12.35; defaults and priors unchanged); recomputed and stored PP4 agreed to within $4.4 \times 10^{-16}$ for all five shared signal pairs. The variant-level posterior conditional on a shared causal variant was summed over the variants inside a construct separately for each signal pair, without averaging across pairs and without a new credible-set threshold. Alleles at each construct variant were phased against the CCT-lowering lead variant of that construct by two-locus expectation–maximization haplotype-frequency estimation in the 1000 Genomes European panel (503 samples; genotypes lifted to GRCh38). The thesis reported alleles only at the lead variant of each construct, so alleles at the other variants are linkage-disequilibrium inferences rather than sequencing results. rs1536484 is palindromic, and its match to the source-strand KC risk allele was not assigned. Reporter activities are the values reported in the thesis (Wilcoxon signed-rank tests).

**CCT-Residualized Corneal Resistance**

CRF⊥CCT was constructed with genomic structural equation modeling[21] as the genetic residual of CRF on CCT. Signed Z statistics were reconstructed from two-sided *P* values and aligned effect directions. Standardized per-variant effects and standard errors were computed as $Z/\sqrt{[N \times 2f(1-f)]}$ and $1/\sqrt{[N \times 2f(1-f)]}$, respectively, on a common allele-frequency scale. The genetic CRF-on-CCT regression coefficient was 0.672 (SE, 0.074). Residual effects were $\beta\_CRF - 0.672 \times \beta\_CCT$, with sampling covariance, including sample overlap, propagated to their standard errors. The unfiltered residual summary statistics were used in the 40-region census.

**Public-Pathway Scan**

The public-pathway universe used MSigDB 2026.1.Hs[40] Reactome, Gene Ontology Biological Process, KEGG MEDICUS, and the 10 canonical NABA sets. Sets were merged by name and duplicate Entrez members were removed. Size filtering used each set's complete Entrez membership before intersection with a source's MAGMA gene universe. The main 10–500-member universe contained 7,132 pathways; the 10–1,000-member extension contained 7,310.

MAGMA v1.10[37] competitive single-set tests were run separately in Hardcastle, MVP European, and FinnGen. The analysis used each source's gene-level results and MAGMA default covariates and truncation. Hardcastle and MVP used their source-specific gene analyses, while FinnGen used its MHC-inclusive analysis. The single-set fits were obtained for the 10–1,000-member universe, and its 10–500-member subset was selected by set name. Benjamini–Hochberg correction[41] was applied separately within each source and size range. Discovery required a positive competitive coefficient and $q < 0.05$ in Hardcastle; independent replication required a positive coefficient and $P < 0.05$ in MVP European or FinnGen. Overlapping replicated collagen pathways were grouped descriptively as one collagen-assembly family, with every pathway estimate retained (Supplementary Table S3; Fig. 3A).

**Public-Pathway Tissue Estimates**

Public pathways were evaluated against the same unshrunk DESeq2[75] or limma gene effects used for the prespecified tissue sets. Duplicate symbols were represented by the row with the largest absolute test statistic, and the established CCN1/CYR61, CCN2/CTGF, and CXCL8/IL8 mappings were applied. Each gene contributed at most one record per arm. Finite effects with positive sampling variances were analyzed by a multilevel random-effects model with random intercepts for arm and for gene nested within arm: metafor::rma.mv(yi = log2FC, V = lfcSE$^2$, random = ~1|arm/gene, method = "REML").[60] The pooled intercept, 95% confidence interval, and two-sided Wald *P* value summarize the pathway's expression difference across the nine analytic arms.

For computational efficiency, large-set fits used grouped inversion of the same covariance matrix, diag(lfcSE$^2$ + $\sigma^2$_gene) + $\sigma^2$_arm Z_arm Z_arm$^T$, to evaluate the restricted maximum-likelihood objective and obtain the generalized least-squares estimate. Direct rma.mv fits were retained for the prespecified comparison sets and the three replicated genetic pathways. Tissue overlap with replicated genetic enrichment was defined by a two-sided pooled $P < 0.05$. The pathway-level tissue BH *q* values were also reported separately for each size range and tissue context. Record counts refer to gene–arm observations, with arm and mapped-gene counts recorded separately.

Whole-cornea/stromal and epithelial estimates used the same restricted maximum-likelihood model. Provenance estimates used the M3 model, which omits E-MEXP-2777 and GSE241715 and fits the remaining seven arms with provenance and tissue-layer moderators by maximum likelihood. Predictions were reported for the four observed provenance-by-layer combinations: postmortem-donor whole cornea/stroma, pathological whole cornea/stroma, pathological epithelium, and healthy/refractive epithelium. The complete public-pathway results and model-status fields are provided in Supplementary Table S3. The primary cross-layer summary uses the nine-arm pooled estimate.

**Pooled Gene-Level Tissue Expression and Control-Provenance Meta-Regression**

Gene-level $\log_2$ fold-changes ($\log_2$FC; maximum-likelihood DESeq2 or limma estimates without shrinkage) and sampling variances of the member genes of each of the eight sets were analyzed by multilevel models across the nine arms (metafor rma.mv),[60] with random intercepts for arm and for gene within arm. The pooled tissue change of a set is the intercept of the model without moderators (restricted maximum likelihood), reported with its 95% CI and two-sided Wald *P*; eight leave-one-source-study-out refits (both GSE204791 arms removed together) record sign retention. The same records entered the provenance models. Provenance-class estimates and their likelihood-ratio comparison came from a maximum-likelihood model with provenance and tissue layer as moderators (whole/stroma as reference), and each class is reported as the model prediction in the corneal layer where its arms were measured (in M1 and M3 the two healthy/refractive arms are epithelial, so that class is reported at the epithelial level; the pathological class has one whole-cornea and one epithelial arm and is reported at whole cornea/stroma; predictions at both layers are in Supplementary Table S7, block B); the exact permutation tests and the leave-one-source-out predictions below used a restricted-maximum-likelihood model with provenance as the only moderator. Four provenance-classification models were used: M0 (three classes, with the living-enucleation arms E-MEXP-2777 and GSE241715 assigned to the postmortem-donor and healthy/refractive classes), M1 (four classes, living enucleation separate), M2 (both enucleation arms folded into healthy/refractive), and M3 (enucleation arms dropped; seven arms). Provenance moderation was tested by exact conditional permutation of provenance labels at source-study level (the GSE204791 mixed block held fixed; 140/630/105/30 valid assignments under M0–M3). Out-of-sample consistency used leave-one-source-out sign prediction of the held-out arm's ECM effect.

**Donor-Level Module Scores and Transcription-Factor Activity**

Donor-level analyses included nine tissue arms from eight studies: GSE151631 (19 KC, 7 controls)[10], GSE77938 (25, 25)[14], GSE241715 (4, 7)[23], GSE112155 (10, 10)[22], GSE204791 stroma (8, 7) and epithelium (8, 8)[24], E-MEXP-2777 (10, 10)[7], PRJNA799648 epithelium (7, 7)[11], and PRJNA1184491 (10, 8)[25]. The analysis comprised 182 unique participants, 93 with KC and 89 controls. Eight KC participants in GSE204791 contributed both epithelial and stromal samples, producing 190 donor–arm observations; the epithelial and stromal control groups comprised different participants. Whole-cornea and stromal analyses included six arms with 76 KC and 64 control observations, and epithelial analyses included three arms with 25 KC and 25 control observations.

Expression processing followed that of the four-cohort analysis, including duplicate-symbol resolution and the cohort-specific $\log_2$(CPM + 1) or $\log_2$(x + 1) transformation. GSE204791 used normexp background correction with an offset of 16, quantile normalization within each tissue arm, and averaging of replicate probes with limma::avereps. E-MEXP-2777 used the archived $\log_2$ expression matrix, GPL5188 annotation, and the highest-variance probe for each gene symbol. The two PRJNA studies used the Salmon quantification files[72] and GENCODE v44 transcript-to-gene mapping[73]. In PRJNA1184491, each regional expression profile was transformed as $\log_2$(CPM + 1), and the central and peripheral values were averaged within each donor before scoring. This donor analysis included all 10 KC donors; the six-case subset corresponding to the stromal expression analysis was also evaluated.

Each gene was standardized within its study and tissue arm across KC and control donors, and each module score was the unweighted mean of the available member-gene z scores. The eight gene sets were unchanged from the primary analysis. Gene aliases were resolved with the mappings for CCN1/CYR61, CCN2/CTGF, and CXCL8/IL8. Module membership and gene coverage are provided in Supplementary Table S2. SMAD3 target output was scored in the same way from the 183 CollecTRI SMAD3 targets (feedback genes excluded), using the signed CollecTRI weights. For the shared-gene comparison, the genes each output set shares with the stress/immediate-early score (CCN2 for the TGF-β pathway set; CCN1, CCN2, EGR2, FOS, JUN, SRF, and ZFP36 for the SMAD3 targets) were removed, and the arm effects were pooled as for the other modules (Supplementary Table S6, block K).

Signed regulons were retained from CollecTRI[61] and DoRothEA confidence levels A–C[62]. DoRothEA edges were weighted 1, 1/2, and 1/3 for confidence levels A, B, and C, respectively, retaining the highest-confidence edge for each factor–target pair. The exclusion lists removed 18 ECM and 15 stress/immediate-early targets, including the CYR61 and CTGF aliases of CCN1 and CCN2. The seven module members outside those lists—COL5A2, COL12A1, BTG2, EGR2, IER2, NPAS4, and ZFP36—remained available as targets. Activities were estimated by the decoupleR univariate linear model[63] with at least five available targets per factor. FOS, JUN, JUNB, FOSL1, ATF3, and EGR1 activities were standardized within each study arm and averaged to give the AP-1/IEG composite. The six factors were fixed after the whole-network analysis and used in every analysis.

For each readout, arm-specific Hedges' $g$[64] quantified the small-sample-corrected KC−control mean difference relative to the pooled within-group standard deviation. Effect estimates and sampling variances were synthesized using restricted maximum-likelihood random-effects meta-analysis with metafor::rma.[60] We report pooled $g$, 95% confidence intervals, two-sided Wald $P$ values, and $I^2$ for all nine arms and separately for the whole-cornea/stromal and epithelial strata. Sensitivity analyses restricted PRJNA1184491 to six KC and eight controls and retained only one of the two GSE204791 arms at a time (Supplementary Table S6).

#### Four-Cohort Regulatory-Activity Analyses

The four-cohort regulatory-activity analyses included 107 participants: 19 KC and 7 controls in GSE151631, 25 and 25 in GSE77938, 4 and 7 in GSE241715, and 10 and 10 in GSE112155. The signed regulons, target exclusions, six-factor definition, and within-cohort activity standardization were those described above. These analyses are reported under their own cohort identities and are distinct from the nine-arm donor meta-analysis.

Within each cohort, the 725 transcription factors (TF) estimable in all four cohorts were ranked by CollecTRI Hedges' $g$, most negative first; the cross-cohort rank product is the geometric mean of these within-cohort ranks, each scaled by the 725 shared factors, and family location was tested by a one-sided Wilcoxon rank test of the six family ranks against the 719 background ranks (Supplementary Table S6, blocks I and I_test). Donor-level inference used per-cohort Hedges' $g$[64] pooled by random-effects meta-analysis (restricted maximum likelihood), with 95% confidence intervals, $I^2$, and leave-one-cohort-out analysis. A cohort-stratified permutation test evaluated the diagnosis coefficient from composite ~ diagnosis + cohort (100,000 two-sided within-cohort label permutations, seed 20260806); when no permutation reached the observed value, the add-one Monte Carlo $P$ value, 1/(permutations + 1), was reported (Supplementary Table S6, block J) [S3]. A network-by-estimator grid added the non-nested Transcriptional Regulatory Relationships Unraveled by Sentence-based Text mining (TRRUST) network [S4] (four family members estimable at regulon size ≥ 5 in all cohorts) and a weighted-mean estimator (Supplementary Table S8). Three objects carry the AP-1 label, each under its own definition: the 20-gene stress/immediate-early module (gene-set and arm-level tests), the six-factor regulon composite of ATF3, EGR1, FOS, FOSL1, JUN, and JUNB (donor-level activity), and the four-factor TRRUST composite (ATF3, EGR1, FOS, JUN).

The ranking, network-by-estimator, and permutation analyses are reported in Supplementary Table S6, blocks F–J. The 182-participant estimates occupy separately labeled sections of Supplementary Table S6.

#### Spatial Module and Marker Scores

The Visium analyses used GSE303036 (8 KC and 5 control donors)[8] and GSE285060 (5 KC and 3 control donors)[29]. Starting from the archived matrices, spots were retained at ≥500 unique molecular identifiers, ≥200 detected genes, and ≤20% mitochondrial transcripts. GSE303036 additionally used the archived in-tissue mask. Each gene was standardized across the retained spots within its cohort. A spot-level module score was the unweighted mean of the available member-gene z scores, and spots were then averaged within each donor to give the inference unit.

The stress/immediate-early, general inflammation, corneal-local inflammation, TGF-β, and ECM scores were evaluated with the fixed 20-gene definitions. Marker scores comprised the five dendritic-cell markers CD1C, CLEC10A, FCER1A, CD207, and CD1A; the three immune markers HLA-DRA, PTPRC, and CD74; and their combined eight-marker set. Analyses used members represented in every donor matrix within the cohort. All 20 genes were available for each of the five modules in both cohorts. GSE285060 lacked HLA-DRA, so its combined and immune scores comprised seven and two detected members, respectively. A gene absent from a donor's input feature list was recorded as unavailable rather than assigned zero expression (Supplementary Table S9).

The reported spatial effect was the KC−control difference in donor-mean score, Δz. Two-sided exact label-permutation tests enumerated all 1,287 allocations in GSE303036 and all 56 allocations in GSE285060. The accompanying 95% confidence intervals were t intervals from the corresponding unadjusted donor-level linear model. Marker-annotated stromal and epithelial domains in GSE303036 were joined to retained spots by GSM identifier and barcode, and donor-level scores were recomputed within each domain. The GSE285060 analysis used whole-section scores. Keratocyte-composition analyses[65] are reported alongside the module results (Supplementary Table S9, block D).

The keratocyte-composition analysis used robust cell-type decomposition (RCTD) to estimate keratocyte composition in both Visium cohorts.[65] Donor-level stress/immediate-early differences were evaluated in unadjusted models, models with a keratocyte-composition term, and analyses restricted to the lowest keratocyte-composition quintile of spots. These analyses used exact donor-level permutation tests and are reported alongside the module and marker results (Supplementary Table S9, block D).

**Comparisons With Infectious Keratitis and FECD**

Disease comparisons used bacterial and fungal keratitis in GSE58291 and GSE241715[23],[26], viral keratitis in GSE241715, and FECD in GSE142538 and GSE171830.[27],[28] GSE58291 contributed 7 bacterial cases, 8 fungal cases, and 12 controls after the source study's processed-data exclusions. Bacterial and fungal cases were combined for the principal infectious-keratitis synthesis and retained as separate groups in the per-study table. GSE241715 contributed 4 KC, 5 viral, 5 bacterial/fungal, and 7 control samples, with corneas obtained from living enucleation for melanoma as the shared controls. GSE142538 contributed 6 late repeat-expansion FECD cases and 9 postmortem-donor controls to the principal FECD synthesis; its non-repeat and presymptomatic groups were reported separately. In GSE171830, the two technical measurements of each specimen were averaged before analysis, giving three FECD pools, each containing three donors, and three individual controls. The FECD synthesis therefore comprised nine case statistical units and 12 control units (Supplementary Table S10).

For the disease comparisons, every gene was centered and divided by its sample standard deviation across all samples of its source study. Module scores were unweighted means of available genes; unexpressed or zero-variance genes did not contribute. The fixed modules and alias-resolution rules were applied. The stress/immediate-early module and the H002_STRESS_IMMEDIATE_EARLY set (Supplementary Tables S10 and S14) contained the same 20 genes and were treated as one readout. CollecTRI AP-1 activities used the signed regulons, target exclusions, and six-factor composite; each factor's activity was standardized across the full study panel before averaging.

RNA-sequencing count matrices used $\log_2$(CPM + 1). GSE241715 used the source study's normalized linear expression values transformed as $\log_2$(x + 1), retaining genes with $\log_2$ expression >1 in at least four samples. GSE58291 used the deposited $\log_2$ signals normalized across its array panel. GSE171830 technical measurements were averaged within pool or individual before gene-symbol mapping. Where multiple probes mapped to a symbol, the highest-variance-probe rule was applied; other array processing followed the study-specific scripts in the Zenodo archive. In this disease-comparison analysis, PRJNA1184491 used the six-case stromal subset, with central and peripheral counts added within each donor before the count-matrix transformation. The resulting KC comparison comprised 97 KC and 89 control arm observations across nine arms from eight source studies.

For each disease-versus-control comparison, Hedges' *g* was calculated[64] as *g* = J(μcase − μcontrol)/S, where S is the pooled within-group standard deviation and J = 1 − 3/[4(ncase + ncontrol) − 9]. The 95% confidence intervals used a first-order delta approximation that retained the separate case and control variances. Within-study *P* values were obtained from two-sided Welch tests of the sample-level scores. KC effects were synthesized by restricted maximum likelihood with study arms nested within source studies (metafor::rma.mv, random = ~1 | source_study/study)[60]. Bacterial/fungal keratitis and FECD were synthesized separately using restricted maximum-likelihood random-effects models. These disease-comparison estimates retained their own preprocessing, sample subsets, and model definitions throughout (Supplementary Table S10).

Cross-disease differences were defined as Δg = gKC − gdisease. Shared-control covariance was incorporated in the GSE241715 contrasts. For two contrasts, 1 and 2, sharing n0 controls with variance $s0^2$, the covariance was

$$Cov(g_1, g_2) = \frac{J_1 J_2 s_0^2}{n_0 S_1 S_2} + \frac{g_1 g_2 (n_0 - 1) s_0^4}{2 df_1 df_2 S_1^2 S_2^2},$$

where Si is the pooled case–control standard deviation and dfi is the pooled-variance degrees of freedom for contrast i. The variance of the direct difference was Var(g1) + Var(g2) − 2Cov(g1,g2). When the pooled KC and disease estimates both included GSE241715, its covariance contribution was multiplied by the study's weight in each synthesis:

$$Var(\widehat{\Delta * g *}) = Var(\hat{g}_{KC}) + Var(\hat{g}_D) - 2 w_{KC,241715} w_{D,241715} Cov(g_{KC,241715}, g_{D,241715}).$$

Contrasts without shared controls had a zero covariance term. Pooled estimates and between-disease differences used two-sided Wald tests. Gene-level limma effects and unweighted module means of $\log_2$ fold-changes were reported separately from standardized score effects. The nine-arm KC $\log_2$ fold-change synthesis used unshrunken gene estimates and a multilevel model (Supplementary Table S10).

**Central–Peripheral Expression in PRJNA1184491**

PRJNA1184491 provided 36 regional samples from 10 keratoconus (KC) and 8 donor corneas, with one sequencing run per regional sample.[25] KC specimens were obtained at deep anterior lamellar keratoplasty. Both KC and donor specimens were divided into a central semicircle 2.5 mm in diameter and a peripheral region. Four KC corneas retained the epithelium, Bowman's layer, and stroma; six KC corneas and all eight donor corneas underwent epithelial debridement and contained Bowman's layer and stroma. Central and peripheral samples were retained separately for regional analyses. The primary paired analysis included all 10 KC corneas; a second analysis used the six KC corneas collected after epithelial debridement. The donor comparison included all eight donor corneas in both analyses (Supplementary Table S10).

Salmon quantifications[72] were imported with tximport[74] and the GENCODE v44 transcript-to-gene mapping.[73] Run-level gene counts were combined by BioSample, rounded, and analyzed with DESeq2 default normalization.[75] Each contrast retained genes with at least 10 counts in at least the smaller group's number of samples. The paired KC and donor models were ~ person + region, and separate central and peripheral KC–donor comparisons used ~ status. The difference in regional effects between KC and donors was estimated with ~ status + status:person_n + status:region, with participants nested within diagnosis. Columns containing only zeros in the nested design were removed before fitting the full-rank design matrix. The contrast assigned +1 to the KC regional coefficient and −1 to the donor regional coefficient.

Within each of the eight fixed gene sets, unshrunken gene-level $\log_2$ fold-changes ($\log_2$FCs) and their squared standard errors were synthesized with metafor::rma.mv[60], a gene random intercept, and restricted maximum likelihood (REML). Module membership and the established CCN1/CYR61, CCN2/CTGF, and CXCL8/IL8 aliases were retained. Differences in regional effects were calculated from the gene-level model contrasts before module synthesis. The principal unadjusted analyses retained 15,108 genes for the 10-pair KC comparison, 14,097 for the 8-pair donor comparison, 17,075 for the 10-KC-versus-8-donor regional interaction, 14,697 for the central case–control comparison, and 14,962 for the peripheral comparison. Analyses of the six KC corneas collected after epithelial debridement retained 14,596 genes for the paired comparison and 16,621 for the regional interaction. Genome-wide coefficients that did not converge were identified in the archived gene tables; all coefficients contributing to the eight module syntheses converged.

#### Adjustment for Epithelial-Marker Expression

The epithelial-marker score was the arithmetic mean of $\log_2$(CPM + 1) expression for *KRT12*, *KRT5*, *KRT14*, and *KRT3*. Gene counts mapping to the same symbol were added before calculating counts per million (CPM), whose denominator was the total gene count in each sample. The marker score was centered within diagnosis in each fitted analysis population. The adjusted paired model was ~ person + epi_score + region; the adjusted regional-interaction model was ~ status + status:person_n + epi_score + status:region. Adjusted gene effects were synthesized using the same module definitions and REML gene-random-intercept model. These analyses were performed in all 10 KC pairs and in the six KC pairs collected after epithelial debridement, each compared with the eight donor pairs where applicable. A companion keratocyte-marker score used the same calculation for KERA, ALDH3A1, PTGDS, and CD34. Adjustment retained the gene-filtering thresholds of each corresponding unadjusted model (Supplementary Table S10).

#### Participant-Level Regional Scores

Expression was variance-stabilized jointly across the 36 regional samples, standardized gene by gene, and averaged over the available members of each module. Module-score analyses retained their own score units and inference alongside the gene-effect syntheses. Paired tests enumerated all 1,024 sign flips for 10 KC pairs, 256 for eight donor pairs, and 64 for six KC pairs. Paired confidence intervals used the paired t distribution. Regional differences between KC and donors were tested with lmer(score ~ status * region + (1|person)). For direct before-and-after adjustment comparisons, the paired score models were score ~ region + (1|person) and score ~ epi_score + region + (1|person), and the regional-interaction score models were score ~ status * region + (1|person) and score ~ status * region + epi_score + (1|person). Mixed-model confidence intervals and two-sided tests used Satterthwaite degrees of freedom. The exact paired tests and the mixed-model estimates were reported separately.

Regional AP-1/immediate-early gene (IEG) activity used signed CollecTRI[61] and DoRothEA confidence levels A–C regulons[62], the established target exclusions, $\log_2$(CPM + 1) expression, and decoupleR::run_ulm(minsize = 5)[63]. Activity for each of the six transcription factors was standardized across the 36 regional samples before averaging. Within-person differences were analyzed using the same paired and regional-interaction procedures. Amsler–Krumeich III-versus-IV comparisons involved five KC participants in each group and enumerated all 252 allocations of five participants to each grade. Confidence intervals for these between-grade differences used the Welch t distribution. Central-minus-peripheral epithelial-marker and module-score differences were also compared by Spearman correlation, with the reported two-sided t-approximation *P* values (Supplementary Table S10).

#### Clinical Severity, Same-Eye Tissue Layers, and Atopy

Clinical analyses used GSE204791, GSE151631, and PRJNA799648[10],[11],[24], with eight fixed gene sets and the AP-1/IEG composite. GSE204791 provided epithelial and stromal samples from the same eight KC eyes and 15 control participants, comprising eight epithelial and seven stromal controls. GSE151631 provided 19 KC participants and seven controls. PRJNA799648 provided epithelial samples and clinical measurements from seven KC participants and seven controls. Together, these datasets contained 63 different participants and 71 expression samples. The clinical-association analyses used the KC participants with the relevant measurements; repeated tissue layers and right- and left-eye clinical values retained their participant identifiers (Supplementary Table S10).

#### Expression Processing and Module Scores

GSE204791 expression was processed separately within each tissue layer using normexp background correction with offset 16, quantile normalization, and the mean of duplicate probes. GSE151631 retained genes with at least 10 counts in at least five samples and used DESeq2::vst(blind = TRUE)[75]. PRJNA799648 used the 14 epithelial Salmon quantifications[72] imported with tximport[74] and GENCODE v44 mapping.[73] Genes with at least 10 counts in at least seven samples were retained, and expression was variance-stabilized using the DESeq2 pipeline. VST values sharing a gene symbol were averaged.

For each tissue arm, individual genes were centered and scaled using the KC participants, and module scores were the mean of the available member-gene z scores. The same KC-derived means and standard deviations were applied to controls. Established module membership and alias resolution were retained, with aliases used when the current symbol was unavailable. AP-1/IEG activity used the established signed CollecTRI network[61], target exclusions, and univariate linear model.[63] RNA-sequencing input was $\log_2$(CPM + 1), and array input was log expression. Activities of FOS, JUN, JUNB, FOSL1, ATF3, and EGR1 were standardized within the KC participants before averaging.

**Same-Participant Clinical Associations in PRJNA799648**

The source publication's Supplementary Table S1 linked KC_T_1–KC_T_7 to age, sex, overall disease severity, and 20 bilateral structural measurements. Five participants were classified as mild and two as severe. For each eye, the available measurements were anterior and posterior K1, K2, mean keratometry, and astigmatism, maximum keratometry (Kmax), and thinnest corneal thickness. The clinical-table age of 17 years was used for KC_T_2. Mild and severe were coded 1 and 2. Higher positive curvature and greater absolute posterior K1, K2, or mean curvature were oriented toward greater severity; thickness was oriented toward lower values. Right- and left-eye clinical measurements were analyzed separately as same-participant measurements. The publication did not identify the eye from which each epithelial specimen was collected.

Spearman correlations were the principal clinical-association estimates, with exact = FALSE and Fisher-z confidence intervals. Ordinary linear regressions of score on the clinical variable, age, and sex provided adjusted slopes and t-based confidence intervals. Thickness slopes were expressed per 10 μm lower thinnest thickness, and Kmax slopes per 1 D higher curvature. Overall severity, individual ABCD grades, and continuous clinical measurements were reported separately under their recorded definitions (Supplementary Table S10).

**Same-Eye Epithelial–Stromal Slopes in GSE204791**

The eight paired KC eyes had recorded ABCD grades: A, anterior curvature; B, posterior curvature; C, thinnest thickness; and D, visual function. The observed A grades were 2 and 4, C grades 1–3, and D grades 1, 3, and 4. B equaled 4 in every KC participant and was recorded as nonestimable. Clinical grades were kept as grades. Separate within-layer linear trends included age and sex, while Spearman correlations were calculated within KC without adjustment.

For direct comparisons between tissue layers, each module score was centered and divided by its score standard deviation within the eight KC observations of that layer. The model score_z ~ severity * layer + (1|person_id) estimated the epithelial slope, the stromal slope, and the stromal-minus-epithelial slope difference. Models used REML, a participant random intercept, and Satterthwaite confidence intervals and tests, without additional covariates. If the random-intercept fit was singular, the within-eye stromal-minus-epithelial score difference was regressed on severity, and the separate layer slopes were also reported. All nine C-grade readouts used the REML random-intercept model. Effect units were layer-score standard deviations per clinical grade. Each paired analysis comprised eight participants, eight eyes, and 16 tissue samples (Supplementary Table S10).

**Atopy Comparisons in GSE151631**

Six KC participants had a recorded atopy/allergy diagnosis: eczema, hay fever, or allergies in one; asthma in one; vernal keratoconjunctivitis in three; and seasonal allergic conjunctivitis in one. Thirteen KC participants had Absent recorded. The seven controls formed the control group. Module scores used the center and scale of all 19 KC participants. Atopic and nonatopic KC groups were compared with one another and separately with controls using Welch t tests, mean-score differences, and t-based confidence intervals. Hedges' *g* used the pooled standard deviation[64] and exact gamma small-sample correction, with normal-approximation confidence intervals. *P* values were two-sided. All 19 KC participants were classified as severe, so this dataset contributed atopy comparisons. The PRJNA1184491 Amsler–Krumeich comparisons used participant scores and permutation tests (Supplementary Table S10).

#### Peripheral-Blood RNA in PRJNA799648

The peripheral-blood-derived RNA analysis used three KC and three control libraries from PRJNA799648.[11] Paired-end FASTQ files were quantified with Salmon 1.4.0[72] against the same GENCODE v44[73] gentrome and decoy definitions used for the epithelial analysis. The index used k = 31 and the gencode option; quantification used automatic library-type detection, validated mappings, and sequence- and GC-bias correction (-l A --validateMappings --gcBias --seqBias). Transcript estimates were imported with tximport[74] using the v44 transcript-to-gene mapping. DESeq2 used disease status[75] as the design term and retained genes with at least 10 counts in at least three samples. Gene effects were unshrunken Wald estimates of the KC−control $\log_2$ fold-change.

For each fixed gene set, the gene effects and squared standard errors entered a restricted maximum-likelihood model with a random gene intercept (metafor::rma.mv, yi = $\log_2$FC, V = lfcSE$^2$, random = ~1 | gene)[60]. We report the model mean, 95% confidence interval, two-sided Wald *P* value, and number of estimable member genes. Available gene effects numbered 12 for general inflammation, 12 for corneal-local inflammation, and 16 for stress/immediate-early expression. The unweighted member-gene mean was also reported as the arm-level direction summary (Supplementary Table S10).

For donor-level scores, counts assigned to duplicate symbols were summed, transformed as $\log_2$(CPM + 1), and standardized gene-wise across the six blood samples. Scores were the unweighted means of available module genes. CollecTRI activity[61] used the archived signed network, the ECM/immediate-early target exclusions of the donor analysis, and decoupleR::run_ulm[63] with minsize = 5. The six AP-1/IEG factor activities were standardized across blood samples and averaged. Hedges' *g*, standard errors[64], normal-approximation confidence intervals, and two-sided normal *P* values were calculated using the donor-analysis procedure. Each tissue, including epithelium, was reported as a separate comparison within PRJNA799648 (Supplementary Table S10).

#### Feedback Relative to Pathway Output

Within study, the model was feedback expression = intercept + SMAD3 target score + KC diagnosis + the study's covariates. Feedback genes were removed from the output score. The diagnosis coefficient measured the $\log_2$ expression difference at the same target-output score. GSE77938 retained discovery/replication adjustment; GSE112155 retained sex adjustment. Study-specific estimates and standard errors were pooled using DerSimonian–Laird random-effects weights and normal intervals; study-specific inference used the model's t distribution. For the immediate-early adjustment, the fixed 20-gene stress/immediate-early score (Supplementary Table S2; equal-weight mean of within-study standardized members, with *CCN1/CYR61* and *CCN2/CTGF* aliases) was added to the SMAD7 model, and the seven genes it shares with the SMAD3 target set (*CCN1*, *CCN2*, *EGR2*, *FOS*, *JUN*, *SRF*, *ZFP36*) were removed from the output score; the unadjusted model was refit on that output score for comparison. The six-gene AP-1/IEG expression score was used in the same way. All seven primary studies entered both models with residual degrees of freedom of at least one.

Each independent study contributed one whole-cornea or stromal comparison to the primary synthesis. GSE204791 contributed stroma; the two spatial studies contributed whole-section estimates, with isolated stromal estimates retained separately. Epithelial-only studies were reported by layer. PRJNA1184491 used donor means from ten KC and eight control corneas. Shared donors, layers, and regions were not counted as independent studies. SMAD7 was unavailable in E-MEXP-2777, giving seven studies for SMAD7 and eight for SMAD6.

#### Same-Donor Input, Output, and Matrix Programs

Ligand expression, latent-complex storage/anchoring, THBS1, and integrin-activation programs were modeled separately against SMAD3 target output. Matrix outcomes comprised the ECM, collagen-assembly, and repair scores. The repair score contained CTHRC1, TNC, FAP, THBS2, COMP, SFRP4, and COL11A1. To remove direct membership overlap, TNC, THBS2, COL1A1, COL3A1, COL5A2, COL6A1, COL6A2, and COL6A3 were removed from the SMAD3 target score while keeping the signed weights. Matrix models included the nonoverlapping output score, SMAD7, diagnosis, and the study covariates. Responses and continuous predictors were standardized within each study's complete-case model, so slopes represent outcome SD per predictor SD. Independent-study estimates were combined by DerSimonian–Laird random effects.

Central–peripheral and stromal–epithelial comparisons were formed within donor after shared standardization across the paired locations. They were analyzed separately from the primary whole-cornea/stromal synthesis. The underlying 356 donor-by-arm records include repeated layers, regions, and paired differences and do not represent 356 independent participants.

#### Secondary Cross-Layer Ordering Test

The secondary test of fully reversed module ordering between the layers (Supplementary Table S6) is a max-*P* intersection–union test over prespecified germline and tissue components. E, A, and N denote the ECM, stress/immediate-early, and NRF2 modules. Germline components are the two-source replicated ordering tests E > A and E > N and the ECM joint enrichment; tissue activity components are the pooled one-sided tests for AP-1 and NRF2. For the tissue ordering components A > E and N > E, each gene's $\log_2$ expression was rank-normal transformed within arm [$\Phi^{-1}$((rank − 0.5)/n)], each module score was the negative mean of its available transformed genes, module displacement was Cliff's delta between KC and control samples (positive means lower in KC), and the study contrast was the AP-1 or NRF2 displacement minus the ECM displacement, with GSE204791's two arms averaged within study; the eight study contrasts were tested by an exact one-sided signed-rank test with zero contrasts removed. A > E therefore asks whether the stress/immediate-early module is reduced more than the ECM module. *p*_cross is the maximum over the four reversal components, and *p*_full is the maximum of *p*_cross and the three activity components. The test passes only if every component passes, and the overall *P* is the bottleneck component's *P*; no component *P* values are multiplied or combined.

#### Perturbation Responses

Module behavior under defined state transitions was measured with four operators applied identically to the disease-side data. Two were signed-regulon composites (CollecTRI and DoRothEA

ABC, ULM, six-TF equal-weight composite after within-accession z-standardization, with the same regulon target exclusions as above); two were member-score operators over the member genes of each module (the six AP-1/IEG genes or the ECM module members), a standardized-mean score and a rank score. The rank score averaged each member's within-library rank among all analyzable genes. All scores are oriented toward the KC tissue signature of their module: AP-1/IEG composite and member scores are sign-inverted so that positive values point toward the AP-1/IEG-low KC state, ECM member scores weight each gene by the sign of its KC − control $\log_2$FC pooled by random-effects meta-analysis (restricted maximum likelihood) across the four whole-cornea and stromal arms GSE151631, GSE77938, GSE241715, and GSE204791 stroma, and every Δ reported in Supplementary Table S11 and in the perturbation results is on the oriented scale of its module (negative Δ = away from KC). Contrasts are treated-condition minus serum-free baseline on independent libraries (score ~ condition; ordinary least squares with HC3 heteroskedasticity-consistent standard errors;[66],[67] exact label permutation; Hedges' *g*). Stability was evaluated by member leave-one-out — regulon composites drop one TF, member scores drop one gene, and a sign flip or zero crossing marks instability — and by matched nulls. Per accession × operator × contrast, 50,000 matched sets of the same size were proposed from candidates outside the module and the 10,000 closest retained (base seed 20260823; the accession-, module-, and target-specific offsets are defined in the deposited perturbation analysis script); the regulon operators draw six-factor sets matched on regulon target-count decile, target-expression summaries, and activity correlation and first-principal-component variance fraction, while the member-expression scores draw gene sets matched on expression decile and the corresponding gene-expression summaries, with two-sided add-one empirical *P* and BH adjustment[41] within each accession's testing family: eight tests in human (four operators × two contrasts) and twelve in rabbit (the ECM and AP-1/IEG sets × two contrasts × three matched-null metrics: standardized-mean shift, rank shift, and vector cosine). Rabbit analyses run the two member-score operators only; regulon operators were not run in rabbit.

Exact label-permutation tests enumerated all 70 allocations for four-versus-four human libraries and all 12,870 allocations for eight-versus-eight rabbit libraries. The complete human and rabbit library-level scores, member-omission profiles, and matched-null results are provided in Supplementary Table S11 (blocks A, A_parametric, G, and G_members); the compact serum comparison is shown in Supplementary Fig. S3A,B.

**Methylome and m6A Analyses**

RRBS methylation was analyzed on 1-kb tiles (409,647 tiles tested; Welch tests on logit methylation, BH false discovery rate [FDR]), with module-level assessment against coverage-matched null draws (1,000 per module, matched on tested-tile quartiles, seed 20260804). For m6A, per-gene levels were the within-sample $\log_2$(IP) − $\log_2$(supernatant) contrast; differential m6A used FDR < 0.05, and module-level assessment used running enrichment along the *t*-statistic ranking plus hypergeometric overlap, each evaluated in the all-gene and the protein-coding background, with the m6A dataset's own module definitions (whose stress set includes *EGR3*). RRBS and m6A inference remained at donor or sample level; tiles and probes were not treated as independent observations (Supplementary Table S15).

**Fixed-Case Control Substitution and Candidate Directions**

GSE241715 provided the same four KC corneas for comparison with seven normal, five viral, and five bacterial/fungal corneas. Each sample was transformed as $\log_2$(author-normalized count + 1), and group effects were differences of sample means. The common-gene analysis retained identical measurable genes and inverse-variance weights estimated from the normal comparison. The native-coverage analysis was retained separately. The 42 experimental contrasts were not reselected after the substitution.

The published cargo collection comprised 1,253 candidates. Stable identifiers and the mapping rules defined common coverage; candidates without an estimable expression effect remained unavailable. Only the expression contribution and its associated supplementation/inhibition direction were replaced. Other evidence components, weights, and eligibility remained those of the published pipeline. Candidate scores were normalized and ranked within the same measurable pool for each paired comparison. This replay therefore measures the consequence of replacing the expression input in the pipeline.

The fixed-case resampling used 2,000 paired iterations that retained the common KC group while resampling the relevant donor groups and genes as specified in the source analysis. It is distinct from the source-category reference replay. The exact margin calculation below used the fitted effects and weights without additional resampling.

The GSE77938 replay held all 25 KC corneas fixed and varied only the control class. The six known pathological classes were bullous keratopathy (10 corneas), scar (5), perforation (4), descemetocele (2), keratitis (2), and Fuchs endothelial dystrophy (1); one graft of unclear diagnosis was analyzed separately. Common-gene weights were fixed at the inverse variance of the pooled 25 controls, and the 42 models and 1,224 jointly measurable candidates were those of the main replay. A model or candidate reversed when its classification or supplementation/inhibition direction differed between control classes in at least one of the 15 pairwise substitutions among the six classes. Excluding the Fuchs graft left 10 pairwise substitutions. In the counterfactual using the 20 controls retained by the dataset's authors,[14] the pooled control was compared with the retained 20 and with the five excluded controls on the same fixed KC group, models, and weights. The retained 20 were also recounted by class (bullous keratopathy 9, scar 2, perforation 4, descemetocele 2, keratitis 2, and one graft of unclear diagnosis analyzed separately), with the ten pairwise substitutions among the five labeled classes evaluated on the same models, candidates, and weights. The margin decomposition used the same equations as for GSE241715.

#### Exact Decomposition of the Model Decision

Let v = K − N be the KC–normal expression vector, d = C − N the alternative-control–normal vector, x the experimental-model vector, and W the diagonal matrix of fixed normal-reference weights. The alternative disease vector is v − d. Define

$$a = x^T W v, b = x^T W d, D = \| x \|_W \| v \|_W .$$

The original margin is $m_0$ = a/D, the control projection is q = b/D, and the alternative margin on the original denominator is $m_0$ − q. The actual alternative cosine divides a − b by its own positive norm. Thus a strict sign reversal occurs exactly when

$$a(a-b) < 0.$$

This identity separates the direction and size of the control projection from the magnitude of the original margin. Exact zero was retained as a boundary and nonestimable comparisons retained their reasons. The decomposition introduced no new decision threshold.

For program attribution, the complete gene dictionary assigned each gene to a primary category with its priority order, retaining overlapping labels and an unassigned group. Signed contributions sum to the total projection or cosine change. Absolute contribution shares summarize the distribution of those contributions; they are not cell fractions or mediation proportions. Contributions to the fixed-denominator projection and to the difference between separately normalized cosines were retained as separate quantities.

**Held-Out Normal Comparisons**

All 35 partitions of the seven GSE241715 normal donors into three candidate controls and four held-out controls were evaluated. The same four KC corneas entered each comparison. Within a split, effects and inverse-variance weights were estimated from the candidate controls and then used for the old infectious-control, new normal-control, and held-out-normal comparisons. Evaluation-donor expression did not set weights. Old and new model signs and cargo directions were classified against the held-out-normal comparison on their common measurable set.

Each split retained all 42 models and the complete 1,253-candidate collection, with unavailable candidates reported separately. Gene-direction agreement used genes with a nonzero evaluable direction; predicted zero did not count as agreement. Continuous effect correlation and error were computed separately. Medians and ranges summarize repeated partitions of the same study. The normal choice in this experiment was identical to selection by the normal label (Supplementary Fig. S1C).

**Family-Balanced Reference and Independent Consequences**

The updated direction reference used eight whole-study holdouts, the expert predictions and availability masks, 120 fixed basis functions, study-equal training loss, and shared-KC covariance. Duplicate GSE77938 predictions were combined. Six families received equal prior mass: M1, DerSimonian–Laird pooling, MetaIntegrator, single-source GSE77938, Niu/source-subset methods, and the earlier conditional-reference variants. For the M1 expert, each gene's input came from the fitted observed M1 cell when one existed and otherwise from the exact-layer, parent-layer or all-layer DL prediction, in that order. Candidate weights and the residual term were fitted within training. The balanced-accuracy-selected candidate used ridge 1 with the residual enabled; predictions were refitted and frozen before independent scoring. A separate Spearman-based selection returned the MetaIntegrator anchor.

Balanced accuracy is the mean of upregulated- and downregulated-gene recall. The spatial comparison used donor resampling with fixed predictions, while published bulk differential-expression tables were evaluated by paired gene-row resampling. External tissue effects were excluded from fitting and selection. Du's reported differential-expression entries and Sun's published mRNA comparison supplied independent bulk evaluations; Sun's sex-stratified results share the same study.[30],[31]

The gene-level results released with Niu et al.[82] (supplementary DataSheet 2; limma $\log_2$ fold changes after ComBat across GSE151631, GSE112155, and GSE77938, 20,307 genes) were mapped to stable gene identifiers (20,300 symbols mapped; 20,276 unique keys, keeping the smallest-$P$ row where two symbols shared a key) and scored as a separate published approach beside our implementation of the ComBat–limma approach. They covered 20,276 stable keys, 17,138 of them also covered by our implementation, with the same sign for 15,781 (92.1%; Spearman $\rho = 0.970$), and were scored on the entries every approach shared, without refitting or changing any prediction. Paired differences used the same donor or gene-row resamples (2,000) as the other comparisons.

The downstream replay rescored the original reference, updated reference, and independent tissue effects on identical measured genes before deciding whether an original decision was corrected, preserved, or reversed. This intersection differed from the larger pool available for a reference-only replay. The source-category, fixed-case control-substitution, and independently evaluated method-update analyses are separate experiments.

#### Patient-Derived Culture Transcriptomes

The stimulation dataset contained three KC donors in paired 2D and printed GelMA cultures under vehicle or TGF-β1, TGF-β2, or TGF-β3, yielding 24 libraries. Cultures received a four-hour 10-ng/mL pulse beginning on day 7, and RNA was collected at 72 hours from pulse onset. DESeq2 fitted donor and the eight format–treatment groups; genes with at least ten reads in at least three samples were retained, giving 15,989 genes, with default independent filtering and Benjamini–Hochberg adjustment within contrast. The baseline dataset contained three KC and three normal donors in both formats, with some KC isolates also represented in the stimulation experiment. Disease contrasts were evaluated within format and with format adjustment; disease-by-format interaction used the paired donor design. These baseline contrasts and the KC-only stimulation experiment were analyzed separately. Wald-statistic-ranked enrichment used the gene sets and multiplicity families of the bioprinted-stroma analysis; the interaction contrast was (3D stimulated − 3D vehicle) − (2D stimulated − 2D vehicle). Patient low-expression sets were frozen before the culture comparison from each tissue's own tested background, using the significant downregulated lists and the first-200 rule. The spatial GSE303036 list contained 126 eligible genes; missing cultured-cell measurements remained missing and were not treated as zero effects.

#### Additional Figure Methods and Display Details

#### Fixed-Case Control Substitution (Figure 1)

The 42 model contrasts in Figure 1B are grouped into 17 dataset blocks. Contrasts may share datasets and samples. Figure 1C contains the 1,090 candidates jointly estimable under all three controls from the original 1,253-candidate collection. The other 163 candidates are outside this common evaluable set. Density cells form a 24 × 24 grid with square-root count shading on a common 0–109 count scale.

In Figure 1D, the alternative margin is evaluated on the original denominator, whereas the alternative cosine is separately renormalized. The exact relationship is given under **Exact Decomposition of the Model Decision**. Heatmap cells sum signed gene contributions to the control projection within 80 ordered record bins per model, covering 1,027,578 contribution records across the 84 model–alternative-control comparisons. Records are ordered within row by program group and gene/key; columns therefore do not align the same genes across models. Warm and cool colors denote negative and positive sums with signed-square-root shading.

**Donor and Regional Displays (Figure 2)**

The five readouts in Figure 2A comprise 190 donor–arm observations: 101 KC and 89 control observations from 93 KC and 89 control people. Eight GSE204791 KC donors contribute both layers. The final composite is the mean of gene-standardized expression or factor-standardized activity and is not standardized again. Pale and dark group strips distinguish controls and KC.

The Figure 2B stromal heatmap contains 39 score cells from the same 13 GSE303036 donors. Spots are nested within donors. The whole-section GSE303036 comparison reuses those donors.

Figure 2C uses the same vertical scale for its three paired-score displays. Its gene strips contain 104 finite effects and 16 missing values across 120 positions, representing 20 ordered positions per module and contrast. Gene identities, order, and contrast labels are provided in the Figure 2C source-data table. Difference-in-differences is (KC central−peripheral) − (control central−peripheral); score effects and gene-model $\log_2$FC retain their respective units.

**Genetic Displays (Figure 3)**

The Figure 3A gray density classes contain 1, 2–3, 4–7, 8–15, or at least 16 pathway records. Collagen fibril organization, assembly, and formation have 79, 61, and 90 members. Their MVP European *P* values are 0.175, 0.402, and 0.152; replication of the three terms occurs in FinnGen. They are grouped descriptively, without a combined-family *P* value. Of their 127 union genes, 124, 124, and 126 have tested MAGMA Z statistics in Hardcastle, MVP, and FinnGen. Warm and cool colors denote lower and higher Z statistics.

Figure 3B contains 101, 124, and 120 source–SNP records for Hardcastle, MVP, and FinnGen, respectively. The residual disease component is β_disease − b_CCTβ_CCT, and the fitted lines use the corresponding multivariable b_CRF. SNPs can recur across sources and can be in linkage disequilibrium.

In Figure 3C, original-region stromal context uses the stromal overlap annotation, while added-window context uses credible-variant PIP overlap with stromal-class-enriched accessible peaks. Candidate windows can overlap. Missing comparisons remain distinct from a posterior probability of zero.

Each Figure 3D association track contains 4,134 SNP records. The CCT track uses GCST006366 (17,803 participants). Density cells count records by genomic position and $-\log_{10}(P)$ in six classes: 1, 2–3, 4–7, 8–15, 16–31, and at least 32. Tracks have separate vertical ranges. The credible-set matrix has 32 recorded memberships across eight variants and retains each trait's original PIP. Component membership and the regional accessible-element annotation are separate quantities.

**Feedback Models and Published-Value Extraction (Figure 4)**

GSE77938 feedback and output–matrix models include batch. Figure 4A scatterplots display raw output and SMAD7 values, whereas the forest shows conditional diagnosis coefficients.

For the output–collagen analysis, eight genes shared with downstream endpoint sets were removed from the 183-member SMAD3 output definition, leaving 175 defined members and 138–174 measured members across the seven plotted arms. The original collagen endpoint was retained. The eight-study ligand, storage, THBS1, and integrin input analyses have 121–174 measured output members and pooled slopes of 0.5953, 0.5686, 0.4069, and 0.5438 output SD per input SD, respectively. These are donor-level transcriptional associations.

The 24 published group means in Figure 4C were digitized from Priyadarsini et al.[68] Approximate reading precision is ±5–15 percentage points and is larger for high bars; it is not a confidence interval. The source reports at least three repeats, without a biological-donor count or numerical error magnitudes. Each endpoint is normalized to its own HCF control. The HCF TGF-β3 SMAD7 mRNA mean is approximately 530% of that control.

Figure 4D uses the cell-number-normalized hydroxyproline means reported by Soiberman et al.[70] The source reports five repeats and SEM but does not provide numerical SEM or repeat-level values for this extraction, so no error bars are plotted. Normalization to each cell source's own vehicle permits within-source treatment comparisons.

#### Reference Evaluation Displays (Figure 5)

Each Figure 5A matrix contains 35 splits × 42 contrasts. Split-specific medians are calculated separately and need not sum to 42. The Figure 5B vertical axis spans 50%–85%, and its lines connect paired splits.

Common evaluable pools in Figure 5C contain 8,420 upward and 5,884 downward genes in spatial stroma, 715 upward and 2,712 downward entries in Du, and 565 upward and 2,310 downward entries in Sun. Twenty equal-count bins per direction contain 421/294–295 spatial genes, 35–36/135–136 Du entries, and 28–29/115–116 Sun entries, respectively. M1 includes four control-source classes, with living enucleation separate, and additive layer terms. All three plotted M1 predictions use the fitted postmortem whole-cornea/stroma reference; the Sun measured comparison uses living SMILE stroma. The original frozen reference and the six-family reference remain separate variants.

Paired gains against the best published approach are 1.266 percentage points for spatial M1 versus MetaIntegrator (95% CI, 0.460–1.956), 1.448 for Du's six-family reference versus ComBat–limma (0.438–2.449), and 2.350 for Sun's six-family reference versus ComBat–limma (1.161–3.477). The intervals use 2,000 paired resamples, with donors resampled for spatial data and published entries for bulk tables.

Figure 5D preserves all 1,253 candidate records in 25 bins of 50–51 original rank positions. There are 381 nonestimable candidates in Du and 190 in spatial stroma. Summary bars use estimable denominators. Nonzero bars have a minimum display width equivalent to two pixels at 200 dpi; the single reversed Du candidate has a six-pixel minimum. Printed counts retain their original values. HEPHL1, the reversed Du candidate, has original rank 1193 and appears in the 24th rank bin. These transitions describe RNA-direction decisions.

#### Risk Directions and Functional Outcomes (Figure 6)

Figure 6A has 54 measured gene–arm positions opposite to at least one accepted risk-expression direction. The expanded 791 genetic-evidence-by-arm records reuse signals and tissue arms. GSE241715 and E-MEXP-2777 are living-enucleation arms; repeated GSE204791 layers retain their shared donor identity.

The five measured MMP1 effects in Figure 6B are −5.98, −1.45, −8.41, −6.83, and +0.248 $\log_2$FC. The other four arm positions are not reported. The nomination comparison uses three layer–control class estimates, distinct from the nine arms: two have 95% CIs supporting the nominated expression direction, none has a significant opposite estimate, and one meets neither category. Repeat-level quantitative immunofluorescence values were unavailable for the joint MMP1/MMP12 knockdown comparison.

Figure 6C shows the reported directions of total and phosphorylated c-Jun immunofluorescence; repeat-level intensities were unavailable. Its culture units are distinct from the rabbit units in Figure 6D.

For untreated KC, KC + PNM, and KC + CPNM, Figure 6D displays source Table 1 means ± SD: CCT, 312.40 ± 4.22, 316.80 ± 13.32, and 408.60 ± 7.23 μm; Km, 48.61 ± 1.03, 48.48 ± 1.16, and 42.57 ± 1.93 diopters.[13] That table reports five rabbits per group, the denominator used for the displayed values; the source Methods separately reports ten animals per group. The table and Methods counts are retained under their respective source locations.

### Supplementary Results

#### Genetic and Tissue Estimates

Among 7,132 public pathways, collagen assembly was the only independently replicated family: collagen formation, collagen fibril assembly, and collagen fibril organization. Expanding eligibility to 7,310 pathways retained this family. Its tissue expression decreased against pathological controls (collagen formation $\log_2$FC −1.470; 95% CI −1.679 to −1.260), despite no pooled change across controls (Supplementary Table S3, block G).

Each 1-mm Hg increase in genetically determined CRF reduced disease odds across the three sources (IVW ORs, 0.410–0.485; Hardcastle, 0.452 [95% CI 0.380–0.537]). Protection persisted across MR estimators and after excluding instruments near disease-associated variants; CAUSE favored causal over sharing models in all three sources (Supplementary Table S4, blocks A, C, and D).

CCT was protective in univariable MR (ORs, 0.975–0.978 per μm). When CRF and CCT were fitted together, CRF retained protection (ORs, 0.392–0.527), whereas CCT had no independent effect ($P$ = 0.31–0.63). Reverse MR linked genetic disease risk to lower CRF in all three sources and lower CCT in Hardcastle and MVP. The joint structural-trait projection captured 35.5%–54.3% of disease common-SNP heritability in Hardcastle and 27.4%–38.6% in FinnGen. Resistance to deformation therefore contributes to KC risk beyond thickness.

Within the 17-region set, ten had stromal cell context: three stroma-only and seven shared with epithelium. Their credible variants were enriched in adult corneal open chromatin (1.54-fold; $P$ = 0.012), including keratocyte and stromal stem-cell peaks (1.88- and 1.94-fold; $P$ = 0.015 and 0.013), without enrichment in corneal basal or superficial epithelial peaks. Open elements included *ALDH3A1*, *FOXO1*, *SMAD3*, and *COL5A1*; the *ALDH3A1* element was enriched only in the two stromal populations.

#### Shared COL5A1 Components and Their Regulatory Context

The primary COL5A1–CCT PP4 values were 0.990791 for Hardcastle, 0.984753 for MVP European, and 0.989459 for FinnGen R13 corneal deformation (H7_CORNEALDEFORM). Corresponding disease/CCT credible-set sizes were 3/5, 8/5, and 6/5, with intersections of three, five, and five variants. rs3118516 and rs3118518 occurred on both sides of each primary comparison. A second Hardcastle–CCT component had PP4 0.845986, and an MVP–CRF component had PP4 0.863155. None of the variants in these shared component credible sets overlapped the two adult stromal-class-enriched peak sets. The functionally studied rs3118515 lay outside both credible sets in each of the five shared component pairs. These component-specific annotations are separate from the region-set enrichment in adult corneal chromatin.

The 938-bp amplicon of reporter region 2 contained five shared variants (rs3118518, rs3132306, rs1536482, rs1536483, and rs1536484). The posterior conditional on a shared causal variant, summed over these five, was 0.707 for Hardcastle, 0.632 for FinnGen, and 0.558 for MVP in the primary COL5A1–CCT pairs; the second Hardcastle–CCT component and the MVP–CRF component placed $5.3 \times 10^{-19}$ and $3.3 \times 10^{-4}$ of their posterior there. Reporter regions 1, 3, and 4 held less than $10^{-18}$ of the primary-pair posterior. At the four non-palindromic variants (rs3118518, rs3132306, rs1536482, and rs1536483), the KC risk allele lay on the CCT-lowering haplotype in every source row; the palindromic rs1536484 was left unassigned. In this amplicon, the CCT-lowering haplotype had lower reporter activity in hTK keratocytes (7.9 versus 31.1; $P = 0.004$) and RITVA cells (0.9 versus 2.9; $P = 0.02$), with no allelic difference in RPE1 cells.[54] The assay compares haplotypes of the amplicon, so the activity difference is attributed to the fragment as a whole rather than to one variant; endogenous *COL5A1* expression was not measured for this region.

#### Source, Layer, Spatial, and Regional Detail

The opposing matrix conclusions of Kabza et al.[14] and Shinde et al.[10] corresponded to these control-defined comparisons. Their ECM effects were −2.46 against pathological grafts and +0.82 against postmortem donors. *COL1A1* and *FN1* reversed between these classes; *LOX*, *LOXL1*, and *LOXL2* were lower only against pathological controls. *TGFB1* and *SMAD7* were lower against both, and *MMP9* was lower in every tissue arm measuring it.

Across 182 donors, inflammatory, immediate-early, and TGF-β scores were lower in KC (Hedges' *g*, −0.519 to −0.929), as was AP-1/IEG activity under CollecTRI and DoRothEA ABC (*g*, −0.932 and −0.976). Reductions were strongest in whole cornea/stroma (*g*, −0.759 to −1.379); epithelial arms showed no pooled difference (Supplementary Table S6). SMAD3 target output was lower in the nine-arm pooled analysis (*g*, −0.681; 95% CI, −1.181 to −0.180) and after removing the seven genes shared with the immediate-early score (*g*, −0.620; 95% CI, −1.118 to −0.123); in whole cornea/stroma the corresponding values were −0.777 (95% CI, −1.451 to −0.103) and −0.648 (95% CI, −1.349 to 0.054). The TGF-β signaling pathway genes, which share only CCN2 with the immediate-early score, were also lower after removing it (*g*, −0.485; 95% CI, −0.966 to −0.004; Supplementary Table S6, block K). The four-cohort AP-1 analysis is reported in Supplementary Table S6, blocks F–J.

Both spatial cohorts reproduced lower donor-mean immediate-early scores (Δz, −0.700 and −0.468; exact $P = 7.8 \times 10^{-4}$ and 0.036; Fig. 2B). In the same GSE303036 stromal sections, inflammation and immediate-early scores were lower (−0.129 and −0.580), while DC/immune markers were higher (+0.081; 95% CI 0.021–0.142). Immune-marker abundance and inflammatory-program activity therefore separated within the same tissue (Fig. 2B; Supplementary Table S9). Protein measurements gave the same direction. In unpooled corneas from five KC patients and five postmortem donors, general-inflammatory, local-inflammatory, and ECM protein scores were lower in KC (Hedges' *g*, −1.54, −1.55, and −3.24; exact $P = 0.0079$ each), and immune-effector proteins as a class did not differ ($P = 0.46$).[88] Within that class, the 11 measurable complement-cascade proteins were lower in KC (donor-level difference, −0.845 $\log_2$ units; 95% CI, −1.184 to −0.506; exact $P = 0.0079$). In the cone/non-cone stromal proteome, complement-cascade proteins (43 measured) were lower in the same direction against eye-bank donors (−0.930 in cone and −0.808 in non-cone stroma; four KC and two control specimens).[89] In cone stroma, IL-18 and MMP-9 were among the proteins reported above eye-bank donor levels.[89]

KC opposed the inflammatory and AP-1 activation in bacterial/fungal keratitis (KC−infection Δg, −3.357 to −4.176; all $P < 4.3 \times 10^{-8}$). KC inflammation was also lower than viral keratitis; FECD instead increased ECM and TGF-β (Supplementary Table S10). Immediate-early reduction also occurred in bacterial/fungal keratitis, separating this readout from inflammatory activation.

Local increases remained within the lower overall state. In ten paired KC corneas, central ECM, inflammatory, and TGF-β expression exceeded peripheral expression after epithelial adjustment; differences from donor regional effects were +0.271 to +0.382 $\log_2$FC (all $P \leq 0.023$), and the increases persisted in the six KC corneas lacking epithelium. Peripheral inflammation remained approximately −1.09 $\log_2$FC below donors. Epithelial inflammation increased with severity, and immediate-early expression with steepening and thinning. Atopic and nonatopic KC groups both remained below controls for inflammation (Supplementary Table S10).

#### Conditional Feedback and Same-Donor Transcriptional Links

SMAD7's pooled conditional KC difference was −0.601 $\log_2$ units (95% CI, −0.975 to −0.227; P = 0.0016; seven studies), and SMAD6's was −0.108 (−0.293 to 0.077; P = 0.2518; eight studies). Individual SMAD7 coefficients were negative in all seven primary comparisons. With the stress/immediate-early score added, the pooled KC difference was −0.570 (95% CI, −0.914 to −0.226; P = 0.0012; $I^2 = 42.8\%$), compared with −0.697 (−1.089 to −0.306) for the same output score without the shared genes; the pooled coefficient of the immediate-early score was 0.194 (−0.080 to 0.467). Omitting each study in turn left every pooled interval below zero, and the six-gene AP-1/IEG score gave −0.525 (−0.855 to −0.196). Study-level estimates are in Supplementary Table S8. GSE303036's whole-section coefficient was −0.578 (−0.805 to −0.350), whereas its isolated stromal estimate was −0.358 (−1.291 to 0.575). Substituting that stromal estimate in the independent-study synthesis yielded −0.564 (−1.036 to −0.092; P = 0.0191). Substituting GSE285060's stromal estimate instead yielded −0.698 (−1.076 to −0.320; P = 0.0003).

The same-donor standardized relationships were as follows. These are transcriptional associations with each study's adjustments.

| Relationship | Independent studies | Standardized slope (95% CI) | P |
|---|---|---|---|
| Ligand RNA to SMAD3 output | 8 | 0.595 (0.383–0.807) | $3.79 \times 10^{-8}$ |
| Latent storage/anchoring RNA to output | 8 | 0.569 (0.378–0.759) | $4.96 \times 10^{-9}$ |

| Relationship | Independent studies | Standardized slope (95% CI) | P |
|---|---|---|---|
| THBS1 RNA to output | 8 | 0.407 (0.255–0.559) | $1.56 \times 10^{-7}$ |
| Integrin-activation RNA to output | 8 | 0.544 (0.310–0.777) | $4.91 \times 10^{-6}$ |
| Nonoverlapping output to repair score, adjusted for SMAD7 | 7 | 0.629 (0.360–0.897) | $4.61 \times 10^{-6}$ |
| Nonoverlapping output to ECM, adjusted for SMAD7 | 7 | 0.588 (0.397–0.779) | $1.65 \times 10^{-9}$ |
| Nonoverlapping output to collagen assembly, adjusted for SMAD7 | 7 | 0.570 (0.400–0.741) | $5.61 \times 10^{-11}$ |

Within 18 donors with paired central and peripheral tissue, the ligand–output slope for the regional differences was 0.721 (95% CI, 0.319–1.123), and the output–ECM slope was 0.743 (0.383–1.104). These paired comparisons connect the regional transcriptional changes within individuals.

#### Culture Responses Retain Their Baseline, Induction, and Endpoint Meanings

TGF-β1 induced SMAD7 in 2D by 2.4421 $\log_2$ units (adjusted P = $1.6139 \times 10^{-20}$) and in 3D by 0.9255 (adjusted P = 0.0010775). The separate baseline disease comparison gave a format-adjusted SMAD7 difference of −0.024818 (adjusted P = 0.98958). The collagen interaction enrichment was positive (NES 1.92, adjusted P = 0.013), while the myogenesis interaction was negative (NES −1.37, adjusted P = 0.037). The frozen contractile module's 3D baseline was lower than its 2D baseline by 1.2479 $\log_2$ units; induction was 2.6590 $\log_2$ units in 3D and 2.4201 in 2D, leaving the stimulated 3D endpoint 1.0090 $\log_2$ units lower. Thus the lower endpoint in this module arose from the baseline difference despite stronger induction in 3D. In the 3D TGF-β1 comparison, the frozen low-expression sets from GSE204791 and PRJNA1184491 had NES −2.1116 and −1.4095 (adjusted P = $5.9589 \times 10^{-7}$ and 0.031538), while the GSE77938 set had NES +2.1952 (adjusted P = $3.9634 \times 10^{-8}$). The spatial GSE303036 set had NES +1.1131 (adjusted P = 0.30668). These results separate matrix-program recruitment from uniform reversal of tissue-low expression.

#### What the Normal Anchor Changes

With KC samples fixed, POSTN's mean $\log_2$ expression difference was +2.1366 against normal cornea, −4.6291 against viral cornea, and −2.9945 against bacterial/fungal cornea. The infectious controls themselves were +6.7657 and +5.1311 above normal. COL1A1 showed the same distinction: KC−normal was +0.4385, while viral−normal and bacterial/fungal−normal were +3.4455 and +3.6775. The seven-gene repair score was approximately unchanged between KC and normal (+0.0013 standardized units), while the two infectious groups were +1.1259 and +1.2038 above normal. This decomposition places each side of the comparison relative to the same normal group.

The common-gene, fixed-weight analysis reversed 27/42 viral-control and 21/42 bacterial/fungal-control model signs. In the native-coverage analysis the corresponding counts were 24/42 and 18/42. Candidate directions changed for 361/1,090 and 385/1,090 in the common estimable pool. These counts are distinct from the fitted source-category replay, which changed 17/42 model signs and 286/1,210 candidate operations.

For every viral-control flip, the control projection had the same sign as the original margin and exceeded it in magnitude; projection-to-margin ratios ranged from 1.084 to 34.890. Of the 15 unchanged models, 14 had oppositely signed projections and one had a same-sign projection below the margin. All 21 bacterial/fungal flips likewise crossed the original margin. As a named example, the corneal-curvature high-versus-flat contrast changed from cosine +0.149496 to −0.030193 after the viral-control substitution; its fixed-denominator control projection was +0.181361.

Among flipped models, epithelial-identity programs accounted for 26.9127% and 29.2383% of the absolute cosine-change contributions in the viral and bacterial/fungal comparisons; ECM/stromal programs accounted for 13.0373% and 12.7956%. Unassigned contributions remained 40.8614% and 40.0727%. These distributed contributions explain why a single repair transcript does not account for every genome-wide decision change.

**Held-Out-Normal Recovery Includes Candidate Operations**

Across the 35 normal-donor partitions, the median numbers of model decisions corrected relative to the held-out-normal comparison were 20 for viral controls and 17 for bacterial/fungal controls, with 4 and 5 formerly concordant decisions reversed. Among all 1,253 candidates, median corrected counts were 238 and 269; corresponding formerly concordant directions reversed were 138 and 133. Median unavailable counts were 199 and 197. Each category median was computed separately.

Median gene-direction agreement increased from 58.81% to 77.38% in the viral comparison and from 59.59% to 77.37% in the bacterial/fungal comparison. Median model-score Spearman correlations with the held-out-normal comparison rose from −0.033 to 0.824 and from 0.047 to 0.826. Candidate-score correlations rose from 0.771 to 0.840 and from 0.788 to 0.839. These repeated partitions measure recovery of the same study's normal comparison.

**Reference Benchmark and Model Rankings**

We selected the whole-cornea/stromal, postmortem-donor reference before scoring the independent GSE303036 cohort of eight KC and five control donors. On the 14,304 stromal genes scorable by every method, direction balanced accuracy was 60.60%, exceeding GSE77938 alone (55.56%), ComBat–limma (58.37%), MetaIntegrator (59.33%), and DerSimonian–Laird pooling (59.31%); every paired gain excluded zero, including 1.27 percentage points (95% CI 0.46–1.96) over the highest-scoring published approach. The reference also ranked first in endothelium (57.46%; gain over MetaIntegrator, 0.93 points, 95% CI 0.14–1.74) and exceeded all four approaches for effect ranking in the stromal, endothelial, immune, and whole-section views. Among 175 jointly measurable genes in the 200 strongest stromal effects, direction balanced accuracy was 84.5%, versus 78.3% for the best published approach. Adding the gene-level results released with Niu et al.[82] restricted scoring to the 13,365 stromal genes they cover: they reached 57.63% there, below the reference (60.51%) and MetaIntegrator (59.22%). The reference ranked stromal, endothelial, and immune effects better than all five published results; in whole-section views the released results ranked effects slightly higher (Spearman ρ 0.583 versus 0.565 for the reference; difference −0.018, 95% CI −0.071 to 0.040). In the 200 strongest stromal effects, the released results were the strongest published approach (80.1% on 172 jointly measurable genes, versus 84.5% for the reference; difference, 4.4 points; 95% CI, 3.1–4.5), and for the strongest 500 and 1,000 effects the reference scored 82.9% and 80.2% versus 81.6% and 78.2%.

The independent Du bulk cohort (31 DALK KC stroma, four eye-bank donors) reproduced lower *FOS*, *EGR1*, *JUN*, and *SMAD7* ($\log_2$FC, −3.88, −3.65, −3.00, and −3.01).[30] The selected reference correctly predicted 50 of 54 measured entries for the reduced tissue programs. Matrix genes also decreased in this cohort. Across 3,427 entries in Du's reported differential-expression table scorable by every method, directional balanced accuracy favored ComBat–limma (91.6% versus 79.1%). The low inflammatory and immediate-early state thus reproduced with different matrix directions, including across cohorts using eye-bank controls. Reference performance also differed between these tissue comparisons.

The closest of 42 experimental contrasts to pooled KC tissue was an irradiated 3T3-J2 feeder layer (cosine +0.113; 95% CI +0.091 to +0.134), which was stiffer, expressed more matrix, and supported less epidermal stem-cell self-renewal.[77] The five layer-by-control references had pairwise cosines of −0.15 to +0.36; each changed the top-ranked model, and the feeder fell to ranks 4–41.

Changing only the whole-cornea/stromal control from postmortem donors to pathological tissue reversed 17 of 42 similarities, eight with both intervals excluding zero. Corneal stretch at 6 h versus 0 h changed from +0.083 to −0.034. Genes outside ECM and the reduced core programs accounted for 92%–101% of these changes. On common genes, exchanging reference effects alone reproduced 14 of 17 reversals. Reference choice therefore changed the model selected to represent KC.

**Independent Bulk and Spatial Reference Comparisons**

The family-balanced direction reference achieved 93.085% balanced accuracy in Du, versus 91.637% for the best published approach (difference, 1.448 percentage points; 95% CI, 0.438–2.449). In Sun's independent KC–SMILE mRNA comparison, the corresponding values were 92.273% and 89.923% across 2,875 commonly evaluated entries (difference, 2.350 points; 95% CI, 1.161–3.477).[31] Spatial stromal accuracy was 59.325%, compared with 59.330% for MetaIntegrator and 60.596% for the original M1 reference. The family-balanced reference's stromal effect-ranking correlation (14,304 entries) was 0.2666, compared with 0.3245 for M1. Against the gene-level results released with Niu et al.,[82] scored on the entries every approach shared (3,232 in Du; 2,738 in Sun), the family-balanced reference reached 93.21% versus 91.16% in Du (difference, 2.06 points; 95% CI, 1.04–3.11) and 92.36% versus 90.05% in Sun (difference, 2.31 points; 95% CI, 1.22–3.39). In spatial stroma on 13,365 genes it scored 59.27% versus 57.63% (difference, 1.65 points; 95% CI, 0.11–2.42).

On the identical measured set used for each downstream comparison, the Du update corrected nine model classifications and reversed one previously correct classification; 29 remained correct and three remained incorrect. Among 872 measurable candidates, 171 directions were corrected and one previously correct direction reversed; 698 remained correct and two remained incorrect. The spatial update corrected eight model classifications and reversed eight; among 1,063 measurable candidates, it corrected 76 and reversed 150, with 436 remaining correct and 401 remaining incorrect. COL1A1 illustrates the context difference: the update corrected its direction against Du but moved away from its measured direction in spatial stroma.

**Culture Handling and Loading History**

Culture studies separated the low tissue state from responses to stimulation. Serum raised immediate-early readouts in normal-source human and rabbit stromal cells (Supplementary Fig.

S3A,B). In Akoto et al.'s KC-derived cells,[71] general and local-inflammatory expression remained lower after adjustment for TGF-β1 and stretch (mean $\log_2$FC, −0.446 and −0.480). All four measured AP-1 members were negative, while inferred factor activities had mixed directions. These were diagnosis effects across culture conditions. Brief tissue digestion independently raised immediate-early expression (+3.04 $\log_2$ units) without raising inflammation or inferred AP-1 activity.[81]

Acute cyclic stretch of human corneal fibroblasts opposed pooled KC tissue (cosine −0.163; 95% CI −0.187 to −0.139), raised immediate-early, AP-1, and *PTGS2* expression, and opposed all five references.[12] *PTGS2* was lower against all three stromal control classes and in GSE303036 sections ($g = -1.97$); neither *PTGS2* nor *PIEZO1* was significantly increased in the study's source cohort, GSE285060.[29]

Longer loading shifted human fibroblast and mouse skin responses toward pooled KC (interaction cosines +0.055 and +0.047), although both endpoints remained negative. The 1,566 genes also lower in KC contributed 130% and 123% of the net shift, offset by other genes; only 13 belonged to the reduced core programs. Week-scale skin expansion became concordant with KC while immediate-early and AP-1 expression increased. Release retained mechanical history: mouse lung fibroblasts moved from a negative pooled-reference projection on rigid culture to a positive projection after transfer to soft 3D collagen, with low AP-1 activity persisting at the loaded level. Rat stromal cells showed a positive pooled-reference projection after release. Global resemblance, the reduced core programs, and recovery after unloading were distinct outcomes.

**Resource Provenance and Operation-Specific Detail**

Comparator definitions propagated into downstream resources and interventions. CorneaMD labeled the GSE77938 non-KC group as healthy (corneamd.cn/transcriptomics/4/), although its controls were pathological grafts. Chao's exosome-cargo pipeline[15] used this dataset to rank *COL1A1* first. Replacing only its differential-expression component with the postmortem-donor stromal reference reversed supplementation versus inhibition for 286 of 1,210 estimable candidates; *COL1A1* moved from first to sixth within that estimable set. The same replay against pathological controls reversed none of 1,144 estimable candidates.

In GSE77938 with all 25 KC corneas fixed, substituting one of six pathological control classes for another reversed 14 of 42 model classifications at least once (11 of 42 across the ten substitutions that exclude the single Fuchs graft) and the supplementation/inhibition direction of 773 of 1,224 jointly measurable candidates (47 without the Fuchs graft). The Fuchs graft alone reversed eight models and 749 of the 773 candidates. Using the 20 controls retained by the dataset's authors[14] instead of the pooled 25 changed 2 of 42 model classifications and none of the 1,224 candidate directions; using only the five excluded controls changed 2 of 42 models and 256 candidate directions, all from supplementation to inhibition. Within the retained 20, substitutions among the five labeled classes (ten pairwise substitutions) reversed 11 of 42 model classifications and the supplementation/inhibition direction of 63 of 1,224 candidates (48 of them among the 773 candidates reversed across the six classes, and 15 new); including the graft of unclear diagnosis as a sixth group (15 substitutions) gave 13 models and 96 candidates.

Of 13 genes with genetic-risk and measured tissue-expression directions, 12 were discordant in at least one KC arm. The *SGCA* risk allele increased fibroblast and skin expression, whereas expression was lower in all six measured KC arms. *LOX* transcript direction varied by layer and control, while published measurements placed its KC defect in enzyme activity and protein distribution [S5]. At *FNDC3B*, the risk allele and seven of nine KC tissue arms showed lower expression, but lowering expression reduced stress fibers in embryonic fibroblasts[101] and increased TGF-β1-induced stress fibers in HeLa cells.[102]

The cargo analysis yielded a direct example. *MMP1* was lower against both postmortem and pathological controls and was nominated as replacement cargo.[15] Yet combined *MMP1/MMP12* knockdown partly restored collagen I after corneal-cell stretch.[12] The disease transcript direction and the matrix-preserving operation pointed in opposite directions. Likewise, *DANCR* had been nominated because its expression was reduced in KC, yet further knockdown reduced oxidative DNA damage and improved survival in non-KC corneal epithelial cells without lowering reactive oxygen species.[104]

S3. Phipson B, Smyth GK. Permutation P-values should never be zero: calculating exact P-values when permutations are randomly drawn. Stat Appl Genet Mol Biol. 2010;9:Article39.

S4. Han H, Cho JW, Lee S, et al. TRRUST v2: an expanded reference database of human and mouse transcriptional regulatory interactions. Nucleic Acids Res. 2018;46(D1):D380-D386.

S5. Dudakova L, Liskova P, Trojek T, et al. Changes in lysyl oxidase (LOX) distribution and its decreased activity in keratoconus corneas. Exp Eye Res. 2012;104:74-81.
