## Supplementary Figures S1-S6 for "Reinterpreting Keratoconus Gene Expression in Light of Inherited Stromal Weakness"

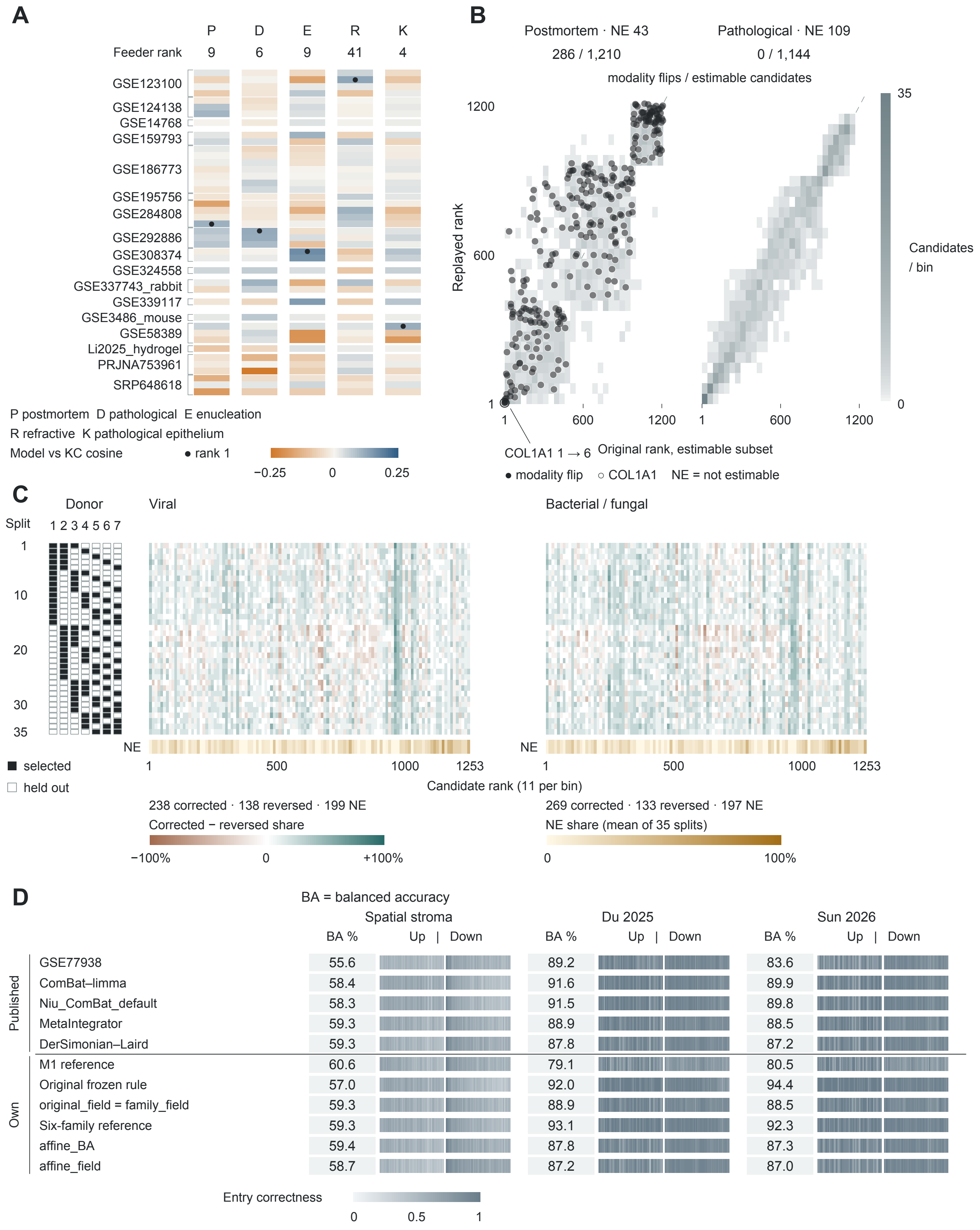
A
P D E R K
Feeder rank 9 6 9 41 4
GSE123100
GSE124138
GSE14768
GSE159793
GSE186773
GSE195756
GSE284808
GSE292886
GSE308374
GSE324558
GSE337743_rabbit
GSE339117
GSE3486_mouse
GSE58389
Li2025_hydrogel
PRJNA753961
SRP648618
P postmortem D pathological E enucleation
R refractive K pathological epithelium
Model vs KC cosine
● rank 1
−0.25 0 0.25
B
Postmortem · NE 43
Pathological · NE 109
286 / 1,210
0 / 1,144
modality flips / estimable candidates
Replayed rank
1200
600
1
35
0
Candidates / bin
1 600 1200 1 600 1200
COL1A1 1 → 6
Original rank, estimable subset
● modality flip ○ COL1A1 NE = not estimable
C
Donor
Split
1 2 3 4 5 6 7
1
10
20
30
35
Viral
Bacterial / fungal
NE
NE
1 500 1000 1253
1 500 1000 1253
Candidate rank (11 per bin)
■ selected
□ held out
238 corrected · 138 reversed · 199 NE
Corrected − reversed share
−100% 0 +100%
269 corrected · 133 reversed · 197 NE
NE share (mean of 35 splits)
0 100%
D
BA = balanced accuracy
Spatial stroma
Du 2025
Sun 2026
BA % Up | Down
Published
GSE77938 55.6 89.2 83.6
ComBat–limma 58.4 91.6 89.9
Niu_ComBat_default 58.3 91.5 89.8
MetaIntegrator 59.3 88.9 88.5
DerSimonian–Laird 59.3 87.8 87.2
Own
M1 reference 60.6 79.1 80.5
Original frozen rule 57.0 92.0 94.4
original_field = family_field 59.3 88.9 88.5
Six-family reference 59.3 93.1 92.3
affine_BA 59.4 87.8 87.3
affine_field 58.7 87.2 87.0
Entry correctness
0 0.5 1

**Supplementary Figure S1. Complete model projections, candidate replays, and independent reference comparisons retain their object-specific outcomes.**

**(A)** Signed weighted cosine similarity of 42 experimental contrasts to five KC−control reference vectors, grouped into 17 datasets. The reference codes identify the control comparison: P, postmortem whole cornea/stroma; D, pathological whole cornea/stroma; E, living-enucleation whole cornea/stroma; R, healthy/refractive epithelium; K, pathological epithelium. Filled dots identify rank 1 within each reference; intervals are not shown. The irradiated 3T3-J2 feeder contrast GSE324558 LQ−SQ ranks 9, 6, 9, 41, and 4, respectively, among 42 contrasts. References and contrasts reuse inputs. Supplementary Table S12 reports projections.

**(B)** Original versus replayed candidate ranks after replacing only the published cargo pipeline's differential-expression component. All other evidence and weights are retained. Against the postmortem reference, 286 of 1,210 estimable candidates change supplementation/inhibition direction, with 43 NE; against the pathological reference, 0 of 1,144 change, with 109 NE. Each begins from 1,253 candidate records. Ranks are within each estimable subset. Density cells count candidates in bins of at most 42 rank positions, with common square-root shading on a 0–35 count scale. Individual flip points are overlaid. The outlined COL1A1 point moves from rank 1 to rank 6 in the postmortem subset; the diagonal marks equal ranks. Numerical summaries are in Supplementary Table S12.

**(C)** Candidate decisions across the 35 overlapping partitions of seven normal donors (three selected, four held out), with four KC cases fixed. Filled and open donor-strip cells identify selected and held-out donors. The two matrices retain 87,710 records (35 splits × 2 old-control groups × 1,253 candidates), ordered by the original normal-comparison rank. Each of 114 bins contains 11 original rank positions, except the last with 10. Color represents (corrected − reversed)/all candidate records in the bin; both-correct, both-incorrect, and NE records remain in that denominator. Teal indicates a positive signed proportion and brown a negative proportion. Separate NE strips show the mean of the 35 split-specific NE proportions per bin. Median corrected/reversed/NE counts are 238/138/199 for viral and 269/133/197 for bacterial/fungal controls; separately computed medians need not sum to 1,253. Supplementary Table S13 contains split summaries.

**(D)** Independent directional BA for five published methods and eight reference entries on GSE303036 stroma, Du bulk, and Sun bulk truth. Two pairs with identical predictions share rows, giving 11 displayed rows for 13 method entries. The merged rows represent Original frozen rule and the field-selected original/six-family references. GSE77938, ComBat–limma, MetaIntegrator, and DerSimonian–Laird identify their corresponding single-arm, batch-adjusted, and meta-analysis predictions; default denotes the default ComBat implementation, field denotes field-selected variants, and affine denotes affine variants. M1 is the fitted meta-regression reference with four control-source classes, living enucleation separate, and additive tissue-layer terms. The spatial, Du, and Sun columns all use its fitted postmortem whole-cornea/stroma route; Sun truth is living SMILE-derived stroma and retains that historical prediction route. All displayed direct M1 rows are fitted predictions. Each truth-direction strip retains 60 ordered equal-count rank bins, ordered by decreasing absolute truth $\log_2$FC; color is correct/n. The common pools contain 14,304 spatial genes (8,420 up/5,884 down), 3,427 Du entries (715/2,712), and 2,875 Sun entries (565/2,310). BA averages the two direction-specific recalls equally; equal strip widths do not imply equal entry counts. Bold rows identify reference anchors. This panel prints point estimates; spatial CIs elsewhere resample donors and bulk CIs resample published entries. Complete scores and paired comparisons are in Supplementary Table S13.

KC, keratoconus; BA, balanced accuracy; M1, model 1; NE, not estimable; CI, confidence interval; SMILE, small-incision lenticule extraction; $\log_2$FC, $\log_2$ fold change.

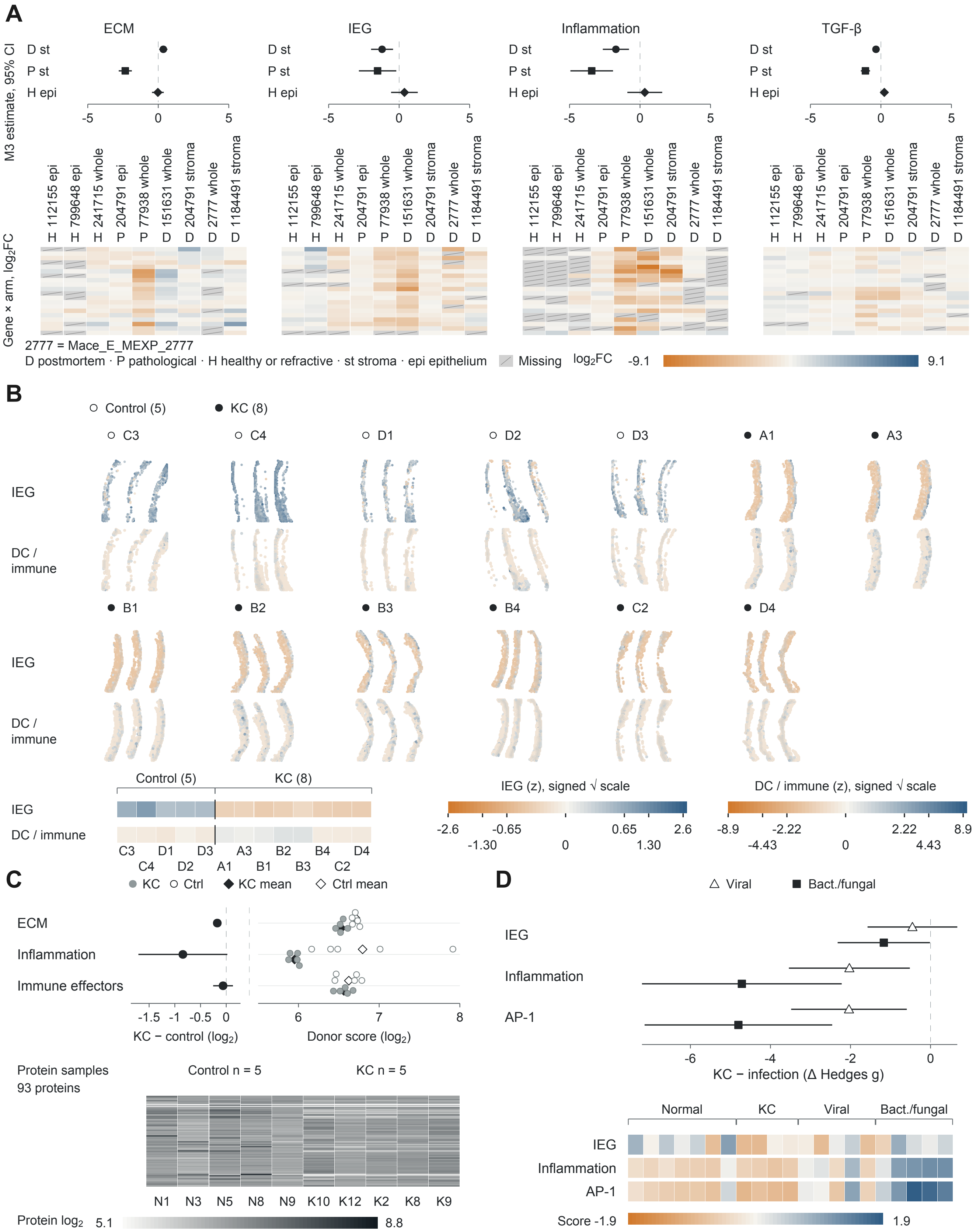

A
ECM
IEG
Inflammation
TGF-β
D st
P st
H epi
M3 estimate, 95% CI
-5
0
5
112155 epi
799648 epi
241715 whole
204791 epi
77938 whole
151631 whole
204791 stroma
2777 whole
1184491 stroma
H H H P P D D D D
Gene × arm, log2FC
2777 = Mace_E_MEXP_2777
D postmortem · P pathological · H healthy or refractive · st stroma · epi epithelium
Missing
log2FC -9.1 9.1
B
Control (5)
KC (8)
C3 C4 D1 D2 D3 A1 A3
IEG
DC / immune
B1 B2 B3 B4 C2 D4
IEG
DC / immune
Control (5) KC (8)
IEG
DC / immune
C3 D1 D3 A3 B2 B4 D4
C4 D2 A1 B1 B3 C2
IEG (z), signed √ scale
-2.6 -0.65 0.65 2.6
-1.30 0 1.30
DC / immune (z), signed √ scale
-8.9 -2.22 2.22 8.9
-4.43 0 4.43
C
KC Ctrl KC mean Ctrl mean
ECM
Inflammation
Immune effectors
-1.5 -1 -0.5 0
KC − control (log2)
6 7 8
Donor score (log2)
Protein samples
93 proteins
Control n = 5
KC n = 5
N1 N3 N5 N8 N9 K10 K12 K2 K8 K9
Protein log2 5.1 8.8
D
Viral
Bact./fungal
IEG
Inflammation
AP-1
-6 -4 -2 0
KC − infection (Δ Hedges g)
Normal KC Viral Bact./fungal
IEG
Inflammation
AP-1
Score -1.9 1.9

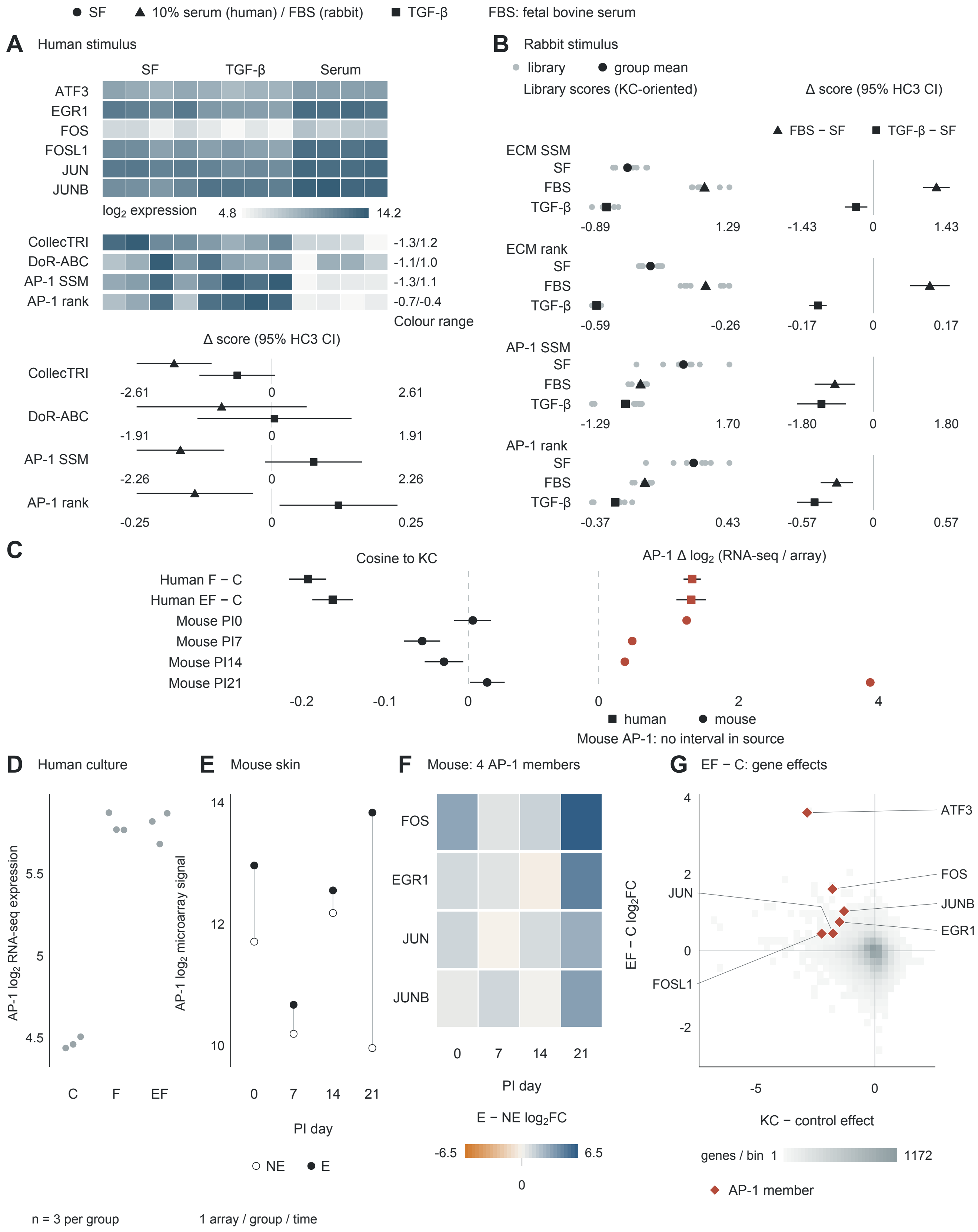

SF
10% serum (human) / FBS (rabbit)
TGF-β
FBS: fetal bovine serum
A Human stimulus
SF
TGF-β
Serum
ATF3
EGR1
FOS
FOSL1
JUN
JUNB
log2 expression
4.8
14.2
CollecTRI
DoR-ABC
AP-1 SSM
AP-1 rank
-1.3/1.2
-1.1/1.0
-1.3/1.1
-0.7/-0.4
Colour range
Δ score (95% HC3 CI)
-2.61
0
2.61
-1.91
1.91
-2.26
2.26
-0.25
0.25
B Rabbit stimulus
library
group mean
Library scores (KC-oriented)
Δ score (95% HC3 CI)
FBS − SF
TGF-β − SF
ECM SSM
ECM rank
FBS
-0.89
1.29
-1.43
1.43
-0.59
-0.26
-0.17
0.17
-1.29
1.70
-1.80
1.80
-0.37
0.43
-0.57
0.57
C
Cosine to KC
AP-1 Δ log2 (RNA-seq / array)
Human F − C
Human EF − C
Mouse PI0
Mouse PI7
Mouse PI14
Mouse PI21
-0.2
-0.1
2
4
human
mouse
Mouse AP-1: no interval in source
D Human culture
AP-1 log2 RNA-seq expression
5.5
5
4.5
C
F
EF
n = 3 per group
E Mouse skin
AP-1 log2 microarray signal
14
12
10
7
14
21
PI day
NE
E
1 array / group / time
F Mouse: 4 AP-1 members
PI day
E − NE log2FC
-6.5
6.5
G EF − C: gene effects
EF − C log2FC
KC − control effect
-5
-2
genes / bin
1
1172
AP-1 member

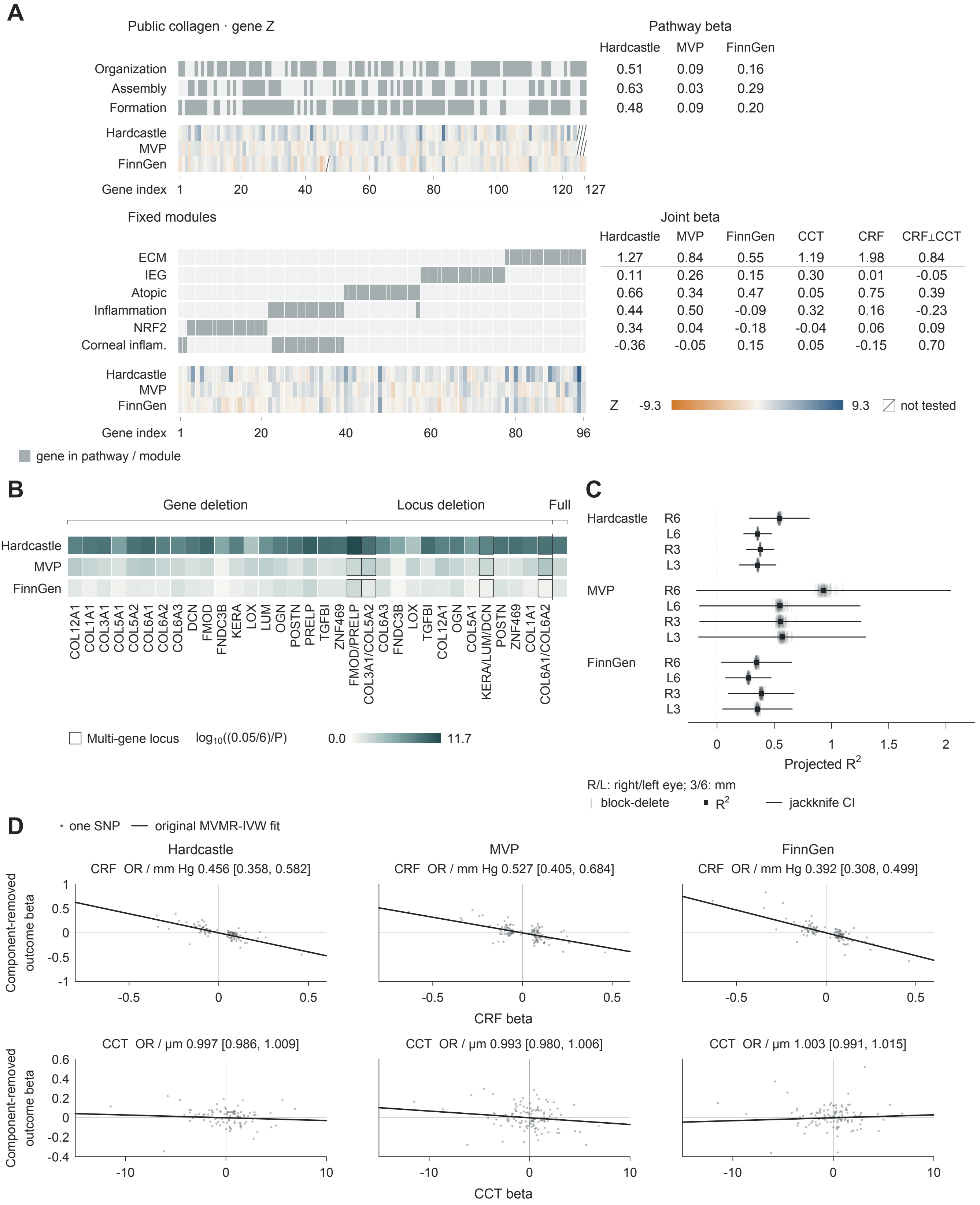
A
Public collagen · gene Z
Pathway beta
Hardcastle MVP FinnGen
Organization 0.51 0.09 0.16
Assembly 0.63 0.03 0.29
Formation 0.48 0.09 0.20
Hardcastle
MVP
FinnGen
Gene index 1 20 40 60 80 100 120 127
Fixed modules
Joint beta
Hardcastle MVP FinnGen CCT CRF CRF⊥CCT
ECM 1.27 0.84 0.55 1.19 1.98 0.84
IEG 0.11 0.26 0.15 0.30 0.01 -0.05
Atopic 0.66 0.34 0.47 0.05 0.75 0.39
Inflammation 0.44 0.50 -0.09 0.32 0.16 -0.23
NRF2 0.34 0.04 -0.18 -0.04 0.06 0.09
Corneal inflam. -0.36 -0.05 0.15 0.05 -0.15 0.70
Hardcastle
MVP
FinnGen
Z -9.3 9.3 not tested
Gene index 1 20 40 60 80 96
gene in pathway / module
B
Gene deletion
Locus deletion
Full
Hardcastle
MVP
FinnGen
COL12A1 COL1A1 COL3A1 COL5A1 COL5A2 COL6A1 COL6A2 COL6A3 DCN FMOD FNDC3B KERA LOX LUM OGN POSTN PRELP TGFBI ZNF469
FMOD/PRELP COL3A1/COL5A2 COL6A3 FNDC3B LOX TGFBI COL12A1 OGN COL5A1 KERA/LUM/DCN POSTN ZNF469 COL1A1 COL6A1/COL6A2
Multi-gene locus
log10((0.05/6)/P)
0.0 11.7
C
Hardcastle R6 L6 R3 L3
MVP R6 L6 R3 L3
FinnGen R6 L6 R3 L3
0 0.5 1 1.5 2
Projected R2
R/L: right/left eye; 3/6: mm
block-delete
R2
jackknife CI
D
one SNP
original MVMR-IVW fit
Hardcastle
CRF OR / mm Hg 0.456 [0.358, 0.582]
MVP
CRF OR / mm Hg 0.527 [0.405, 0.684]
FinnGen
CRF OR / mm Hg 0.392 [0.308, 0.499]
Component-removed outcome beta
1 0.5 0 -0.5 -1
-0.5 0 0.5
CRF beta
CCT OR / µm 0.997 [0.986, 1.009]
CCT OR / µm 0.993 [0.980, 1.006]
CCT OR / µm 1.003 [0.991, 1.015]
Component-removed outcome beta
0.6 0.4 0.2 0 -0.2 -0.4
-10 0 10
CCT beta

**Supplementary Figure S4. Gene membership, deletion analyses, structural-trait projection, and MR instruments resolve the genetic signal.**

**(A)** Membership of collagen fibril organization, assembly, and formation (79, 61, and 90 genes; union, 127; three-way intersection, 28), with source-specific competitive MAGMA coefficients and gene Z statistics. The separate six-module display contains 96 distinct mapped genes and 114 memberships, with joint coefficients for Hardcastle, MVP European, FinnGen R13, CCT, CRF, and CRF⊥CCT. Each narrow column is an Entrez gene; the two gene universes have separate ordering and index scales. Gene order is provided in the Source Data. Gray membership cells identify included genes, and hatching identifies genes untested in a source. Public-pathway Z statistics are available for 124, 124, and 126 genes by disease source; all 96 mapped fixed-module genes have Z statistics in each source. Coefficients and gene Z statistics retain separate scales. Supplementary Tables S2 and S3 provide definitions and enrichment results.

**(B)** Marginal ECM enrichment after each of 19 single-gene deletions and 14 physical-locus deletions in each of three disease sources (57 and 42 deletion analyses). Locus labels identify all genes removed together; outlined cells mark multigene loci. The separate Full cells show each source's undeleted marginal ECM model. Color encodes $\log_{10}[(0.05/6)/P]$; zero marks the six-module Bonferroni threshold, and every deletion remains below that *P*-value threshold. Genes share loci and analyses share input data. Supplementary Table S3 contains the deletion results.

**(C)** Joint structural-trait projections onto disease common-SNP heritability for right- and left-eye 3-mm and 6-mm measurement families. Squares and lines show the original $R^2$ projections and 200-block jackknife 95% CIs; pale ticks show the 200 block-deletion estimates for each source–family combination (2,400 values). The final combinations include CCT, CRF, and five corneal measurement families. Eye, measurement-family, and disease-source estimates share inputs. Intervals retain their original endpoints, including values outside 0–1. Results are in Supplementary Table S5.

**(D)** Multivariable CRF and CCT components for Hardcastle, MVP, and FinnGen. Each point is a harmonized source–SNP record (101, 124, and 120 per exposure), plotting the selected exposure's effect against β_KC minus the fitted contribution of the other exposure. Lines through the origin use the corresponding no-intercept multiplicative random-effects IVW coefficient. The two exposure rows reuse each source's instruments and have separate coordinate scales; within each exposure row, the y-axis scale is shared across disease sources. CRF ORs per mm Hg are 0.456 (95% CI, 0.358–0.582), 0.527 (0.405–0.684), and 0.392 (0.308–0.499); CCT ORs per µm are 0.997 (0.986–1.009), 0.993 (0.980–1.006), and 1.003 (0.991–1.015). Complete estimates are in Supplementary Table S4.

KC, keratoconus; ECM, extracellular matrix; IEG, immediate-early genes; NRF2, nuclear factor erythroid 2-related factor 2; Corneal inflam., corneal inflammation; MAGMA, Multi-marker Analysis of GenoMic Annotation; MVP, Million Veteran Program; MR, Mendelian randomization; MVMR, multivariable Mendelian randomization; IVW, inverse-variance weighted; CRF, corneal resistance factor; CCT, central corneal thickness; CRF⊥CCT, genetically CCT-residualized CRF; SNP, single-nucleotide polymorphism; OR, odds ratio; CI, confidence interval; R/L, right/left eye.

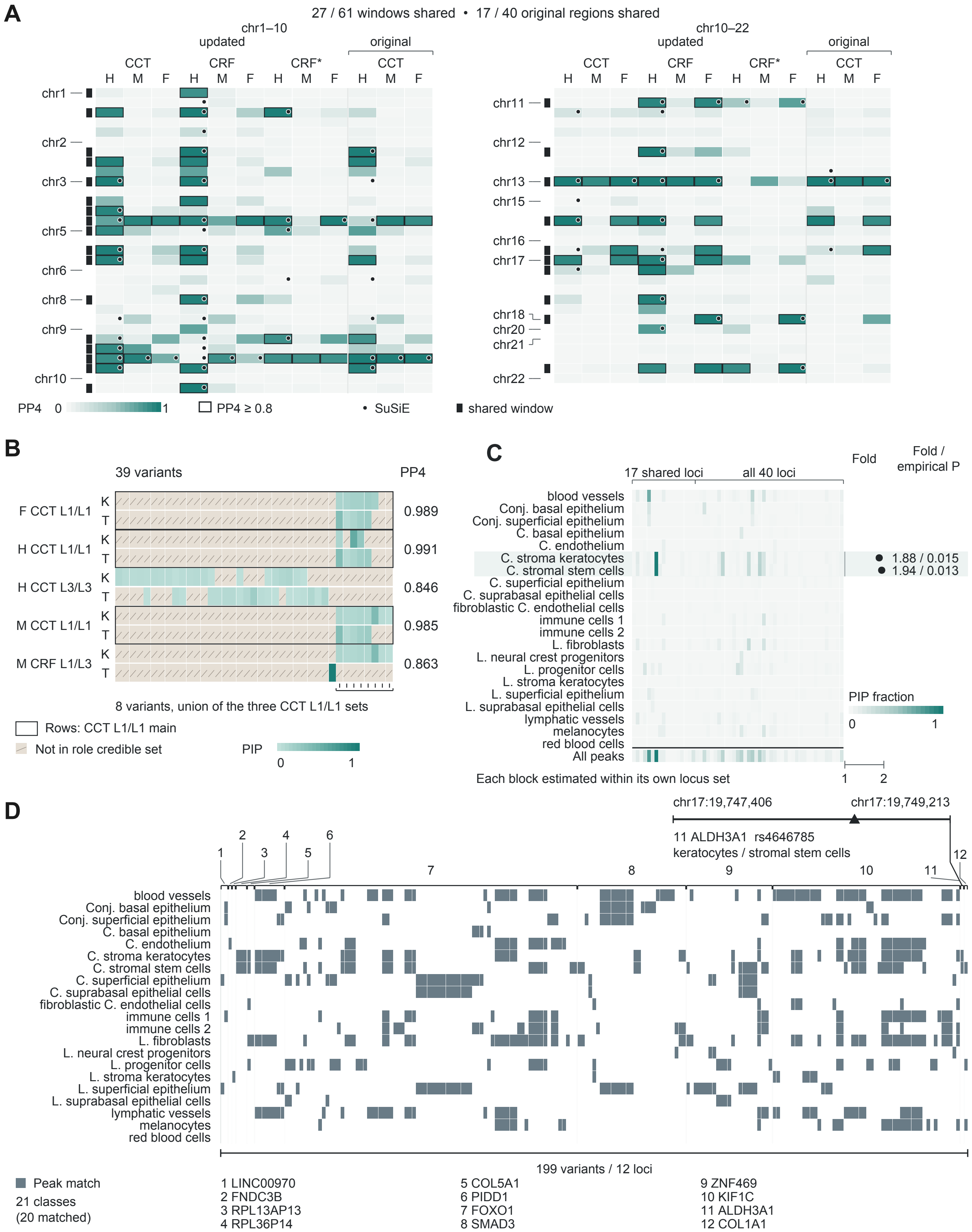

A
27 / 61 windows shared • 17 / 40 original regions shared
chr1–10
updated
original
CCT
CRF
CRF*
H M F
chr1
chr2
chr3
chr5
chr6
chr8
chr9
chr10
chr10–22
chr11
chr12
chr13
chr15
chr16
chr17
chr18
chr20
chr21
chr22
PP4 0 1
PP4 ≥ 0.8
SuSiE
shared window
B
39 variants
PP4
F CCT L1/L1
H CCT L1/L1
H CCT L3/L3
M CCT L1/L1
M CRF L1/L3
K
T
0.989
0.991
0.846
0.985
0.863
8 variants, union of the three CCT L1/L1 sets
Rows: CCT L1/L1 main
Not in role credible set
PIP
0 1
C
17 shared loci
all 40 loci
Fold
Fold / empirical P
blood vessels
Conj. basal epithelium
Conj. superficial epithelium
C. basal epithelium
C. endothelium
C. stroma keratocytes
C. stromal stem cells
C. superficial epithelium
C. suprabasal epithelial cells
fibroblastic C. endothelial cells
immune cells 1
immune cells 2
L. fibroblasts
L. neural crest progenitors
L. progenitor cells
L. stroma keratocytes
L. superficial epithelium
L. suprabasal epithelial cells
lymphatic vessels
melanocytes
red blood cells
All peaks
1.88 / 0.015
1.94 / 0.013
PIP fraction
0 1
Each block estimated within its own locus set
1 2
D
chr17:19,747,406
chr17:19,749,213
11 ALDH3A1 rs4646785
keratocytes / stromal stem cells
1 2 3 4 5 6 7 8 9 10 11 12
blood vessels
Conj. basal epithelium
Conj. superficial epithelium
C. basal epithelium
C. endothelium
C. stroma keratocytes
C. stromal stem cells
C. superficial epithelium
C. suprabasal epithelial cells
fibroblastic C. endothelial cells
immune cells 1
immune cells 2
L. fibroblasts
L. neural crest progenitors
L. progenitor cells
L. stroma keratocytes
L. superficial epithelium
L. suprabasal epithelial cells
lymphatic vessels
melanocytes
red blood cells
199 variants / 12 loci
Peak match
21 classes
(20 matched)
1 LINC00970
2 FNDC3B
3 RPL13AP13
4 RPL36P14
5 COL5A1
6 PIDD1
7 FOXO1
8 SMAD3
9 ZNF469
10 KIF1C
11 ALDH3A1
12 COL1A1

**Supplementary Figure S5. Structural sharing extends from risk windows to credible components and corneal accessible elements.**

**(A)** PP4 for 61 candidate windows across three disease sources and four trait/version channels: updated CCT (GCST90000654; Choquet 2020), CRF, CRF⊥CCT, and original CCT (GCST006366). Genome-ordered windows are split into 31 and 30 rows, with chromosome 10 continuing across the split. Cell color shows PP4 from 0 to 1; borders indicate PP4 ≥ 0.8, corner circles indicate SuSiE, and the black track identifies sharing in any of the three primary channels, excluding the original-CCT comparison channel. These channels yield 27 of 61 shared windows, including 17 of the original 40. Windows can overlap. Source-specific posteriors and methods are in Supplementary Table S5.

**(B)** COL5A1 credible-set memberships for five shared component pairs, ordered by GRCh38 position. The 39 distinct variants contribute 81 component–role–variant PIP records. K and T identify KC and structural-trait credible-set roles. Teal cells show original PIP; hatched gray cells denote variants outside that role's credible set. Outlines identify the three primary CCT L1/L1 pairs; their eight-variant union is bracketed. The Hardcastle, MVP, and FinnGen primary-component unions contain 5, 8, and 6 variants, respectively. Displayed PP4 values are FinnGen CCT L1/L1, 0.989; Hardcastle CCT L1/L1, 0.991; Hardcastle CCT L3/L3, 0.846; MVP CCT L1/L1, 0.985; and MVP CRF L1/L3, 0.863. L1 and L3 retain the fitted component identifiers. PP4 is the component-pair posterior; cell color represents variant PIP.

**(C)** KC credible-set PIP fractions overlapping accessible peaks in 21 author-defined cell classes and a separate all-deposited-peaks aggregate. The shared-region field contains 17 × 22 values and the background field 40 × 22 values. The 17 regions are included in the 40, but each field retains its own signal inputs and aggregation: shared pairs are grouped together, whereas the background is averaged by source groups. Thus the same region–class key can have different values in the two fields, and columns have independent region ordering. The two highlighted stromal classes are selected displays: keratocytes have 1.88-fold enrichment (empirical $P$ = 0.015), and stromal stem cells 1.94-fold enrichment ($P$ = 0.013). All class results, including other enriched classes, are in Supplementary Table S5, block D. Empirical $P$ values use 1,000 draws of 17 regions from the original 40; stored null intervals describe the permutation distribution.

**(D)** Recorded accessible-peak matches across 199 locus–SNP columns from 12 loci and 21 cell classes. Slate cells identify at least one peak match: 683 occupied positions derive from 1,165 source rows, 373 signal–SNP identities, and 96 peaks. Twenty classes have matches; the red-blood-cell row has no recorded match. Empty cells denote absence of a recorded match. Block numbers identify the locus names below the matrix. The ALDH3A1 inset shows rs4646785 at GRCh38 chr17:19,748,589 (PIP, 0.955), within peak chr17:19,747,406–19,749,213, matched in corneal stromal keratocytes and stromal stem cells. The regional COL5A1 matches are distinct from the eight primary-component variants in B. Supplementary Table S5 summarizes regulatory context.

H/M/F, Hardcastle/MVP European/FinnGen R13; MVP, Million Veteran Program; KC, keratoconus; CCT, central corneal thickness; CRF, corneal resistance factor; CRF* or CRF⊥CCT, genetically CCT-residualized CRF; PP4, posterior probability that two traits share a causal variant; PIP, posterior inclusion probability; SuSiE, sum of single effects; SNP, single-nucleotide polymorphism; GWAS, genome-wide association study; ATAC, assay for transposase-accessible chromatin; GRCh38, Genome Reference Consortium human genome build 38; C., corneal; Conj., conjunctival; L., limbal.

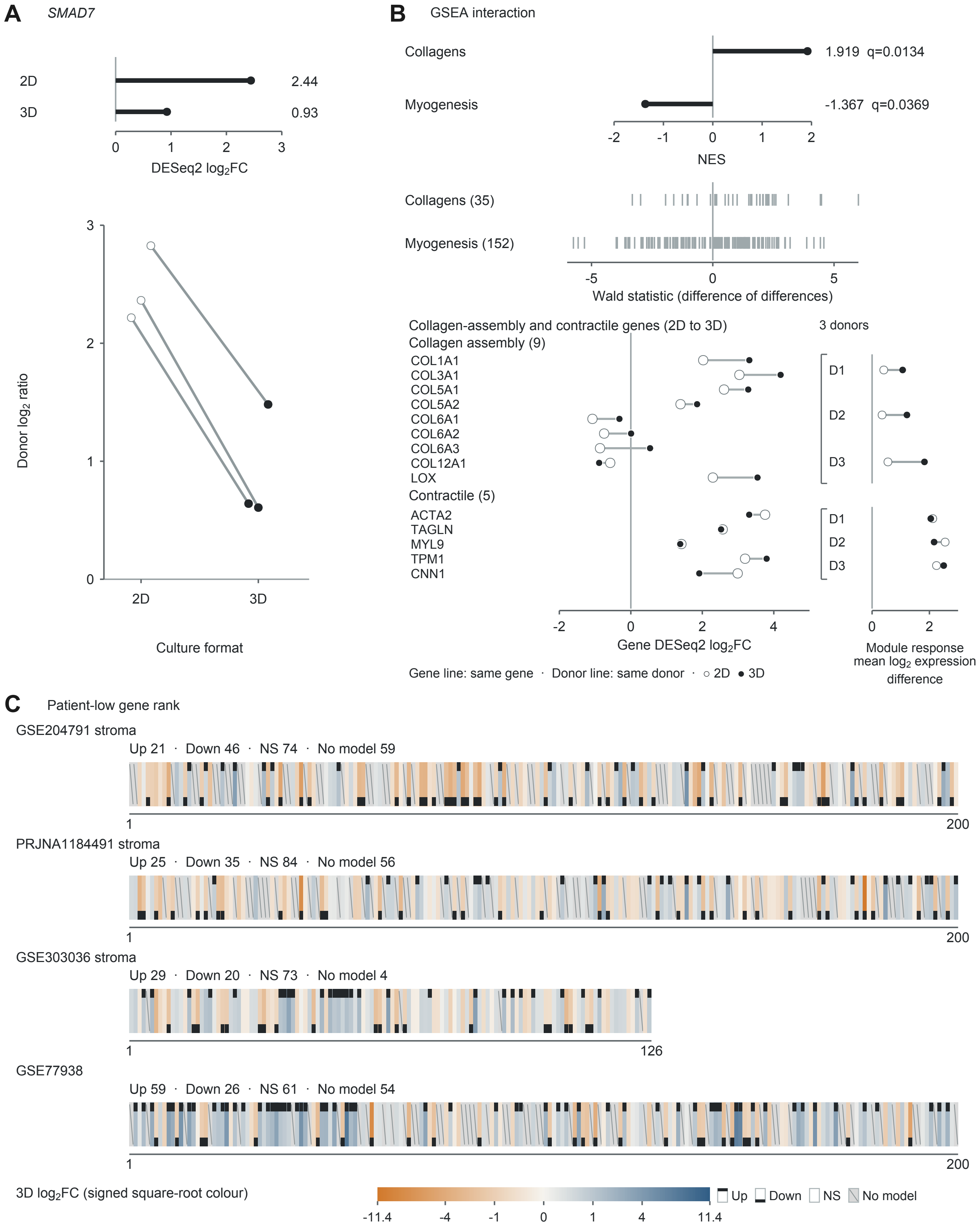
A
SMAD7
2D
3D
2.44
0.93
0
1
2
3
DESeq2 log2FC
Donor log2 ratio
Culture format
B
GSEA interaction
Collagens
Myogenesis
1.919 q=0.0134
-1.367 q=0.0369
NES
Collagens (35)
Myogenesis (152)
-5
0
5
Wald statistic (difference of differences)
Collagen-assembly and contractile genes (2D to 3D)
3 donors
Collagen assembly (9)
COL1A1
COL3A1
COL5A1
COL5A2
COL6A1
COL6A2
COL6A3
COL12A1
LOX
Contractile (5)
ACTA2
TAGLN
MYL9
TPM1
CNN1
D1
D2
D3
-2
4
Gene DESeq2 log2FC
Module response mean log2 expression difference
Gene line: same gene · Donor line: same donor · ○ 2D ● 3D
C
Patient-low gene rank
GSE204791 stroma
Up 21 · Down 46 · NS 74 · No model 59
200
PRJNA1184491 stroma
Up 25 · Down 35 · NS 84 · No model 56
GSE303036 stroma
Up 29 · Down 20 · NS 73 · No model 4
126
GSE77938
Up 59 · Down 26 · NS 61 · No model 54
3D log2FC (signed square-root colour)
-11.4
-4
-1
1
11.4
Up
Down
NS
No model

**Supplementary Figure S6. KC-derived cells retain feedback and matrix induction across culture formats without uniform reversal of tissue-low genes.**
